# ThermiQuant™ MegaScan: High-throughput isothermal reactor with quantitative colorimetric readout for paper-based nucleic acid amplification tests

**DOI:** 10.64898/2026.01.09.696240

**Authors:** Bibek Raut, Gopal Palla, Virendra Kumar, Andrew Fleck, Bilal Ahmed, Josiah Levi Davidson, Ryan F. Relich, Jon P. Schoonmaker, J. Alex Pasternak, Mohit S. Verma

## Abstract

Isothermal nucleic acid amplification tests (NAATs), such as loop-mediated isothermal amplification (LAMP) implemented on microfluidic paper-based analytical devices (µPADs), enable inexpensive and rapid (≤60 min) colorimetric molecular diagnostics; however, no existing instrument supports high-throughput (>100 reactions) quantitative analysis of colorimetric isothermal assays on paper substrates under controlled laboratory conditions. To address this gap, we developed ThermiQuant™ MegaScan, a scanner- and water-bath–based platform that accommodates a 160-reaction µPAD cartridge, maintains uniform incubation at 65 ± 0.5 °C, and enables real-time imaging every 30 s. We also developed accompanying software, Amplimetrics™, for automated µPAD detection and kinetic colorimetric analysis. Using paper-based colorimetric LAMP targeting the SARS-CoV-2 *orf7ab* region, the assay achieved a limit of detection at 95% probability (LOD95) of 34 copies per reaction (5 copies/µL) and a limit of quantification (LOQ) of 250 copies per reaction (33 copies/µL) using purified synthetic DNA targets, and achieved 72% sensitivity and 100% specificity relative to digital PCR (dPCR) for diluted human nasopharyngeal (NP) swab virus samples. We further evaluated the effects of viral and universal transport media (VTM/UTM) on assay performance and found that linear calibration derived from synthetic targets do not reliably translate to clinical samples in these media. Together, these results establish ThermiQuant™ MegaScan as a high-throughput laboratory research platform for standardized evaluation, optimization, and benchmarking of paper-based colorimetric nucleic acid amplification assays.

## Introduction

While liquid-based nucleic acid amplification tests (NAATs) benefit from commercial thermocyclers integrated with optical sensors that support high-throughput testing in standard 96-384 well-plate formats, film pouches, or onboard low-to-high-throughput automated platforms, no equivalent instruments exist for performing colorimetric isothermal NAAT reactions on microfluidic paper-based analytical devices (µPADs). To address this gap, we developed ThermiQuant™ MegaScan, a flatbed scanner-integrated water-bath reactor that accommodates up to 160 paper reactions in a custom acrylic cartridge with a standard 96-well plate footprint (12.1 cm × 85.0 cm).

In recent years, isothermal NAAT methods have gained traction as alternatives to polymerase chain reaction (PCR) for point-of-need (PON) diagnostics. Although both approaches require specialized instrumentation to maintain reaction conditions, isothermal NAATs operate at a constant temperature, simplifying the design requirements compared to systems that rely on thermal cycling^1–3^. Among NAATs, loop-mediated isothermal amplification (LAMP) is widely adopted, typically performed at 60–65 °C^4–8^. When paired with pH-sensitive dyes such as phenol red in an unbuffered reaction, LAMP produces a red-to-yellow color shift visible to the naked eye, caused by proton accumulation as amplification progresses and the reaction pH decreases^9–11^. Incorporating these colorimetric formulations into µPADs makes them particularly attractive: paper supports passive wicking of fluids, enables storage of dried reagents, accommodates multiplexed layouts, and is inexpensive and disposable^12–19^.

However, amplification on µPADs often generate weak or spatially uneven signals near the limit of detection (LOD) especially in complex minimally-processed samples^20–22^. As a result, visual interpretation of colorimetric outcomes is often unreliable, particularly when signal strength or hue variation is subtle. For individuals with color vision deficiencies, this challenge is even greater, as small shifts in color tone (for example, between red and yellow) may be difficult to discern reliably^23,24^. Image data collected during the COVID-19 pandemic by the UK REACT-2 program, comprising more than 595,000 participant-submitted SARS-CoV-2 antibody lateral flow assay images from 2020-2021, showed that weak positives were the main source of disagreement between user reads and experts, primarily due to low-intensity bands^25^. Similar difficulties have been documented in CRISPR-Cas13-based nucleic acid lateral flow assays, where low-intensity test bands were a major source of false calls^26^. Consistent with these reports, our pilot study with µPAD LAMP sensors found misclassification rates of up to 25%, largely due to ambiguous mixed yellow-red outcomes that could not be reliably judged even with a reference chart^13^.

To overcome the subjectivity of visual color interpretation, there remains a need for digital readouts that can reliably quantify colorimetric signals using cameras or color sensors integrated into heating modules. Representative devices reported over the past five years for colorimetric and fluorometric monitoring of LAMP reactions in both µPAD and liquid formats are summarized in Table 1. Commercial thermocyclers, while capable of high throughput workflow, employ small-aperture (< 2 mm diameter) optics that sequentially scan individual wells or rows of wells. This approach is well suited to homogeneous liquid reactions, where each well produces a uniform optical signal and a single-point measurement is sufficient. However, µPADs generate spatially heterogeneous color patterns across the reaction zone, making such point-based optics poorly suited for accurate full field of view analysis. Consequently, existing thermocyclers cannot provide the wide-field imaging necessary for µPAD assays. Beyond commercial systems, reported instruments for µPADs are primarily portable point-of-need devices emphasizing compact size (<20 cm) and low weight (<1 kg), which limits throughput to fewer than a dozen reactions per run. Yet systematic assay optimization in the laboratory often requires high-throughput workflows (>100 reactions), for which no commercial or published systems currently exist.

**Table 1:** Comparison of liquid- and paper-based isothermal nucleic acid amplification test (NAAT) instruments with optical (color/fluorescence) tracking capability. Acronyms: PCB, printed circuit board; PTC, positive temperature coefficient; µPAD, microfluidic paper-based analytical device; LAMP, loop-mediated isothermal amplification; CRISPR, clustered regularly interspaced short palindromic repeats.

| Device or commercial product | Reaction format | Heater type & temperature | Imager or sensor type | Throughput |
| --- | --- | --- | --- | --- |
| ThermiQuant™ MegaScan (this work) | μPAD | Water bath, 65 ± 0.5 $^{\circ}$C | Flatbed scanner | 160 μPADs |
| Field Applicable Rapid Microbial LAMP (FARM-LAMP<sup>21</sup>) | μPAD | Water bath, 65 $^{\circ}$C | Mini camera (16 MP) | 12 μPADs |
| Ubiquitous NAAT (UbiNAAT<sup>14</sup>) | μPAD | PCB heater, 65 $^{\circ}$C | Smartphone with fluorescence optics | 8 μPADs |
| LAMP-CRISPR device<sup>28</sup> | μPAD | Resistive thin-film heater, 65 $^{\circ}$C | Fluorescence detectors | 6 μPADs |
| Microfluidics Integrated LED-Photodiode (MILP device<sup>29</sup>) | μPAD | PTC heater, 65 $^{\circ}$C | Photodiodes with optical filters | 3 μPADs |
| Thermocyclers (Eg. Analytik Jena qTOWER<sup>3</sup>, QuantStudio<sup>6</sup> & 7 Pro, CFX Opus 384, Cielo3) | Liquid | Metal heat block, 20 to 100 $^{\circ}$C (± 0.1-0.5 $^{\circ}$C) | Fluorescence detectors | 96 to 384 tubes or wells |
| Smartphone operated LAMP (SMART-LAMP<sup>30</sup>) | Liquid | Polyamide heater, 65 ± 0.9 $^{\circ}$C | TCS3472 color sensor | 8 tubes |
| Quantitative colorimetric LAMP (qcLAMP device<sup>31</sup>) | Liquid | PCB heater, 63 $^{\circ}$C | Mini camera | 8 tubes |
| Integrated Gene box (i-Genbox<sup>32</sup>) | Liquid | Thin-film heater, 65 $^{\circ}$C | Smartphone camera | 7 wells |
| LAMP Assay Reader Instrument (LARI<sup>33</sup>) | Liquid | Aluminum heat block, 69 ± 0.4 $^{\circ}$C | Digital light sensor | 12 tubes |

To address the throughput limitations of existing colorimetric evaluation instruments, we introduce ThermiQuant™ MegaScan (hardware) and Amplimetrics™ (software), which together provide a unified workflow for high-throughput colorimetric NAATs on µPADs. The ThermiQuant™ MegaScan combines precise thermal regulation (“Therm”) with integrated imaging (“i”) for quantitative (“Quant”) analysis across a large scanning field (“MegaScan”) capable of capturing over 100 reaction zones simultaneously. The system achieves uniform isothermal control at 65 ± 0.5 °C using a repurposed consumer-grade circulating water bath coupled to a transparent imaging window that allows distortion-free time-lapse imaging at sub-minute intervals. An exploded view of the instrument is shown in Figure 1A, and the custom cartridge supporting 160 µPAD reaction zones is shown in Figure 1B. Complementing the hardware, Amplimetrics™ software automates the analysis of scanner-captured images acquired during the 60-minute LAMP reaction, generating hue-versus-time curves (Figure 1C, rightmost panel) and computing a quantification time (Tq) analogous to the quantification cycle (Cq) in quantitative PCR (qPCR)^27^.

**Figure 1.**
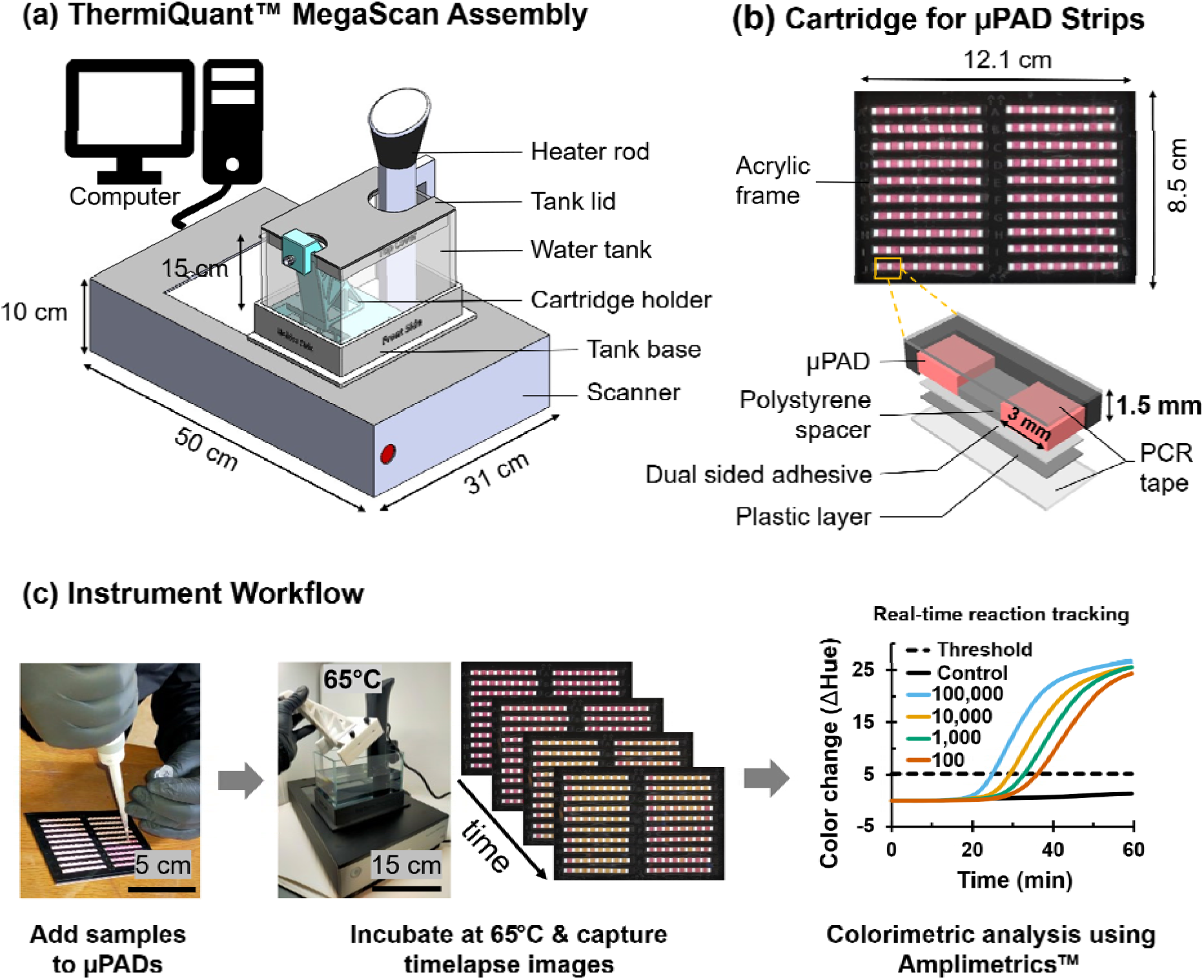
Design and workflow of the ThermiQuant™ MegaScan platform. (A) Schematic of the instrument with labeled components. (B) Cartridge containing 160 independent microfluidic paper-based analytical device (µPAD) reaction zones; inset illustrates the layer-by-layer configuration of the µPAD reaction strip. (C) Typical workflow, including sample loading onto µPADs, incubation at 65 °C in an isothermal water bath, time-lapse imaging with a flatbed scanner positioned underneath, and subsequent analysis of colorimetric changes over time using the Amplimetrics™ software.

We tested the system using synthetic targets and diluted human nasopharyngeal swabs containing SARS-CoV-2 by targeting the *orf7ab* region of the virus. The paper-based LAMP assay achieved an LOD at 95% detection probability (LOD95) of 34 copies per reaction (5 copies/µL) and a limit of quantification (LOQ) of 250 copies per reaction (33 copies/µL), achieving a strong linear calibration using synthetic targets (R^2^ = 0.97). In the case of 5% diluted clinical SARS-CoV-2 virus samples, the assay achieved 72% sensitivity and 100% specificity relative to digital PCR (dPCR). Together, these tools establish ThermiQuant™ MegaScan as a high-throughput platform for quantitative µPAD assay development, enabling rigorous kinetics-based optimization of the point-of-need µPAD for application in the downstream One Health applications in clinics, on farms, and outdoors.

## Results

### ThermiQuant™ MegaScan design enables high-throughput analysis

To overcome the limited throughput of existing paper-based color NAAT systems (maximum of 12 µPADs per run), we developed the ThermiQuant™ MegaScan platform, which integrates a flatbed charged coupled device (CCD) scanner with a precision sous-vide water bath maintained at 65°C ± 0.5 °C to enable real-time colorimetric tracking of LAMP reaction kinetics (Figure 1, Figure S1, and Figure S2). The modular configuration was intentionally constructed using off-the-shelf components to allow straightforward reproducibility and assembly in any research laboratories working on paper-based molecular sensors. The system is intended as a laboratory-based research tool rather than for direct clinical translation. Further validation, hardware and software integration, and regulatory approval would be required for clinical deployment. The accompanying disposable acrylic cartridge with customizable design accommodates anywhere from 1 to 160 µPADs. Figure 1B shows an example of 160 µPADs in two columns of 10 rows, each row containing eight µPADs each of size 3 mm × 3 mm and Figure S3 shows step by step µPAD fabrication process. This throughput nearly doubles that of a conventional 96-well liquid platform and far exceeds most reported paper-based systems, which typically support ≤12 reactions^21,28,29,34–36^ (Table 1). Although the scanner’s maximum imaging area (21.6 cm × 29.7 cm) could physically accommodate two water tanks side by side, raising the potential throughput to 320 µPADs, the scanner instrument frame is not engineered to support the ∼10 kg load of dual water tanks, so we limited operation to a single tank in this study. Figure S1 provides an exploded view of the assembly, Figure S2 drawings of 3D printed parts, Figure S4A illustrates alternative cartridge formats designed for smaller batch experiments besides the high-throughput 160 µPAD reactions, and Figures S4B-C shows the three µPAD strip design. Table S1 shows key scanner settings. Note 1.1 describes instrument assembly and operation manual; Note 1.2 scanner timelapse guide; Note 1.3 Amplimetrics™ installation guide; and Note 1.4 explanation of the Amplimetrics™ software. For LAMP demonstration, we used the 2 and 3 µPAD strip version of the cartridge.

The layered design of the µPAD strips^7,8,21^, shown in the inset of Figure 1B and Figure S3, minimized crosstalk by interleaving µPAD with plastic spacers backed with double-sided adhesive. Each µPAD consistently retained 7.5 µL of liquid without leakage owing to strong capillary retention, and spillover was observed only when volumes exceeded 9 µL. In addition, sealing the acrylic cartridge on both faces with PCR sealing tape created a watertight barrier, and no cross-contamination or leakage was detected during operation. Together, these findings confirm that the µPAD-cartridge assembly provided robust fluid handling and reliable compartmentalization of individual reactions.

The µPAD strips were fabricated using a semi-automated workflow utilizing a leather cutter and manual assembly (Figure S3), enabling routine production of >1,000 2-8 µPAD strip assemblies per day by a single operator. To scale up manufacturing in the future, the process can be further adapted to roll-to-roll processing and injection molding for potential production of tens to hundreds of thousands of devices per day.

The hardware cost of the system was USD 1,573.30 (Table S2), and each cartridge supported up to 160 reactions at a cost of USD 122.89 (USD 0.77 per reaction, Table S3). The operational workflow, summarized in Figure 1C, required 10 minutes for sample pipetting (using a single-channel pipette) to µPADs and cartridge sealing, 2 minutes for scanner setup, and 3 minutes for image processing and data analysis (using Amplimetrics™ software) after experimentation was complete. Thus, preparation for a full 160-reaction run, along with data analysis, could be completed in under 15 minutes of hands-on time (although the LAMP reaction itself requires 60 minutes of hands-off time). These results demonstrate that MegaScan is a rapid, reliable, and cost-effective platform for high-throughput NAATs on µPADs in laboratory settings.

### Cartridge thermal stabilization is rapid and uniform

The MegaScan water bath enabled rapid and uniform heating of the cartridge during amplification. Depending on the water volume (3-3.5 L), it required 10-15 minutes to reach the setpoint of 65 °C from room temperature. Next, when a room-temperature cartridge was inserted into the preheated 65 °C water bath, thermocouple measurements at four diagonal positions (Figure 2A) showed that the µPADs reached 65 °C within 2 minutes (Figure 2B) and that spatial temperature variation across the cartridge remained within ±0.5 °C over the 60-minute duration (Figure 2C, data shown for the first 2 minutes). Efficient temperature equilibration was facilitated by the 1.5 mm thin acrylic cartridge, which allowed rapid heat transfer into the µPADs as well as excellent heat distribution through the heater water circulator. In addition, the use of a commercial immersion (*sous vide*) heater added built-in safety, and durability features absent in many custom systems: it automatically shut off when the water level fell outside the operating range and is IPX-7 rated for water resistance, reducing risks of overheating or device failure during extended use.

**Figure 2.**
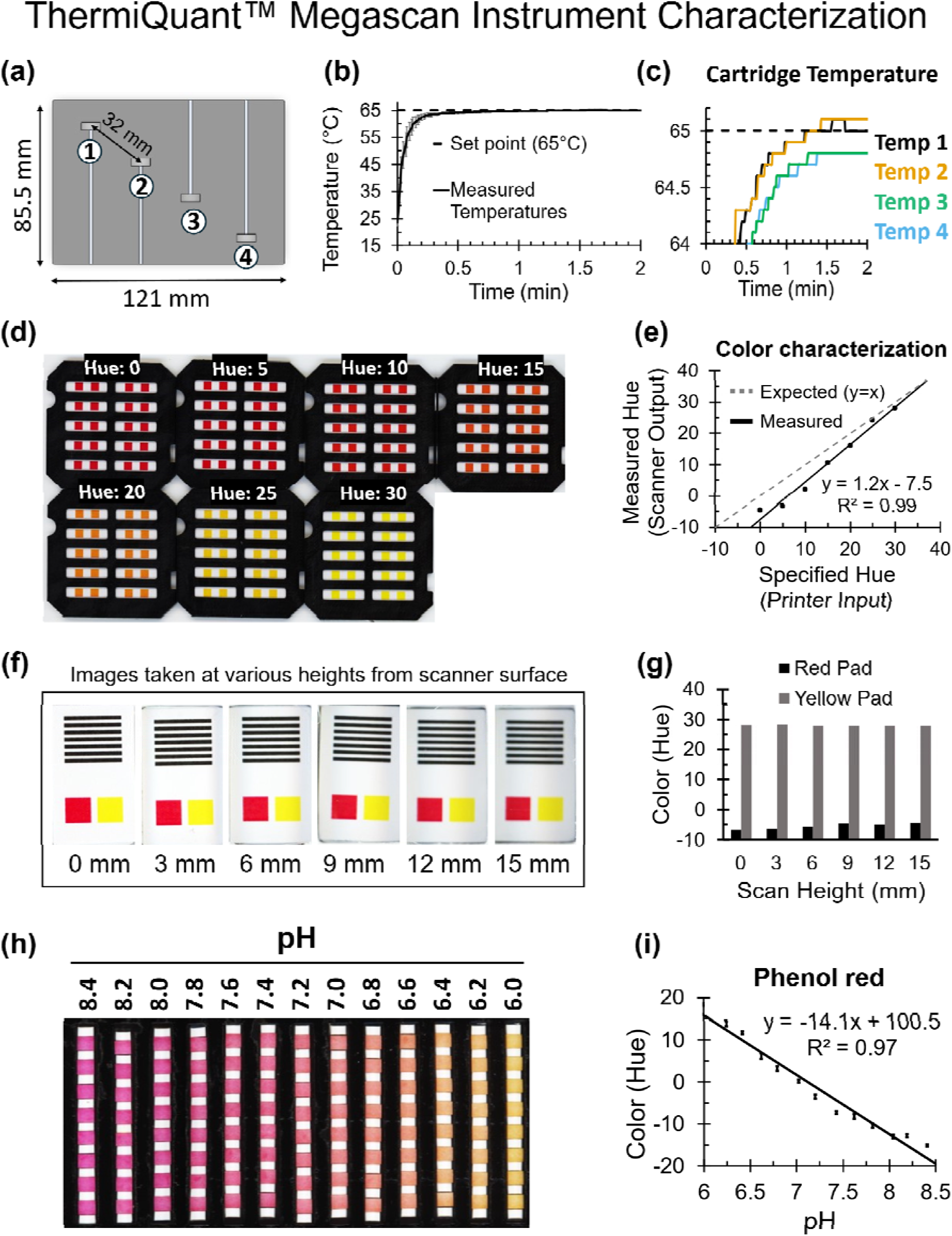
ThermiQuant™ MegaScan instrument characterization. (A) Schematic showing thermocouple placement at four diagonal positions (T1–T4) across the cartridge with 32 mm diagonal spacing. (B) Average cartridge temperature (± SD) when immersed in a 65 °C water bath. (C) Individual probe measurements showing temperature stabilization within 2 minutes and maximum variation within ±0.5°C. (D) Array of color pads (5 × 4 grid) housed in a 3D-printed holder. Color pads were designed in Adobe Illustrator with specified hue values, printed on a standard office color printer, and scanned with a flatbed scanner. (E) Comparison of measured color hue values (± SD) from the scanned pads with the target hues specified during design showing strong correlation but with an offset to the specified hue. (F) Calibrator consisting of two-color pads (red and yellow) representing positive and negative colorimetric reactions, along with parallel printed lines (0.5 mm thickness, 0.5 mm spacing). The printed sheet was positioned at varying heights (0-15 mm) above the scanner surface. (G) Depth-of-field evaluation of color pads positioned at different heights in Figure F; hue values extracted from red and yellow pads showed minimal change with height. (H) Hue-pH relationship of phenol red chemistry on µPADs. Strips pre-soaked in color LAMP master mix (adjusted to pH 6.0–8.4), dried in air for 2 hours and rehydrated with water exhibited a visible red-to-yellow transition. (I) Linear regression of hue values (± SD) measured from µPADs across the pH range, demonstrating strong correlation (R^2^ = 0.97).

### Scanner color detection accuracy is reproducible despite systematic offset

The CCD flatbed scanner provided reproducible hue measurements across the red-to-yellow transition characteristic of phenol red in µPAD LAMP reactions. To assess color accuracy, a printed hue gradient spanning 0–30° in the HSV color space was scanned within a 3D-printed mount (Figure 2D). Hue values were extracted after RGB-to-HSV conversion using the OpenCV color model implemented in Amplimetrics™ software. As described in methods section (“CCD scanner color and scan height characterization”), hue values near the red boundary were remapped to generate a continuous −20° to 30° axis (equation 1 & 2). This correction eliminated discontinuities that otherwise caused averaging artifacts near 0°/180° and enabled linear hue progression consistent with the expected red-to-yellow color shift (Figure S6). The scanner output showed a systematic offset relative to the nominal printed hue color patches (Figure 2E), which likely arises from both printer inaccuracy in reproducing digital colors and scanner spectral response offsets. Although absolute values were shifted, replicate measurements were highly reproducible (<1 hue unit variance) with a strong correlation to designed hues (R^2^ = 0.99). Because amplification kinetics depends on the relative change in hue rather than absolute values, we used relative hue (Δhue) as the primary metric for colorimetric LAMP analysis and to differentiate between positive and negatives. Here, the Δhue value was defined as the hue at a given time point minus the initial hue at 0 min, While advanced calibration algorithms can reduce scanner and printer offsets^37–39^, this adjustment was not pursued here because the key parameter of interest was the relative color change during amplification rather than their absolute value.

### Scanner depth of field is sufficient for cartridge imaging

The CCD scanner maintained stable hue detection across a range of imaging heights, confirming adequate depth of field for cartridge operation. Because the cartridge sits 7.5 mm above the scanner glass during use (5 mm glass thickness, 1.5 mm gap between cartridge and tank base, and 1 mm tank base support thickness), we evaluated color stability by elevating printed red and yellow pads, along with 0.5 mm wide black lines spaced at 0.5 mm to assess focus, in 3 mm increments up to 15 mm (Figure 2F–G). As expected, the black line features became progressively blurred with increasing height (Figure 2F), indicating reduced spatial resolution. In contrast, the measured hue values for red and yellow across all tested distance pads remained nearly constant, with only a 6.8% net hue change between scan heights of 0 mm and 15 mm (Figure 2G). This corresponds to 0.5% hue variation when considering 0.5 mm maximum height variation when placing the holder in the tank. These results demonstrate that the CCD scanner provides sufficient depth of field for µPAD imaging, ensuring robust colorimetric readout even when the cartridge is positioned several millimeters above the scanner surface.

### Hue correlates linearly with pH in phenol red chemistry

Hue values correlated linearly with pH across the phenol red transition range, validating hue as a sensitive optical metric for amplification chemistry. µPADs were pre-soaked in LAMP master mix adjusted to pH values from 6.0 to 8.4, dried, and rehydrated with water before imaging. As expected, the µPADs shifted progressively from red at alkaline pH to yellow at acidic pH (Figure 2H). Image analysis of the hue channel showed a strong linear relationship with pH (R^2^ = 0.97), with a slope of –14.1 hue units per pH unit (Figure 2I). The consistency of the trend confirmed that hue provides a reliable, quantitative proxy for proton release during amplification.

### Amplimetrics™ with YOLOv8 achieves robust µPAD detection and efficient data handling

With the aim of reducing user burden and simplifying image-based analysis, we developed Amplimetrics™, a Python-based software package that integrates automated image processing with downstream data analysis. Figure 3A provides an algorithmic overview of the workflow, while SI Notes 1.3-1.4 include a detailed user guide and explanation of the algorithm. The software is divided into two modules. In the first module, scanner timelapse images are processed by automatically detecting µPADs, segmenting individual pads, and extracting hue-time data. In the second module, the resulting hue–time datasets are baseline-corrected, smoothed, and analyzed to classify reactions using user-defined positivity thresholds, followed by calculation of Tq based on the second-derivative method (see SI Notes 1.4 and Figure S5). In this derivative-based approach, the Δhue–time curve is first processed by taking its derivative with respect to time, then fitted with a Voigt function. Next, the derivative of Voigt function is computed and Tq is defined as the time at which the peak of the second derivative appears. Therefore, Tq corresponds to the onset of the exponential amplification phase and represents the time of maximum reaction acceleration (Figure S5). This modular design enables end-to-end analysis within minutes of experiment run completion, minimizing manual data interpretation while ensuring reproducibility. Moreover, a YOLOv8n object segmentation model trained on 240 manually annotated images and tested on 60 images with more than 1,300 µPADs achieved 100% object detection accuracy. Figure 3B illustrates a representative example of automatic µPAD labeling in a 96-well–style alphanumeric format and Figure 3C shows the confusion matrix. In our experience, a fixed-threshold or geometry-based indexing approach (data not shown) resulted in occasional failures in ROI detection. In contrast, the trained YOLOv8n model achieved 100% instance segmentation accuracy (Figure 3C).

**Figure 3.**
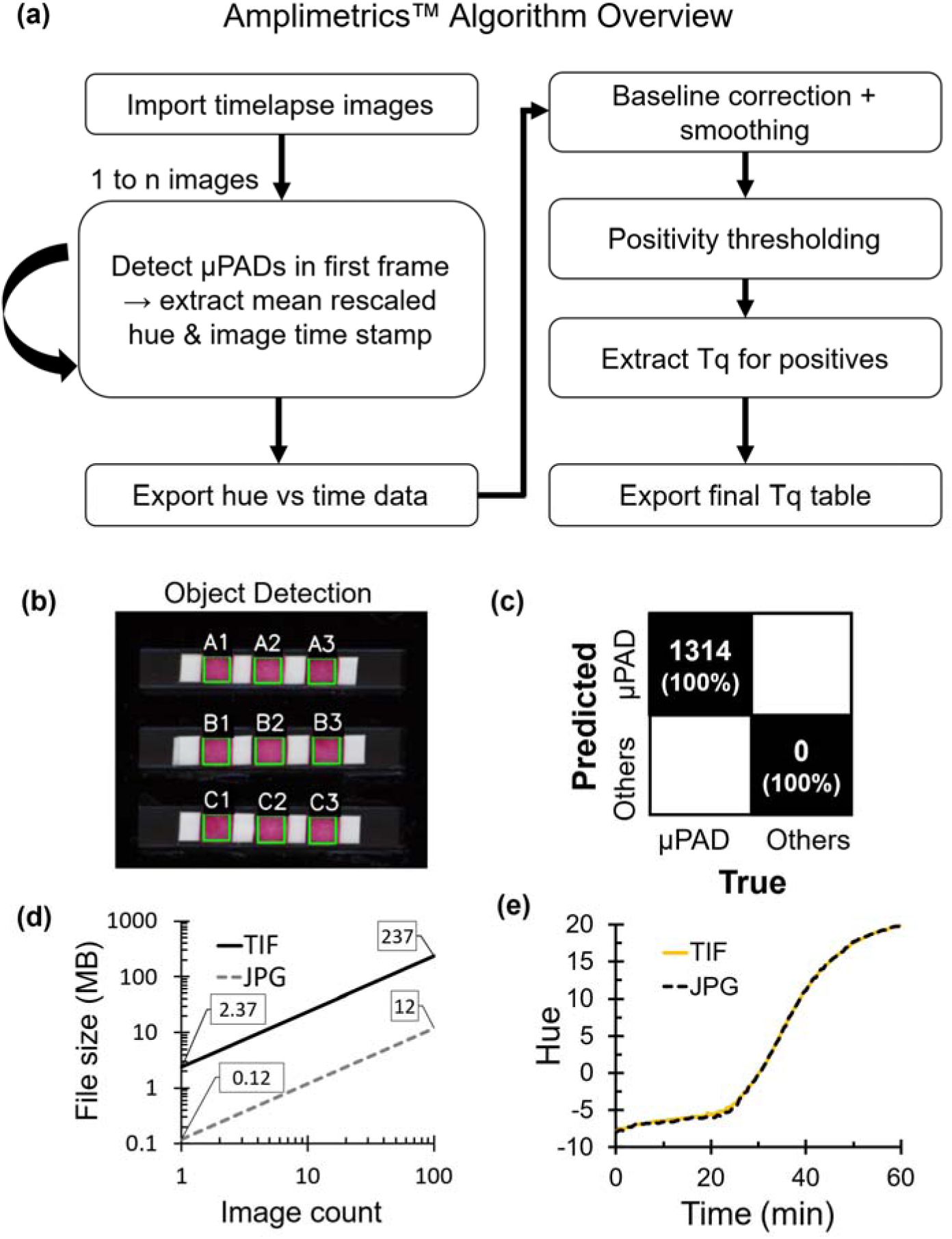
Amplimetrics™ software characterization. (A) Overview of the software workflow, integrating automated image processing with data analysis. (B) Automatic µPAD detection using a YOLOv8-n model trained on 240 images and tested on 60 images. (C) The model generalized to an independent dataset with 100% detection accuracy. (D) For a 100-image timelapse sequence, saving images as JPG with 90% compression reduced total file size by more than 10-fold compared to TIF, demonstrating substantial storage efficiency. (E) Hue-time curves extracted from JPG and TIF images for a typical positive amplification reaction were identical.

To address the large data volumes (several hundred megabytes (MB)) generated by timelapse imaging, we compared file formats supported by the VueScan platform that was used to capture timelapse images. A 100-image sequence of the cartridge (single image: 800 × 1988 pixels at 600 dpi, corresponding to ∼33.9 mm × 84.2 mm which was half-version of the full 160-µPAD high-throughput cartridge) stored as JPG with 90% compression occupied only 12 MB in total (∼121 KB per image), compared with 237 MB for TIF (∼2402 KB per image), representing a >20-fold reduction in digital storage requirements (Figure 3D). For the full 160-µPAD cartridge, the file size would be expected to approximately double. Importantly, hue-time trajectories extracted from compressed JPGs and uncompressed TIFs for a typical positive LAMP reaction were indistinguishable, indicating that lossy compression did not compromise analytical fidelity in this context (Figure 3E). Overall, Amplimetrics™ enables robust µPAD detection, supports the practical use of JPG to minimize storage footprint, and streamlines analysis through a simple two-module workflow. Complete image and data processing for a typical dataset required less than 3 minutes on a Razer Blade 15 laptop (Razer Inc., USA).

### Automated image analysis improves LAMP color interpretability and eliminates subjective bias near the limit of detection

We demonstrated the function of ThermiQuant™ MegaScan platform in combination with Amplimetrics™ software by performing colorimetric LAMP on µPADs using synthetic DNA fragment from the SARS-CoV-2 *orf7ab* gene (NCBI Reference Sequence: NC_045512.2, Table S4). Although SARS-CoV-2 is an RNA virus, a DNA surrogate was chosen to provide a stable, controlled template free of matrix effects while retaining clinical relevance^40^. We have previously demonstrated one-pot reverse-transcription LAMP reactions, where the RNA template undergoes reverse transcription to cDNA at the initiation of the reaction, after which the LAMP process amplifies the resulting DNA^13,41^. Serial dilutions from 10^6^ copies to 1 copy per reaction (7.5 µL reaction volume) were tested in triplicate, along with no-template controls (NTCs), and the experiment was repeated twice (run1 and run2). At high input concentrations (≥250 copies/reaction, 33 copies/ µL), all technical replicates produced a clear red-to-yellow endpoint transition which can be subjectively evaluated as positive by the naked eye (Figure 4A). In contrast, at lower concentrations near the LOD95 (≤34 copies/reaction or 5 copies/µL), endpoint colors were faint, spatially heterogeneous, and difficult to classify reliably by visual inspection.

**Figure 4.**
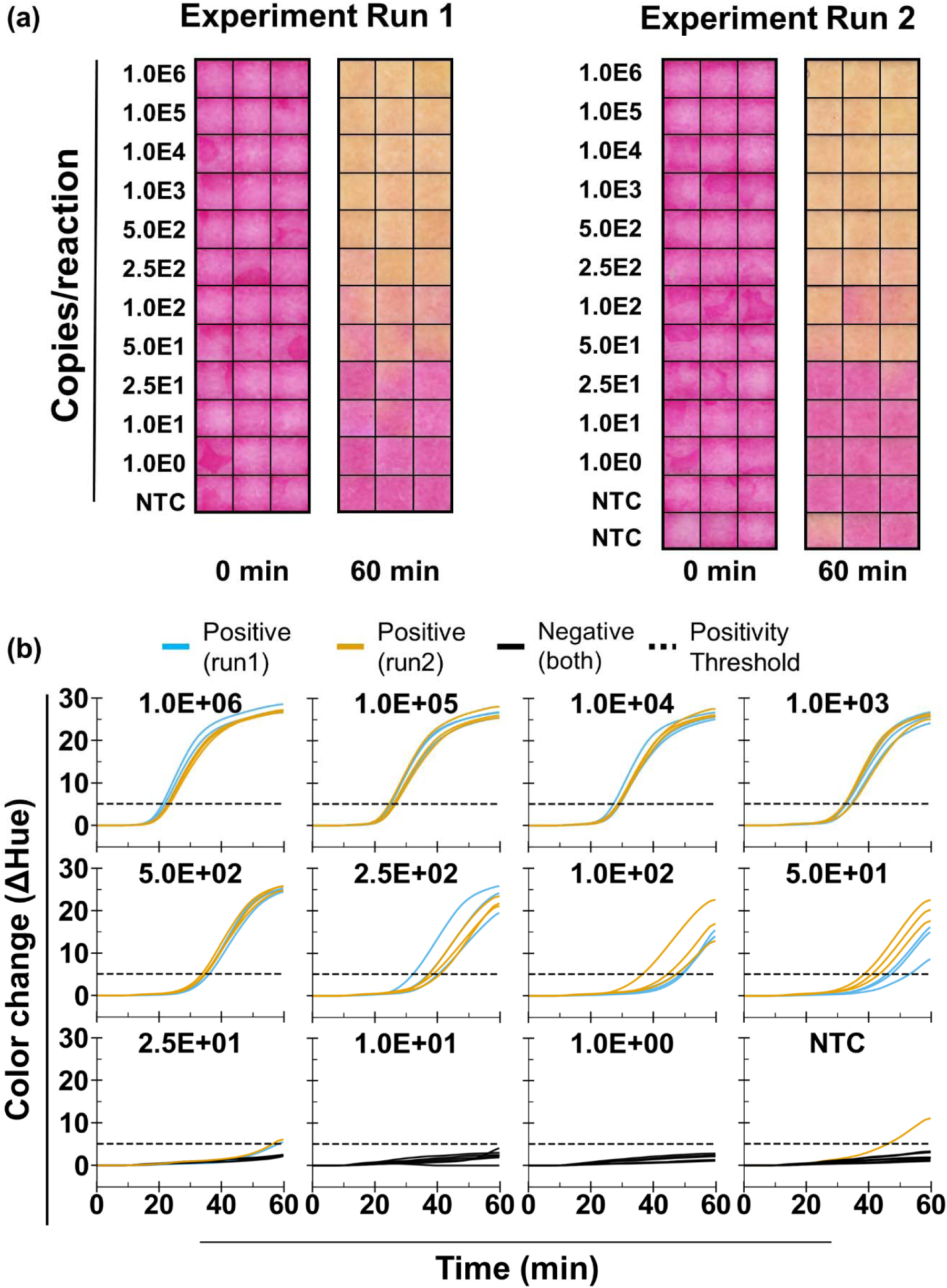
Analytical test of colorimetric LAMP on µPADs. (A) Endpoint images of µPADs loaded with serial dilutions of synthetic SARS-CoV-2 *orf7ab* DNA (1–10^6^ copies/reaction), including no-template controls (NTC), with three technical replicates across two independent experimental runs (second run has six NTCs). Clear visual differentiation is observed between positive samples (yellow) and NTCs (red) at higher concentrations, while mixed coloration appears at or near 50 copies/reaction which is close to the limit of detection at 95% probability (LOD95). (B) Net change in hue channel versus time for run 1 and run 2 across different concentrations of synthetic *orf7ab* DNA. Positive reactions exhibit a logistic sigmoidal pattern, whereas NTCs remain linear. A positivity threshold line was used to distinguish positive from negative outcomes. Images shown in A were post-processed in Microsoft PowerPoint by adjusting brightness (+20%) and contrast (+40%) for improved visual representation; however, all image analyses presented in B were generated from raw image data using custom Python-based software, Amplimetrics™.

Automated analysis with Amplimetrics™ converted scanner time-lapse images into Δhue-time trajectories (Figure 4B). Positive reactions generated characteristic sigmoidal S-shaped curves, whereas NTCs remained linear (Figure 4B). Classification was performed using a threshold defined as the mean NTC signal plus k × SD, where k is an empirically selected parameter to maximize separation between NTC and amplification curves, and SD is the standard deviation. Any outlier or false amplification detected in NTCs was excluded (in our experiments, one nonspecific amplification was observed in a single NTC replicate in run 2, but none in run 1). This NTC-anchored threshold enabled positive/negative classification, including at low template concentrations where endpoint color alone was ambiguous.

We estimated the LOD95 using probit regression of pooled binary amplification outcomes from two independent experiments (Figure S8A; n = 6 replicates per concentration). The model predicted a 95% detection probability at 34 copies per reaction (5 copies per µL).

### Piecewise quantification time (Tq) and log_10_ concentration calibration reveals reproducible quantification only at ≥250 copies per reaction

We next evaluated quantitative precision to determine the LOQ. Based on a coefficient of variation (CV ≤ 10%) criterion applied to replicate Tq values, we determined the LOQ to be 250 copies per reaction (Figure S8B). Linear regression of Tq versus log_10_ concentration using data at or above the LOQ demonstrated strong linearity (R^2^ = 0.97) (Figure S8C).

To examine calibration behavior at and above the LOQ, we fitted a continuous piecewise linear model across two independent experiments (Run1 and Run2) with a fixed breakpoint at the LOQ (250 copies per reaction, Figure 5A). Tq was extracted with Amplimetrics™ as the point of maximum kinetic acceleration, determined from the second derivative of the Δhue-time curve (Figure S5, also see methods). Figure S7 shows the ΔHue-time traces split into high (10^3^-10^6^ copies/reaction) and low (50-10^3^ copies/reaction) concentration segments for Run 1 and Run 2, with NTC controls included. In high copy segment (concentrations ≥ 250 copies per reaction), Tq exhibited strong linear dependence on log_10_ concentration in both runs (run 1: R^2^ = 0.98; run 2: R^2^ = 0.92). In contrast, for low copy segment (concentrations < 250 copies per reaction), linear correlation decreased to R^2^ = 0.68 in run 1 and R^2^ = 0.80 in run 2, indicating greater dispersion in amplification kinetics at lower template inputs.

**Figure 5.**
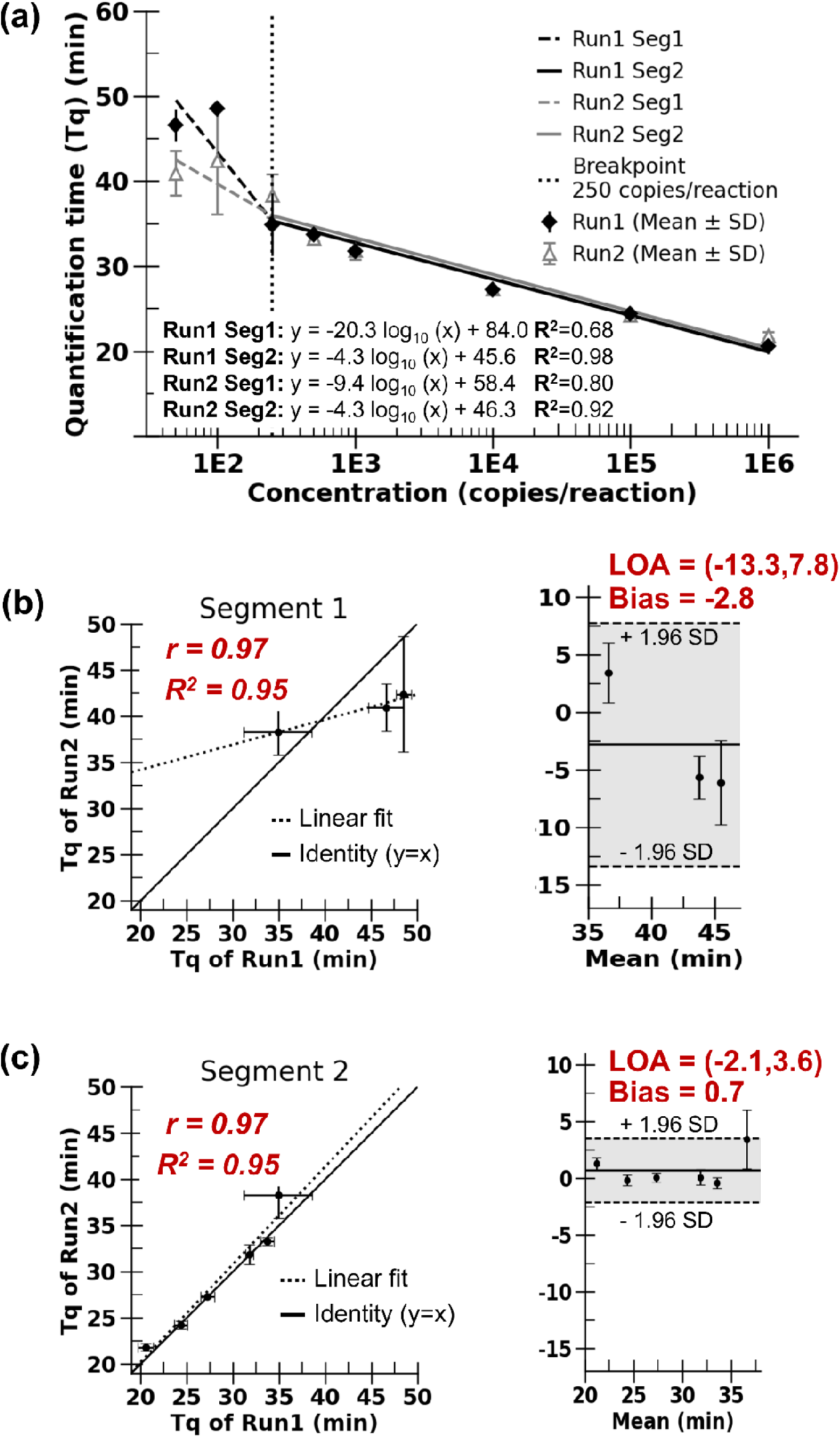
Quantification time (Tq) extraction from hue-time amplification curves. (A) Tq values for each replicate, shown as mean ± SD across two independent experimental runs. Data were fitted with a two-piecewise linear model with breakpoint at limit of quantification (LOQ), dividing the amplification curves into segment 1 and segment 2. (B) Comparison of segment 1 between run 1 and run 2, shown as scatter plots with SD error bars and Bland-Altman analysis. The segment 1 results demonstrate poor agreement, with bias falling outside the instrument’s acceptable error range of ±2 min. (C) Comparison of segment 2 between run 1 and run 2, showing improved correlation and Bland-Altman analysis with bias within maximum allowable instrument error. Results indicate that repeatability is achieved when using segment 2 for quantitative evaluation but not with segment 1.

Bland–Altman analysis further helped explain this divergence (Figure 5B and Figure 5C). At concentrations ≥ 250 copies per reaction (Segment 2), the mean bias between runs was 0.7 min with 95% limits of agreement (LOA) from –2.1 to 3.6 min. Below 250 copies per reaction (segment 1), the bias shifted to –2.8 min with substantially wider LOA (–13.3 to 7.8 min), reflecting increased variability and offset across runs in the lower concentration segment. In the higher concentration segment, the bias was within the ±2 min instrument tolerance, which reflects the scanner cycle time (0.5–1.0 min) combined with cartridge loading and setup variability (up to 1 min), and support reproducibility when considering instrument error alone. While Bland-Altman analysis^42^ is typically used to evaluate agreement between different instruments or methods, in this study we applied it to independent runs on the same instrument to quantify bias and variability across repeated measurements. It is important to note that, beyond instrument-related variability captured in this analysis, reproducibility in practice may also be influenced by factors such as serial dilution error, reagent degradation, and variability in reaction chemistry or sample matrix, which should be considered when interpreting calibration performance. In this study, we limited the scope to instrument and software design and therefore did not evaluate the effects of the sample and reagent variability (except use of VTM and UTM in the clinical sample). However, given the high-throughput capability of the ThermiQuant™ MegaScan system, assessing the influence of other complex sample factors would be an interesting direction for future work.

### Transport media effects necessitate dilution of media for paper-based colorimetric LAMP test

Before evaluating clinical samples, we examined the compatibility of transport media (UTM and VTM) with paper-based colorimetric LAMP (Fig. S9). When we spiked synthetic *orf7ab* targets into undiluted VTM or UTM, we did not observe amplification (Figure S9A), and the initial color of the reaction pads shifted toward orange, indicating pH effects from the transport media. In contrast, targets spiked into nuclease-free water produced the expected sigmoidal hue trajectories.

We next evaluated whether dilution with water could mitigate transport media-associated interference. Diluting VTM to 5% and 10% (v/v) restored amplification (Figure S9B); however, reactions at 10% VTM proceeded more slowly than at 5% and did not reach full signal saturation within 60 min (Figure S9B). Based on these observations, we selected a 5% v/v dilution as a compromise that preserved amplification kinetics while minimizing media effects. We applied this dilution strategy to all subsequent clinical sample tests.

### Test of paper-based RT-LAMP on clinical samples enables robust SARS-CoV-2 detection above limit of detection

We evaluated paper-based RT-LAMP using 24 clinical nasopharyngeal swab SARS-CoV-2 samples that had been previously characterized by dPCR and stored at −80 °C in VTM or UTM following multiple freeze-thaw cycles. Samples were diluted to 5% (v/v) in nuclease-free water prior to testing and analyzed in duplicate, except for two samples analyzed in triplicate, yielding a total of 50 independent paper RT-LAMP measurements. This 20-fold dilution reduced the available target RNA concentration by 20-fold prior to amplification, thereby directly impacting analytical sensitivity under the tested conditions.

Each independent test strip contained two µPADs: a no-primer control (NPC) pad and an RT-LAMP reaction µPAD containing primers. We analyzed time-lapse images using Amplimetrics™ to extract mean hue values over time from each µPAD and subtracted the mean hue of the NPC pad from that of the corresponding RT-LAMP pad at each time point to generate a single background-corrected hue–time trajectory per reaction, thereby removing sample- and substrate-related background signal. Figure 6A shows cumulative hue change–time curves for all independent tests. Positive samples exhibited characteristic sigmoidal amplification behavior, whereas negative samples remained linear and near baseline; we therefore classified reactions as positive or negative using a fixed hue threshold of 3 units, which clearly distinguished amplifying from non-amplifying reactions.

**Figure 6.**
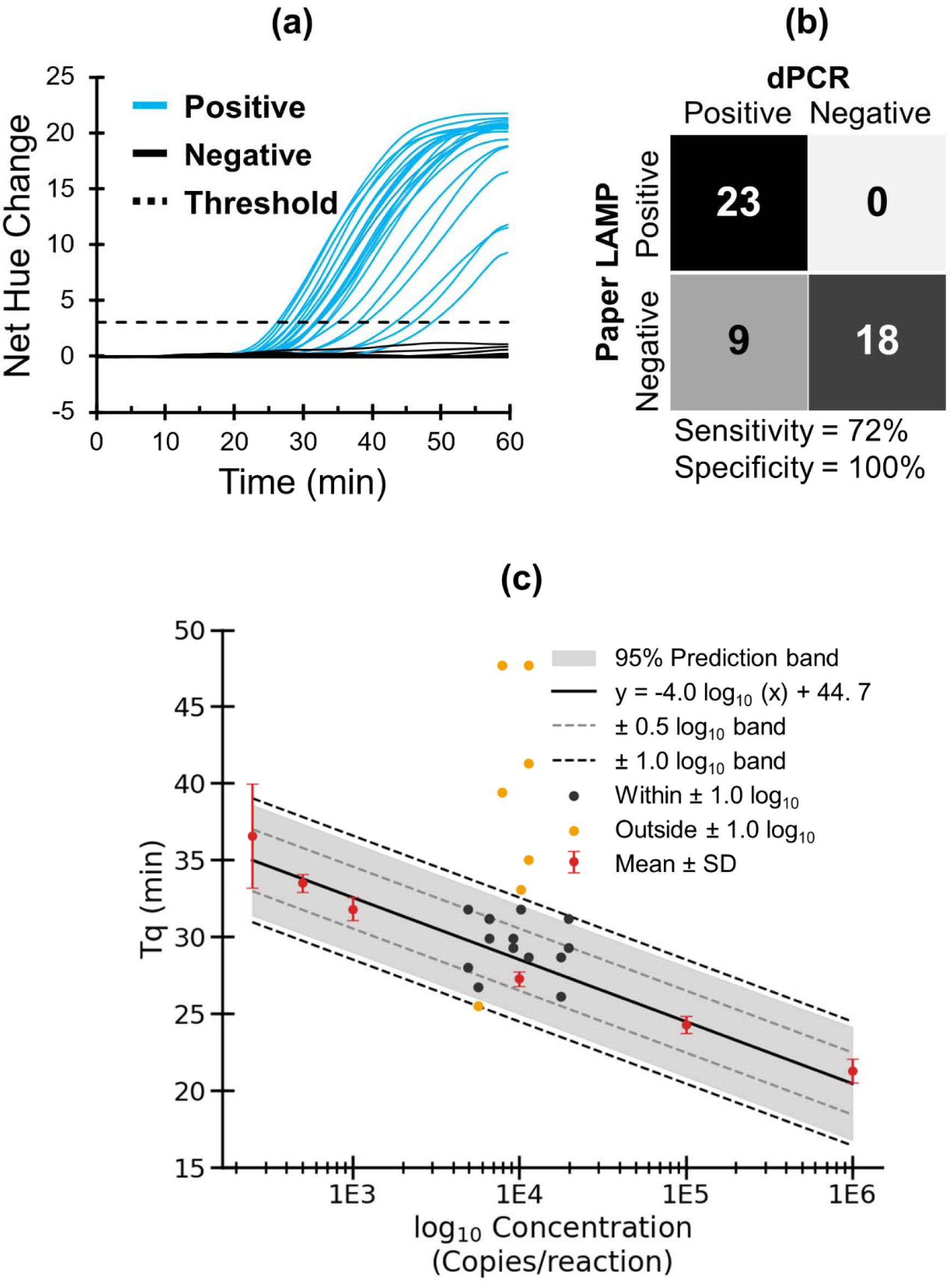
Comparison of paper-based qLAMP with digital PCR for SARS-CoV-2 virus samples. (A) Net hue changes over time from paper qLAMP timelapse images, with reactions classified as positive (blue) or negative (black) using a fixed threshold of 3 hue units. (B) Comparison of paper qLAMP test results with digital PCR (dPCR) ground truth. (C) Quantitative analysis using quantification time (Tq) versus log_10_ target concentration. The linear calibration curve was generated using serial dilutions of synthetic *orf7ab* DNA (mean ± SD, n = 6), with the 95% prediction band shown as a gray shaded region. Clinical SARS-CoV-2 virus samples quantified by dPCR are overlaid, with black points indicating Tq values within ±1.0 log_10_ of the calibration and orange points indicating values outside this range; gray dashed lines denote ±0.5 log_10_ and black dashed line show ±1.0 log_10_ confidence bands.

We then compared paper-based RT-LAMP classifications with dPCR. In dPCR, we considered samples positive only when both replicate measurements showed detectable target, to account for stochastic detection at very low copy numbers (<1 copy per reaction). Using this binary comparison (Figure 6B), the paper-based RT-LAMP assay achieved a specificity of 100% and a sensitivity of 72%. All false-negative results corresponded to samples with dPCR-estimated concentrations near the LOD95 (34 copies per reaction).

### Limited quantitative transferability of DNA-based calibration curves to clinical samples

We evaluated whether the standard calibration curve relating Tq to log_10_ input DNA concentration, established using synthetic *orf7ab* target DNA (Figure 5), could predict viral concentration in clinical samples. To this end, we constructed a single linear calibration curve using all technical replicates (n = 6 per concentration) above the LOQ by pooling data from two independent experiments (Figures 4&5); we show the calibration as mean ± SD with the corresponding linear fit and 95% prediction band (Figure 6C). Across the calibrated range, the 95% prediction band largely remained within ±0.5 log_10_ units, a range commonly considered acceptable for relative quantification in amplification-based assays (for example, qPCR calibration precision and assay variability are often on the order of 0.2–0.5 log_10_ in method verification studies and standard curve analyses)^43^.

When we overlaid dPCR-quantified clinical measurements onto this calibration curve using their corresponding Tq values extracted from paper-based colorimetric RT-LAMP hue–time trajectories, only 33.3% of Tq from dPCR-positive measurements fell within ±0.5 log_10_ band of the calibration curve. Expanding the acceptance window to ±1.0 log_10_ increased agreement to 61.9%, corresponding to quantification within a tenfold range. These results indicate that, although calibration curves obtained using synthetic DNA targets provide repeatable quantitative performance for purified samples, their direct transfer to clinical paper-based RT-LAMP measurements yield limited quantitative accuracy, likely due to sample matrix effects and substrate-related variability.

## Discussion

NAATs on paper substrates have lacked a high-throughput (>100 reactions) laboratory-grade reference instrument comparable to commercial thermocyclers. While liquid-based thermal cycler systems such as QuantStudio 6 & 7 Pro, CFX Opus 384, and Cielo3 provide precise (< ±0.5 °C) temperature control and real-time (every few seconds) fluorescence detection for 96–384 samples, no analogous platform exists for colorimetric isothermal assays on µPADs. We developed the ThermiQuant™ MegaScan, together with Amplimetrics™ software, to fill this gap by enabling scalable, real-time kinetic analysis of paper-based NAATs under controlled laboratory conditions. The platform serves as a proof-of-concept research tool to support systematic optimization and benchmarking of paper-based amplification assays rather than as a portable clinical diagnostic device. Translation into a fully integrated standalone system would require additional hardware and software integration as well as regulatory development, which was beyond the scope of this study. From the instrument’s implementation across synthetic and clinical workflows, we identified four key insights (i) robust and uniform heating with consistent imaging can be achieved using simple, low-cost (< USD 1,600) instrumentation; (ii) hue serves as an interpretable and reproducible optical metric for phenol red–based LAMP amplification chemistry; (iii) quantification on paper occurs in distinct kinetic regimes, in which reactions near the LOQ show high variability, while quantification repeatability emerges only at higher template concentrations (>250 copies/reaction in this study using synthetic target); and (iv) calibration curves derived from purified synthetic DNA targets do not reproducibly predict concentrations in clinical samples.

A true high-throughput NAAT analyzer requires a transparent heating element that enables simultaneous temperature control and optical detection. To achieve this requirement, several designs (as summarized in Table 1) have implemented single-sided heating using thin-film resistive elements while reserving the opposite face for imaging or sensing. Although not all reports specify heating precision, those that do generally achieve ±1 °C accuracy. However, single-sided heaters inherently suffer from thermal gradient bias. For instance, the authors of UbiNAAT reported that their device required a heater setpoint of 71.5 °C to maintain an internal chamber temperature of 63 °C under ambient laboratory conditions^34^. This implies that when environmental conditions vary, such as during outdoor use, calibration and setpoint adjustments are necessary for each use case. We address this limitation by employing water-bath heating, which leverages the high specific heat capacity of water (4,184 J/kg·°C) to provide stable thermal regulation^44^. When coupled with a circulator, the system maintains temperature uniformity within ±0.5 °C (Figure 2A-C). Although we did not specifically evaluate the influence of ambient temperature in this study (as MegaScan is intended for standard indoor laboratory use), our earlier IsoHeat study^45^ tested a similar circulator-equipped water-bath system across outdoor environments ranging from 5 to 33 °C and observed less than ± 0.5 °C deviation in set temperature. However, the main drawbacks of water-bath heating are its slower thermal ramping, the need for water replacement, and higher power consumption. In our setup, the system required approximately 750 W of power, water replacement on a weekly or as-needed basis, and a 10–15 min (4.5 to 3.0 °C/min) ramp-up time to reach 65 °C. Consequently, the cartridge was introduced into the bath only after the target temperature was reached.

For imaging, we moved away from camera-based systems, as our experience with the FARM-LAMP platform revealed issues with uneven illumination, lens distortion, and shadow artifacts caused by reflections, particularly when imaging through a water-bath setup^21^. Instead, we employed a CCD flatbed scanner (Epson Perfection V850 Pro) that provides geometrically accurate and uniform imaging across the scan area, with variation within ±1 hue unit across the 0-30 hue range (Figure 2D, E). In reflective scanning mode, the scanner’s focal plane is typically located at the glass surface and image gradually goes out of focus as the imaging height increases (Figure 2F). Because the manufacturer does not specify the exact focal depth, we experimentally characterized the effect of imaging height on color fidelity in µPADs and found that µPADs positioned up to 15 mm above the scanner exhibited less than 7% total hue variation (∼0.5% per mm) between positive (yellow) and negative reactions. In our setup, the cartridge was positioned 7 ± 0.5 mm above the scanner, and for this placement tolerance the corresponding hue variation was only ∼0.25% between runs, sufficient for reproducible colorimetric analysis. Furthermore, since LAMP reactions are typically sampled every 30–60 seconds^32,46^, the scanner’s 15 s acquisition time per cartridge provides adequate temporal resolution for real-time monitoring.

We used hue as the analytical parameter to track phenol red-based colorimetric changes (red/pink to yellow) in µPAD LAMP assays because it isolates chromatic information from brightness and remains less sensitive to illumination variability than raw RGB intensity, as demonstrated in several prior studies^32,46^. Since OpenCV defines hue on a 0–180° scale where red spans both 0–10° and 160–180°, we rescaled the 160–180° to –20–1° to obtain a continuous red-to-yellow rescaled hue range. In our experiments, the rescaled hue correlated linearly with pH (R^2^ = 0.97) across the phenol red transition interval, similar to the linear trend we previously measured by spectrophotometry^44^. This finding confirms hue’s direct correspondence to the underlying amplification chemistry. Importantly, our results in Figure 4B show that amplification trajectories consistently display sigmoidal shapes for positives and linear for negatives similar to real time real-time qPCR^27,47^, reinforcing hue as an interpretable and transferable optical metric for colorimetric assays.

The hue-time trajectories for positive reactions can be further used to construct a standard calibration curve for predicting unknown DNA concentrations, thereby enabling quantitative measurement. To achieve this quantification, we identified the time point of maximum acceleration corresponding to the inflection of the sigmoidal curve by evaluating the second derivative of the hue–time profile (see SI Notes 1.4 and Figure S5). This time point, defined as the Tq, serves as an analog to the cycle threshold (Ct) or quantification cycle (Cq) in real-time qPCR. Plotting Tq values against the logarithm of input DNA concentration produced a linear correlation relationship for synthetic *orf7ab* target. Typically, a global linear fit is applied to estimate unknown template concentrations. However, our results with colorimetric paper LAMP revealed that amplification on paper occurs in two distinct kinetic regimes. For synthetic targets at low copy numbers below the LOQ (<250 copies/reaction), replicate variability was high, reflecting stochastic amplification onset and heterogeneity within the paper substrate. In contrast, at higher template inputs above the LOQ (250 copies/reaction), Tq values showed perfect repeatability across independent runs (R^2^ = 0.95; bias <1 min, Figure 5C), with Bland–Altman analysis confirming bias within the ±2 min instrument tolerance. This two-regime behavior supports a conceptual model in which paper-based NAATs provide reliable qualitative classification near the LOD95 (34 copies per reaction in this study), while reproducible quantitative calibration is achievable only in the high-copy regime for purified targets. The divergence in behavior between low- and high-copy regimes arises largely from substrate heterogeneity in paper, which amplifies stochastic variability at low template concentrations^48^.

In regard to analytical sensitivity, our LOD95 of 34 copies per reaction (5 copies/µL or ∼5,000 copies/mL) for purified synthetic targets is less sensitive than that of many established laboratory NAAT platforms, including high-throughput real-time PCR based assays that have demonstrated limits of detection as low as 10–74 copies/mL in controlled comparative evaluations of commercial systems and 167–511 copies/ml for sample-to-answer type system^49^. This difference likely reflects slower diffusion and spatial constraints inherent to the paper matrix, indicating the need for further improvements to lower the effective LOD. ThermiQuant™ MegaScan and Amplimetrics™ therefore provide a high-throughput framework for assay developers to better understand these limitations and to focus optimization efforts on strategies that enhance analytical sensitivity.

Clinical sample evaluation further revealed that this quantitative framework does not transfer robustly from synthetic standards to real samples. When applied to clinical SARS-CoV-2 samples processed in transport media, dPCR-quantified samples frequently deviated from the synthetic calibration curve, even when clearly positive (Figure 6). These deviations indicate that sample matrix-related effects introduce additional variability that compromises the predictive accuracy of Tq-based calibration in the clinical domain. Together, these findings demonstrate that while paper-based LAMP supports robust binary detection above its LOD, absolute concentration prediction using calibration curves derived from purified targets remains limited in clinical samples. In this context, instruments such as ThermiQuant™ MegaScan play a critical role by enabling objective, high-throughput kinetic analysis that defines both the capabilities and boundaries of quantitative interpretation on paper substrates.

While MegaScan supports relative quantification in the high-copy regime (≥250 copies/reaction), absolute quantification remains best achieved through digital PCR (dPCR)^50,51^. Unlike bulk amplification, dPCR partitions reactions into thousands of nanoliter compartments, each containing zero or one template. By applying Poisson statistics to the fraction of positive partitions, dPCR avoids calibration curves and minimizes stochastic effects, enabling absolute quantification. In contrast, MegaScan complements dPCR by supporting high-throughput relative quantification on paper substrates, making it valuable for assay development tasks such as primer benchmarking, reagent optimization, and stability testing.

Taken together, these findings establish MegaScan not as a replacement for portable point-of-need devices, but as a complementary laboratory reference framework. Its throughput demonstrated consistency across runs, and automated analysis position it to guide assay developers in calibrating new nucleic acid chemistries, validating novel paper formats, and benchmarking emerging diagnostic devices. Importantly, this work also defines the practical boundaries of paper-based quantification and highlights directions for improving classification, miniaturization, and sample validation.

Future work will focus on three areas: (i) software enhancements, including machine-learning–based automated positivity classification and streamlined Amplimetrics™ analysis; (ii) hardware development toward more compact and integrated imaging configurations, including alternative heating mechanisms to replace the current water bath design; and (iii) expanded test across broader clinical sample types and matrices.

## Methods

### ThermiQuant™ MegaScan design, fabrication, and assembly

The ThermiQuant™ MegaScan instrument has eight major components: (i) a flatbed scanner (Epson Perfection V800, B11B223201), (ii) a 3D-printed tank base (“tank_base.step”), (iii) a rimless borosilicate glass water bath (Awxzom nano tank, B0D3F36CTY), (iv) a 3D-printed cartridge holder (“cartridge_holder_base.step”, “cartridge_holder_handle.step”, and “M3_screw_knob.step”, see Figure S2 for assembly), (v) a laser-cut acrylic cartridge, (vi) a 3D-printed tank lid (“tank_lid.step”), (vii) a consumer-grade water heater rod (Anova Precision® Cooker Nano 3.0, AN425-US00), and (viii) a Windows-based personal computer (Dell, USA). Exploded views of the assembled instrument are shown in Figure 1A and Figure S1. CAD files are available in Mendeley Dataset.

All 3D models were designed using SolidWorks (Dassault Systèmes SolidWorks Corp., USA). 3D-printed components were fabricated from polyethylene terephthalate glycol-modified (PETG) and polycarbonate using a Bambu Lab X1 Carbon 3D printer (Bambu Lab, China) at 100% infill density. Polycarbonate did not warp in hot water (65 °C) and was therefore used for parts directly exposed to water (cartridge holder and tank lid), whereas PETG was used for components not in direct contact with water (tank base). Schematics of 3D printed parts are shown in Figure S2.

Detailed assembly and operation procedures are described in supplementary information (SI) Notes 1.1 and illustrated in Figure S1, with the complete bill of materials (BOM) provided in Table S2. For time-lapse image acquisition, the instrument utilized VueScan software (Hamrick Software, USA; professional license required and optimized software settings shown in Table S1). The software operation workflow is detailed in SI Notes 1.2, and the overall device operation is summarized in Figure 1C. Captured images were processed using Amplimetrics™, a custom Python-based software application. Installation and usage instructions for the Amplimetrics™ analysis module are described in SI Notes 1.3, and the associated algorithmic framework is detailed in SI Notes 1.4.

### µPAD strip fabrication

A µPAD strip without LAMP reagents consisted of four structural components: (i) 0.83-mm-thick Grade 222 chromatography paper pads (Grade 222, Ahlstrom-Munksjö, Finland), (ii) 0.508-mm-thick polystyrene spacers (HIPS Litho Grade sheets, Tekra, USA), (iii) a double-sided adhesive layer (ARclean® 90178, USA), and (iv) a 0.076 mm optically clear MELINEX® 454 polyester film (Tekra, USA). An exploded view of the µPAD strip assembled within the acrylic cartridge is shown in Figure 1B.

Chromatography paper and polystyrene spacer strips were cut using a leather strip-belt-cutting machine (Zhixumm, B0D8THH9LW, Amazon, USA) equipped with a 3 mm blade-to-blade spacing. Figure S3 provides a step-by-step guide to the preparation and assembly of the µPAD strip.

### Cartridge design and fabrication

The acrylic cartridge was designed to hold the µPAD strip and to prevent evaporation and water leakage. Designs were created in SolidWorks, exported as DXF files, and further edited in Adobe Illustrator (Adobe Inc., USA) before conversion to SVG format for laser cutting. A 1.5-mm-thick acrylic panel (Outus, B09MRFHFTR, Amazon, USA) was laser-cut using a Glowforge Plus 40 W laser cutter (Glowforge Inc., USA) with optimized settings (100% power, speed factor 160). Following laser cutting, acrylic panels were rinsed with reverse osmosis (RO) water, treated sequentially with 70% ethanol and RNase AWAY™ (Thermo Fisher Scientific, USA), and wiped with lint-free Kimwipes® (Kimberly-Clark, USA). Depending on the µPAD strip configuration (2-, 3-, 4-, or 8-pad designs) and slot type (full or half), the cartridge design can be easily modified. Figure S4A illustrates representative examples of 8-pad and 3-4-pad µPAD cartridge layouts while Figure S10 shows 2 µPAD high throughput layout.

### Copy number calibration by digital PCR (PCR)

Synthetic DNA corresponding to the SARS-CoV-2 *orf7ab* target sequence (Table S4; NCBI Reference Sequence: NC_045512.2) was obtained as 4 ng lyophilized pellets (IDT, USA) and rehydrated to a final stock volume of 40 μL with nuclease-free water (Fisher Scientific, 43-879-36). De-identified remnant human nasopharyngeal (NP) swab samples that tested positive or negative for SARS-CoV-2 by the FilmArray RP2.1 multiplex real-time PCR assay (bioMérieux) were obtained from the Indiana University Health Division of Clinical Microbiology between October 2024 and March 2025. The swab samples were stored at −80 °C in Remel Viral Transport Medium (VTM) or Copan Universal Transport Medium (UTM). Specimen de-identification, storage, and transportation were performed in compliance with Indiana University Institutional Review Board (IRB) #16895 and Purdue University IRB-2022-1452.

dPCR primers and probes were custom designed using the PrimerQuest™ online tool (https://www.idtdna.com/PrimerQuest/Home/Index) following recommended settings mentioned in the online tool and the primers and probes are listed in Table S5. Each dPCR reaction for the synthetic DNA target was prepared in a total volume of 40 μL, consisting of 10 μL of 4× Probe PCR Master Mix (Qiagen, 250102; final concentration 1×), 4 μL of 10× primer–probe mixture (final concentration 1×; 0.8 μM forward primer, 0.8 μM reverse primer, and 0.4 μM FAM-labeled probe; see Table S5), 0.5 μL of *Eco*RI-HF restriction enzyme (New England Biolabs, R3101S), 20.5 μL of nuclease-free water, and 5 μL of DNA template. The reaction mixtures were dispensed into a 26K 24-well Nanoplate (Qiagen, 250001) and processed on a QIAcuity One 5-plex dPCR platform (Qiagen, 911021). The amplification program consisted of an initial denaturation at 95 °C for 2 min, followed by 40 cycles of 95 °C for 15 s, 55 °C for 15 s, and 60 °C for 30 s. Quantification of the absolute DNA copy number was carried out using the QIAcuity Software Suite (Qiagen). The resulting synthetic DNA stock solution was subsequently stored at −80 °C until further use.

For the diluted NP swab samples containing SARS-CoV-2, each dPCR reaction was prepared in a total volume of 12 μL, consisting of 3 μL of 4× Probe PCR Master Mix (Qiagen, 250132; final concentration 1×), 0.12 μL of 100x One Step advanced RT mix, 0.6 μL of 20× primer–probe mixture (final concentration 1×; 0.8 μM forward primer, 0.8 μM reverse primer, and 0.4 μM FAM-labeled probe; see Table S5), 1.5 μL of Enhancer GC, 2.78 μL of nuclease-free water, and 4 μL of DNA template. The reaction mixtures were dispensed into an 8.5K 96-well Nanoplate (Qiagen, 250021) and processed on a QIAcuity One 5-plex dPCR platform (Qiagen, 911021). The amplification program consisted of an initial reverse transcription at 50 °C for 40 min, followed by RT enzyme inactivation at 95 °C for 2 min, then 40 cycles of 95 °C for 15 s, 55 °C for 15 s, and 60 °C for 30 s. All sample preprocessing for remnant specimens were conducted under enhanced biosafety level 2 (BSL-2+) conditions.

### Preparation of LAMP and RT-LAMP reactions and assembly of µPADs into cartridge

The colorimetric LAMP formulation used for the paper-based assay was adapted from our previous work^41^. To prepare 1,000 μL of 2× LAMP mix, the solution was prepared by combining 100 μL KCl (1,000 mM; Sigma-Aldrich, P9541), 160 μL MgSO_4_ (100 mM; Sigma-Aldrich, M2773), 280 μL dNTP mix (10 mM; Fisher Scientific, FERR0182), 2.8 μL dUTP (100 mM; Fisher Scientific, FERR0133), 0.4 μL Antarctic Thermolabile UDG (1 U/μL; New England Biolabs, M0372S), 5.4 μL Bst 2.0 DNA polymerase (120 U/μL; New England Biolabs, M0537M), 20 μL phenol red (25 mM; Sigma-Aldrich, P3532), 100 μL Tween-20 (20%; Sigma-Aldrich, P9416), and 331.4 μL nuclease-free water (Fisher Scientific, 43-879-36). For diluted NP swab samples containing SARS-CoV-2, this formulation was modified to enable reverse transcription by replacing the 40 µL of RO water with an equivalent volume of WarmStart RTx Reverse Transcriptase (Fisher Scientific, 43-879-36).

To prepare 200 μL of the LAMP final master mix, 125 μL of the 2× LAMP solution was combined with 25 μL 10× primer mix (16 μM FIP/BIP, 2 μM F3/B3, and 4 μM LF/LB; final concentrations 1.6 μM FIP/BIP, 0.2 μM F3/B3, 0.4 μM LF/LB; see Table S6), 0.67 μL Bst 2.0 DNA polymerase, 1 μL betaine (5 M; Sigma-Aldrich, B03005VL), 3.13 μL bovine serum albumin (BSA) (40 mg/mL; Sigma-Aldrich, A2153), 36.0 μL trehalose (1.75 M; Thermo Scientific Chemicals, 182550250), and 9.2 μL nuclease-free water. The pH of the master mix was adjusted to ∼7.9 with 0.1 M KOH, producing a uniform red solution.

A 7.5 μL aliquot of the final master mix was pipetted onto each µPAD and allowed to dry inside a PCR workstation (Mystaire, MY-PCR32) for 2 h. For loading the µPAD strips into the acrylic cartridge, one side of the acrylic cartridge was first sealed with PCR plate seal (Fisher Scientific, AB-0558). Using tweezers, µPAD strips preloaded with the LAMP master mix were placed into the cartridge. For future use, the assembled cartridges can be stored in airtight plastic sealing bags at −20 °C; however, in this study, cartridges were used on the same day of preparation.

### Paper LAMP and RT-LAMP reactions in ThermiQuant™ MegaScan

Two identical LAMP experiments were performed on the ThermiQuant™ MegaScan system to evaluate instrument’s functionality. The cartridge loaded with µPAD strips prepared as described in earlier mentioned methods was used. For this demonstration, a modified cartridge version of the full 160-µPAD design was utilized (Figure S4A, top left panel), with each µPAD strip containing three µPADs (Figure S4B-D). The dPCR-quantified synthetic DNA was serially diluted to a range of concentrations (10^6^, 10^5^, 10^4^, 10^3^, 5×10^2^, 2.5×10^2^, 10^2^, 5×10^1^, 2.5×10^1^, 10^1^, and 1 copy per reaction), and three technical replicates were tested for each dilution. A 7.5 μL aliquot of each dilution was used to rehydrate the µPADs. For no-template controls (NTCs), an equivalent volume of nuclease-free water was applied. Number of replicates of NTCs were different between the two experimental runs (three in Run 1 and six in Run 2). Figure S4 (B-D) shows the images before and after rehydrating the μPADs. After pipetting the DNA samples onto the µPADs, the cartridge was sealed on the open face with PCR sealing tape and placed into a 65 °C preheated water bath of the ThermiQuant™ MegaScan instrument. Time-lapse images were captured using the VueScan software as outlined in SI Notes 1.1-1.4.

For clinical sample analysis, reaction conditions and imaging procedures were identical to those used for synthetic DNA samples, except that samples stored in UTM or VTM were diluted to (5% v/v) in nuclease-free water prior to loading onto the µPADs. This dilution was selected to mitigate inhibitory and buffering effects associated with transport media. Each µPAD strip contained paired reaction pads corresponding to a sample reaction and a no-primer control (NPC, Figure S10). Duplicate paper RT-LAMP measurements were performed for each clinical sample except two samples where triplicate measurements were done. Samples were prepared using BSL-2+ procedures.

### YOLOv8 instance segmentation and Image analysis workflow

Time-lapse images captured by the ThermiQuant™ MegaScan were analyzed using the custom Python-based Amplimetrics™ software. The analysis workflow is described in SI Notes 1.3, and the software video guide is provided as Supplementary movie 1.

Instance segmentation of individual µPAD reaction zones was performed using the YOLOv8n pretrained convolutional neural network (Ultralytics implementation)^52^. Regions of interest (ROIs) corresponding to individual µPADs were manually annotated using the LabelMe annotation tool (https://labelme.io/) to generate a custom training dataset consisting of 240 scanner images. The model was fine-tuned on this dataset for 100 training epochs and evaluated on a held-out test set of 60 images containing >1,300 µPADs. Retraining would be required if µPAD geometry or color chemistry were substantially modified. For clinical sample analysis, the average hue values of both the NPC and sample µPADs were first computed at each time point. The hue of the NPC µPAD was then subtracted from that of the corresponding sample µPAD to correct for baseline color shifts arising from the sample matrix. The net hue change was subsequently calculated as the cumulative change in background-corrected hue relative to the value at 0 min. Duplicate technical replicates for each clinical sample were treated as independent observations in downstream paper-based LAMP analysis.

### Determination of LOD95 and LOQ

The LOD95 was estimated using probit regression analysis^53^, in which binary amplification outcomes (positive/negative) from two independent experimental runs (combined n = 6 replicates per concentration) were pooled and modeled as a function of log_10_-transformed input concentration. LOD95 was defined as the concentration predicted to yield 95% positive detection probability. The LOQ was determined based on quantitative precision of Tq values: for each concentration at or above the LOD95, replicate Tq values (n = 6 per concentration) were used to compute the coefficient of variation (CV% = 100 × standard deviation / mean), and the LOQ was defined as the lowest concentration at which CV ≤ 10%, with a minimum of three numeric replicates required for CV estimation. Although a CV of up to 35% is often accepted for qPCR assays^54^, studies evaluating LAMP assays have reported typical CV values of approximately 5–10% as indicative of acceptable repeatability across reactions and runs^55–58^. Based on these reports, a CV threshold of 10% was applied in this study. For standard calibration analysis, log-linear regression of Tq versus log_10_ concentration was performed using replicate-level Tq values at concentrations ≥ LOQ. The 95% prediction interval of the regression model was calculated to represent the expected dispersion of individual measurements.

### Statistical Analysis and assay performance evaluation

Statistical analysis was performed in Python (v3.11) using NumPy, pandas, and Matplotlib through a custom script. The aim was to evaluate the agreement and reproducibility of Tq between two independent paper color LAMP experiments using synthetic targets. For each run, replicate Tq values were summarized as means with standard deviations. A continuous piecewise linear regression of mean Tq versus log_10_ concentration was applied with a shared breakpoint at LOQ and assess segment-wise linearity. Run-to-run agreement was evaluated by scatter plots (Run 2 vs. Run 1) with standard deviation (SD) error bars to represent replicate variability, and by Bland-Altman analysis^42^ of the two piecewise segments to quantify bias and 95% limits of agreement (LOA), with vertical error bars representing the propagated standard error of the difference between means.

For clinical sample evaluation, diagnostic performance was assessed by direct comparison of paper-based qLAMP results with dPCR (designated as ground truth). Paper LAMP tests were classified as positive or negative based on the fixed positivity threshold that clearly separated the amplification and non-amplification looking curves. This result was then compared against dPCR-obtained results to generate the paper LAMP assay sensitivity and specificity. For quantitative comparison, Tq values obtained from clinical samples were overlaid onto the synthetic *orf7ab* calibration curve, and agreement was evaluated relative to ±0.5 log_10_ and ±1.0 log_10_ concentration bands.

### Water bath temperature characterization

Thermal equilibration and spatial uniformity of temperature across the cartridge were evaluated by placing four thermocouples diagonally across an acrylic cartridge at 32 mm spacing (Figure 2A). The cartridge was immersed in the water bath maintained at 65 °C, and temperatures were recorded continuously for 15 min using a multichannel data logger (AZ Instruments K-type thermocouple recorder with 8G SD card, Amazon, USA). Equilibration time and spatial variation across probes were calculated from these recordings.

### CCD scanner color and scan height characterization

A 4 × 5 grid of 3 mm × 3 mm printed color squares spanning the red–yellow range was glued in a custom 3D-printed holder positioned on the scanner platen. The grids were scanned at 600 dpi and analyzed in the hue saturation value (HSV) color space using Amplimetrics™ software, which uses the OpenCV (Python) library that distributes hue over 0–180° scale. Because the hue component in OpenCV is cyclic, with red hues appearing near both 0° and 180°, discontinuity occurs at the red boundary. To maintain a continuous red–yellow hue axis, hue values between 160° and 180° were linearly remapped to −20° to 1° according to Eq. (1) & Eq. (2):

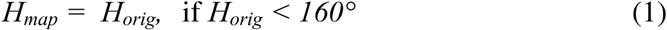

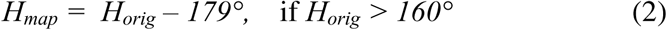

where *H_orig_* is the original hue value (in degrees), and *H_map_* is the remapped hue value used for color quantification. This transformation resolves the discontinuity at the red boundary by re-mapping hue values from the cyclic 0°–180° range to a continuous red-to-yellow scale (Figure S6). The depth of field of the imaging system was evaluated using a 3D-printed ladder mount that elevated the color grids in 3 mm increments from 0 mm to 15 mm above the glass surface.

### pH and color study

To assess the correlation between hue and pH in phenol red chemistry, LAMP final master mix (see earlier Methodology on LAMP reagent preparation) was adjusted between pH 6.0 and 8.4 using 0.1M KOH and 0.1M HCl. µPADs were pipetted with 7.5 µL with the adjusted mixes (0.2 pH interval), dried at room temperature for 2 h, and then rehydrated with 7.5 µL of nuclease-free water. µPADs were imaged using the ThermiQuant™ MegaScan setup, and hue values were extracted with Amplimetrics™ software.

## Supporting information

Supplementary Software

## Data availability

All relevant raw data presented here are available on Mendeley Data (DOI: 10.17632/thrhdkrbdt.1).

## Code availability

All relevant software code presented here are available in the Supplementary Software and in the Mendeley Data (DOI: 10.17632/thrhdkrbdt.1; “Software” folder).

## Acknowledgements

We are grateful to Dr. Rachel A Munds, Dr. Jiangshan Wang, Dr. Mohamed Kamel, Cindy Morigya, and Nafisa Rafiq for their participation in the initial user testing of the hardware and software tools and for providing insightful feedback. In addition, we are grateful for feedback on design and usability of the devices built here to the following people: Aaron Ault, Aaron Gilbertie, Dr. Patrick Zollner, Dr. Arezoo Ardekani, Dr. Darryl Ragland, and Dr. Timothy Johnson.

## Use of Generative AI

During the preparation of this work, the authors used Grammarly and ChatGPT to check for grammar errors and improve their academic writing language as well as debugging Amplimetrics™ software. After using this tool/service, the authors reviewed and edited the content as needed and take full responsibility for the content of the publication.

## Funding

This work was supported in part by the Foundation for Food and Agriculture Research under award number – Grant IDs: FF-NIA20-0000000087 and ICASATWG-0000000022. The content of this publication is solely the responsibility of the authors and does not necessarily represent the official views of the Foundation for Food and Agriculture Research. This work is partially supported by the Agriculture and Food Research Initiative Competitive Grants Program Award 2020-68014-31302 from the U.S. Department of Agriculture National Institute of Food and Agriculture. This project is supported partially by USDA’s Animal and Plant Health Inspection Service (APHIS) through the National Animal Health Laboratory Network (Sponsor Award # AP22VSD&B000C022). Funding for this project was provided partially by the American Rescue Plan Act through USDA APHIS (Sponsor Award # AP23OA000000C015). The findings and conclusions in this publication are those of the authors and should not be construed to represent any official USDA or U.S. Government determination or policy. This work was also partially supported by Purdue University’s Colleges of Agriculture and Engineering Collaborative Projects Program 2018, the College of Agriculture and Wabash Heartland Innovation Network Graduate Student Support program, an Agricultural Science and Extension for Economic Development (AgSEED) grant, and the Disease Diagnostics INventors Challenge (created by the Purdue Institute of Inflammation, Immunology and Infectious Disease in partnership with the Department of Comparative Pathobiology, which contributed the funds to realize the project, the Indiana Clinical and Translational Sciences Institute, and the Indiana Consortium for Analytical Science and Engineering). The work was partially supported by the Purdue University College of Engineering and Indiana University School of Medicine Engineering-in-Medicine Pilot Project Program. The project is also partially supported by Purdue University’s 2025 Bridge Funding program.

## Author contributions

BR: Conceptualization, software, data curation, formal analysis, investigation, methodology, validation, visualization, writing – original draft preparation. GP: Data curation, formal analysis, investigation, methodology, validation, writing – review and editing. VK: Data curation, investigation, methodology. AF: Software. BA: Data curation, writing – review and editing. JLD: Methodology. RFR: Methodology. JPS: Supervision, funding acquisition. JAP: Supervision. funding acquisition, validation, writing – review and editing. MSV: Conceptualization, supervision, project administration, funding acquisition, validation, and writing – review & editing. All authors have reviewed the manuscript and approved the final version.

## Competing Interest

A US patent related to aspects of this work has been filed (Application No. US20250222460A1). Inventors: Mohit S. Verma, Nafisa Rafiq, and Bibek Raut. Applicant/Assignee: Purdue Research Foundation. Status: Pending. The patent covers all aspect of the technology developed in this manuscript. MSV and JPS have interests in Krishi, Inc., which is a startup company developing molecular assays. Krishi, Inc. did not fund this work. M.S.V. has interests in Simply Experiment LLC, which did not fund this work either. The other authors do not have a competing interest.

## Additional Information

### Supplementary information

All essential data is included in the supporting documents.

### Copyright claim

ThermiQuant™ and Amplimetrics™ are copyrighted by © Purdue Research Foundation 2024.

## Notes

https://data.mendeley.com/datasets/thrhdkrbdt/1

### Summary of Updates

Additional data was included to strengthen the manuscript.

