## Supporting Information for "ThermiQuant™ MegaScan: High-throughput isothermal reactor with quantitative colorimetric readout for paper-based nucleic acid amplification tests"

for

### 1. Supplementary Notes

#### 1.1 ThermiQuant™ MegaScan assembly and operation manual

This manual provides a step-by-step guide to the first assembly and regular operation of the ThermiQuant™ MegaScan, as well as a cleanup guide after use. Figure S1 visually shows these steps and Figure S2 shows the 3D design schematics of 3D printed parts. CAD design files are in the “Design_files” folder of the dataset.

For first-time assembly, refer to the exploded view showing: (1) Scanner, (2) Tank base, (3) Tank, (4) Cartridge holder, (5) Cartridge loaded with µPAD strips, (6) Tank lid/top cover, and (7) Heater rod.

##### Assembly Instructions

i. Power on the scanner and connect the USB cable to a personal computer (PC). Launch the VueScan software (professional license required) and verify that the scanner is detected and able to scan. Refer to Supplementary Notes section 1.2 (VueScan software installation and use with ThermiQuant™ MegaScan) for details on first-time installation of VueScan and integration with the ThermiQuant™ MegaScan platform.
ii. Remove the removable scanner cover (used to protect the scanner when not in use) and thoroughly wipe the scanner glass window with a lint-free wipe moistened with either water or 70% ethanol.
iii. Set the 3D printed tank base on the scanner so that the side marked Front Side faces you and the base window is centered over the scanner glass.
iv. Lower the water tank straight into the 3D printed base until it sits flush. Do not press too hard. If press fitting is required because of 3D printing imperfections, perform this outside the scanner to avoid overloading it.
v. Place the heater rod into the tank through its vertical port (illustrated as the blue rod). Ensure it is fully seated and unobstructed. The Anova heater rod includes a screw lock to secure it to the side wall-tighten gently, just enough for the rod to stand vertically.
vi. Fill the water tank with a 5-10 mL of surfactant (Tween 20) and Reverse Osmosis (RO) water. The surfactant reduces surface tension, preventing bubbles from attaching to the tank or cartridge surfaces during heating. RO water is recommended over tap water, which contains salts and impurities that quickly build up and accumulate in the heater rod; using RO water reduces the frequency of water changes. The water level should be within the minimum and maximum markings on the heater rod. Water is typically changed after ten runs or weekly or whenever it begins to look dirty. Set the water temperature to 65 °C using the Anova heater rod’s touch interface.
vii. Assemble the cartridge with µPAD strips, add samples, and seal with PCR sealing film (see Figure 1C in the main manuscript). To load the cartridge into the holder, insert the long edge forward into the slot. Ensure the cartridge lies flat with a sufficient gap underneath for water flow. If it touches the tank glass, heat exchange is blocked, creating temperature gradients and condensation inside the cartridge. Gently lock the four screw nuts to secure the cartridge. Do not place the assembled cartridge holder into the water bath yet.
viii. Prepare the VueScan software. Refer again to Supplementary Notes section 1.2 (VueScan software installation and use with ThermiQuant™ MegaScan) for details. Create folder names for saving images and perform a preview scan to select the region of interest (ROI). This step can be adjusted later but preparing it now saves time and prevents delays. Delays in capturing first image means an offset in LAMP kinetic analysis from timelapse images.
ix. Once the water has reached 65 °C, insert the cartridge holder in the water bath. Ensure it sits flat on the tank base and secure it to the side wall using the screw lock. Do not overtighten, as it risks breaking the 3D printed holder neck over time.
x. Place the top cover on the tank to minimize evaporation. Immediately return to VueScan. If the region of interest (ROI) was not previously selected, perform a preview scan and manually crop the ROI. Start the scan, then enable Autorepeat = 20 seconds to capture timelapse images. At the end of the experiment (usually ~60 minutes), disable timelapse by setting Autorepeat to “None”. Always check the destination folder to confirm that first few images are being saved, as misconfigured folder settings may cause file saving errors.
xi. Turn off the heater rod and unplug it from the power source. Verify that the scanner’s Autorepeat function is set to “None”. Doing so will stop the timelapse capture. Carefully remove the cartridge holder, taking care not to spill water. Use a Kimwipe® around the scanner to absorb drips and prevent water from seeping between the tank and the scanner glass.
xii. Remove the water tank from the scanner. Use heat resistant gloves or let the water cool to less than 40°C. While the scanner can tolerate continuous use, prolonged loading with a full water tank may cause the scanner surface to warp inward over several weeks due to weight and heat. To avoid this, do not operate continuously for more than three hours. If extended use is necessary, allow the scanner to cool by removing the tank between runs. Never store the water tank on the scanner when not in use, keep it separately on a desk with a Kimwipe® underneath to prevent scratches. Always keep the lid on to prevent dust accumulation inside the tank. Water is typically replaced after ten uses or weekly, or sooner if it becomes dirty. Wipe the scanner surface to remove any water droplets and dust. Cover the scanner with its protective cover/jacket when not in use.

#### 1.2  VueScan software installation and use with ThermiQuant™ MegaScan

This manual describes the step-by-step installation of the VueScan software and its use with the ThermiQuant™ MegaScan instrument for capturing timelapse images of the colorimetric LAMP reaction on the µPADs. Table S1 provides additional scanner settings and supplements this manual. The settings tested were from VueScan version 9.8.01. Table S1 shows scanner settings that are most critical for operation while leaving rest to default settings.

##### Software instructions

i. Go to <https://www.hamrick.com/> to download the VueScan software. A professional license is required to enable the timelapse feature. At the time of writing, the cost is approximately $130 for a single installation or about one-third of this price for an annual subscription.
ii. After installation, connect the Epson Perfection V800/850 scanner to the computer while ensuring the scanner is connected to a power supply. Verify that the software automatically detects the scanner and that a preview or test scan can be performed.
iii. Once the setup is confirmed, configure the settings as outlined in Table S1. The table highlights the key parameters; the headings correspond to the menus in the software. Leave all other settings at their default values. Always maintain these settings to ensure reproducible results. If modifications are necessary, use the same settings consistently throughout the entire experiment.
iv. Test the timelapse feature by first clicking Preview and then setting Autorepeat under the Input tab to Continuous. Save images in JPG format to minimize file size. No significant differences in image quality were observed between JPG and TIF formats, although TIF files were 20 times larger (refer to Figure 3D). To facilitate file management, keep folder names and filenames short and meaningful. Ensure that the full file path length (including folders and filenames) does not exceed 255 characters.

#### 1.3  ***Amplimetrics™ software installation manual***

AmpliMetrics™ is a Python-based software package that requires installation of essential software dependencies, along with two additional tools: Miniforge (Conda distribution) and Visual Studio Code (VS Code). We recommend using a Windows operating system, as the Amplimetrics™ software has been tested on Windows 11. Because software updates occur regularly, the instructions provided here should be treated as reference only. Both the Miniforge and Visual Studio Code download websites (linked in the manual below) include installation instructions, and we recommend consulting them for the most up-to-date versions.

##### Miniforge installation

i. Download and install the conda-forge/Miniforge installer from <https://conda-forge.org/download/>.
ii. Open the Miniforge Prompt from the Windows Start Menu and create a Python virtual environment named Amplimetrics with Python 3.11 by typing the following command. Respond to the prompt by typing y and hitting enter to continue:

***conda create --name Amplimetrics python=3.11***	# Creates new env with python 3.11

iii. Activate the environment by typing:

***conda activate Amplimetrics***	# Activates Amplimetrics environment

iv. Install the required libraries:

***pip install ultralytics***	# Tested with ultralytics Version: 8.3.187

***pip install pyqt5***	# Tested with pqqt5 Version: 5.15.11

***pip install pandas***	# Tested with pandas Version: 5.15.11

v. Reboot computer

##### VS Code installation and first-time setup

i. Download and install Visual Studio Code from <https://code.visualstudio.com/>. During installation, check “Add to PATH” (important for command-line usage). Install the Python Extension either during setup or later from the Extensions Marketplace.
ii. Launch VS Code from the Start Menu. Press Ctrl+Shift+X to open the Extensions Marketplace, search for Python, and install the extension by Microsoft.
iii. Open Microsoft Store app from start menu and look for Python 11 and install it.
iv. Press Ctrl+Shift+P to open the Command Palette, type ***Python: Select Interpreter***, and select environment Amplimetrics (the path may look like:

***C:\Users\YourName\anaconda3\envs\Amplimetrics-soft***

v. Download the Amplimetrics folder from the dataset (Software/Amplimetrics-V1.0). In VS Code, open this folder. Run the script “Amplimetrics_V1.0.py”. The graphical user interface (GUI) should appear within a few seconds (up to a minute on slower computers). If the GUI loads, installation is successful. For detailed usage, continue to the Amplimetrics User Guide below.

##### Amplimetrics user manual

A visual walkthrough is shown in Supplementary movie 1.

i. Open VS Code, navigate to the Amplimetrics folder, and run “AmplimetricsV1.0.py”. The GUI should appear within 10–15 seconds (up to 1 minute on first launch or on slower computers).
ii. The Amplimetrics interface has two sections: Image Processing and Data Processing.

###### Image processing:

This workflow converts time-lapse images of µPAD cartridges into per-pad hue time series. The output includes an annotated image, per-frame patch grids, and a CSV file of mean hue versus time.

i. Launch Amplimetrics and select the Image Processing tab.
ii. Click “Select Folder” and choose the directory containing the time-lapse images. Select the folder itself, not an individual image. The first image is displayed on the canvas on the right.
iii. In Chip Type, select Linear grid (rectangular array) to match the cartridge used. The software will automatically detect the location of each colored pad and assign alphanumeric labels. A metadata subfolder is created inside the image folder, containing “image_metadata.json” and an annotated image file “AnnotatedImage_with_labels.png.” This step runs automatically in the background using object detection models.
iv. Launch Amplimetrics and select the Image Processing tab.
v. Click “Process Images”. The software arranges all images in natural alphanumeric order and extracts date and time information from each file. If EXIF timestamps are missing (e.g., if images were edited or resaved), the software assigns time 0 to all frames. In this case, the user should manually edit the CSV file to overwrite the time column with the actual capture intervals. The resulting CSV (csv_hue_mean.csv) is saved in a “processed_data” subfolder and contains the time in minutes and the mean hue value for each µPAD. This CSV is required for downstream data processing. At the same time, the raw hue-versus-time data are plotted in the right canvas, and cropped image regions for each pad and timepoint are saved in the “combined_patches” subfolder.

###### Data Processing:

This workflow processes CSV files exported from the image analysis pipeline into baseline-corrected traces, applies thresholding, and computes quantitative values (Tq) using derivative and curve-fit strategies (Sigmoid fit, Voigt fit). The output includes processed plots, metadata-enhanced CSV files, and Tq result tables.

i. Launch Amplimetrics and select the Data Analysis tab.
ii. Click Load CSV and select the file “csv_hue_mean.csv” generated in the image processing step. The software automatically applies initial smoothing and baseline correction, displaying the raw data on the left canvas with shaded regions for ignored and baseline windows, and the processed data on the right canvas. A processed data table with metadata rows is also shown below the canvas.
iii. Adjust the baseline range by choosing the start (ignore time) and end (baseline stop) values from the dropdowns, then click “Smooth Normalize”. The smoothing and baseline correction are re-applied using the new parameters, and the plots and table are updated.
iv. Enter the total assay time in minutes under “Exp Duration” and click Apply. Data beyond this time are truncated, and a Threshold Method panel appears.
v. Apply a positivity threshold to differentiate positive from negative reactions. For a manual threshold, type a numeric value in the “Adjust Threshold” field and click Apply. For a control-based threshold (e.g., using no template control, NTC or no primer control, NPC), select “Suggest Threshold”. A panel with clickable label buttons will appear, showing up to six curves at a time. Click the labels corresponding to control data; selected labels are highlighted and displayed as dotted colored traces in the processed plot. Enter the multiplier (default = 5) and click Suggest Threshold. The suggested value is inserted into the threshold field; click Apply to update the plot. After applying, traces above the threshold line are classified as positive (sky-blue) and those below as negative (black). If the threshold does not appear appropriate, adjust the multiplier until there is clear visual separation between sigmoidal S-shaped curves (positive) and flat or linear curves (negative/control).
vi. Optionally, click “Process Derivative” to review data in batches of six. The display switches to a 2×3 grid where each sensor trace is shown along with derivative curves and fitted models if it crosses the threshold. The derivative method is used to determine the Tq, which is useful later for constructing calibration curves between Tq and initial template concentration. Use Previous and Next to navigate between batches. Control selection only affects the threshold suggestion in step 5 and does not alter this derivative view.
vii. Click Compute Tq to calculate quantification time. For positive curves, two values are reported: Tq_Voigt, derived from the second derivative of a Voigt fit to the first derivative, and Tq_Sigmoid, derived from the second derivative of a sigmoid fit to the raw trace. Results are accepted only if the fit quality is R^2^ ≥ 0.75; otherwise, the entry is marked as “Tq_failed.” Curves that never cross the threshold are reported as “No Tq.” Results are shown in a table along with metadata rows recording baseline times, threshold, and experiment duration.
viii. Save results by clicking “Save Tq Data”, which exports the table with metadata and sensor Tq values as <input>_Tq.csv, or by clicking “Save Processed Data”, which saves the baseline-corrected dataset with metadata rows as <input>_processed.csv.

#### 1.4  Amplimetrics™ Software algorithm for quantification time (Tq) estimation

This section describes how Amplimetrics software processes raw hue-versus-time colorimetric curves from LAMP reactions to classify positive and negative outcomes and calculate Tq. The algorithm first converts noisy raw signals into processed amplification curves, then distinguishes positive reactions from negatives using a positivity threshold. Tq values are estimated only for positive reactions, while negatives are excluded from further analysis and reported as “No Tq.” An overview of the workflow and representative examples of the fitting strategies are provided in Figure S5.

a. **Data Input and Zero-Referencing:** Raw hue-time data are imported from CSV files generated by the image processing pipeline (csv_hue_mean.csv). These files contain time values (first column, “Time_min”) and hue values from each reaction zone (columns labeled by alphanumeric IDs). As illustrated in Figure S5A (top panel), the raw traces begin at different initial hue values. To standardize comparisons across reactions, each trace is zero-referenced by subtracting its initial hue value from all subsequent points (Figure S5A, bottom panel). This transformation ensures that all curves start from a common zero baseline, representing the net hue change over time rather than absolute hue values.
b. **Signal Denoising and Smoothing:** To reduce noise while preserving reaction kinetics, the zero-referenced hue-time traces are smoothed in two stages. First, a centered 5-point moving average suppresses high-frequency fluctuations. Second, a Gaussian filter (σ = 2, reflect padding) further reduces residual irregularities, yielding smooth amplification curves that better reflect the underlying kinetics while minimizing the impact of measurement noise (Figure S5A, bottom panel).
c. **Ignore Region Selection and Baseline Correction:** The initial phase of a colorimetric LAMP reaction often displays irregularities caused by temperature ramping, pH shifts, or dye response artifacts, which produce transient dips or fluctuations unrelated to true amplification. To prevent these early artifacts from biasing quantification, Amplimetrics automatically designates the first few minutes (typically 3–10 minutes) as an ignore region, flattening values within this interval to zero. The boundary of this region is determined by examining the derivative (rate of change) of the hue curve and identifying when the signal stabilizes, though users may override this selection. Immediately following the ignored region, a baseline region is defined to capture the stable pre-amplification phase. This region is identified as the interval where the hue signal remains within ±1.5 standard deviations of its 5-point rolling mean. The baseline is then smoothed, interpolated, and subtracted from the entire curve to correct for slow drifts or background offsets, ensuring that subsequent hue increases represent true amplification. Users may also adjust this baseline manually if desired. Figure S5A (top panel, shaded regions) illustrates both the ignore region and the baseline region as evaluated by the software.
d. **Thresholding and Positivity Classification:** To distinguish true amplification from background variation, Amplimetrics applies a decision threshold. As illustrated in Figure S5A (bottom panel), this threshold is represented as a horizontal line against which each processed trace is evaluated. This threshold can be entered manually by the user or automatically suggested from negative control reactions using the formula:

***Threshold = μ_controls_+k σ_controls_***	(1)

where μ and σ are the mean and standard deviation of selected controls at the end of the experiment, and k (default = 5) is a user-defined multiplier. Outliers among control values are filtered using z-score or interquartile range criteria, depending on the number of controls. A reaction is classified as positive if its processed hue trace crosses this threshold at any time, while those that remain below the threshold are classified as negative. Only positive reactions proceed to Tq estimation.

e. **Tq Estimation by Sigmoid Fitting:** For positive reactions, Amplimetrics fits the processed hue–time curve to a logistic-type sigmoid function, constrained such that *F(0) = 0* (Figure S5B, top panel). This mathematical model captures the S-shaped growth typical of amplification. The fitted curve is then differentiated twice numerically, and the quantification time (Tq-sigmoid) is defined as the time point at which the second derivative reaches its maximum (Figure S5B, bottom panel). This corresponds to the time point of maximum acceleration in the amplification curve, marking the transition from baseline to exponential growth.
f. **Tq Estimation by Voigt Fitting:** As an alternative to the sigmoid approach, Amplimetrics calculates the discrete first derivative of the processed signal, which is often noisy, and fits it to a Voigt profile (Figure S5C, bottom panel). The Voigt function, a convolution of Gaussian and Lorentzian line shapes, is constrained to equal zero at t = 0. The Voigt-fitted first-derivative curve is then differentiated once more to obtain a second derivative, and the quantification time (Tq-voigt) is defined as the time at which this second derivative reaches its maximum. Like the sigmoid method, this identifies the point of greatest acceleration in the amplification curve but derived from the derivative profile rather than directly from the raw signal.
g. **Fit Quality Control:** While both the sigmoid and Voigt approaches provide robust strategies for Tq estimation, paper-based LAMP reactions can sometimes produce weak or atypical curves, particularly under low-template or very slow amplification conditions. In such cases, the fitted models may capture spurious features of noisy derivatives rather than genuine amplification kinetics. To minimize false detections, Amplimetrics applies a quality control step based on the coefficient of determination (R^2^). Only fits with R^2^ ≥ 0.75 are accepted as reliable; otherwise, the corresponding Tq is flagged as “Tq_failed.” Reactions classified as negative never undergo fitting and are reported as “No Tq.” This safeguard ensures that reported Tq values are based on robust amplification signals rather than noise. Looking ahead, in the future development we plan to incorporate more advanced statistical and machine learning approaches to recover useful information from borderline or noisy reactions and further strengthen Tq estimation.
h. **Output and Metadata:** For each reaction zone, Amplimetrics reports the estimated Tq values from both the sigmoid and Voigt methods, together with their corresponding R^2^ values as a measure of fit quality. In addition, key metadata are recorded, including the automatically or manually selected ignore and baseline regions, the applied decision threshold, the total experiment duration, and the unit of measurement (minutes). These outputs provide a comprehensive record of both the analysis parameters and the results, ensuring transparency, reproducibility, and interpretability of the reported Tq values.

### 2. Supplementary Figures

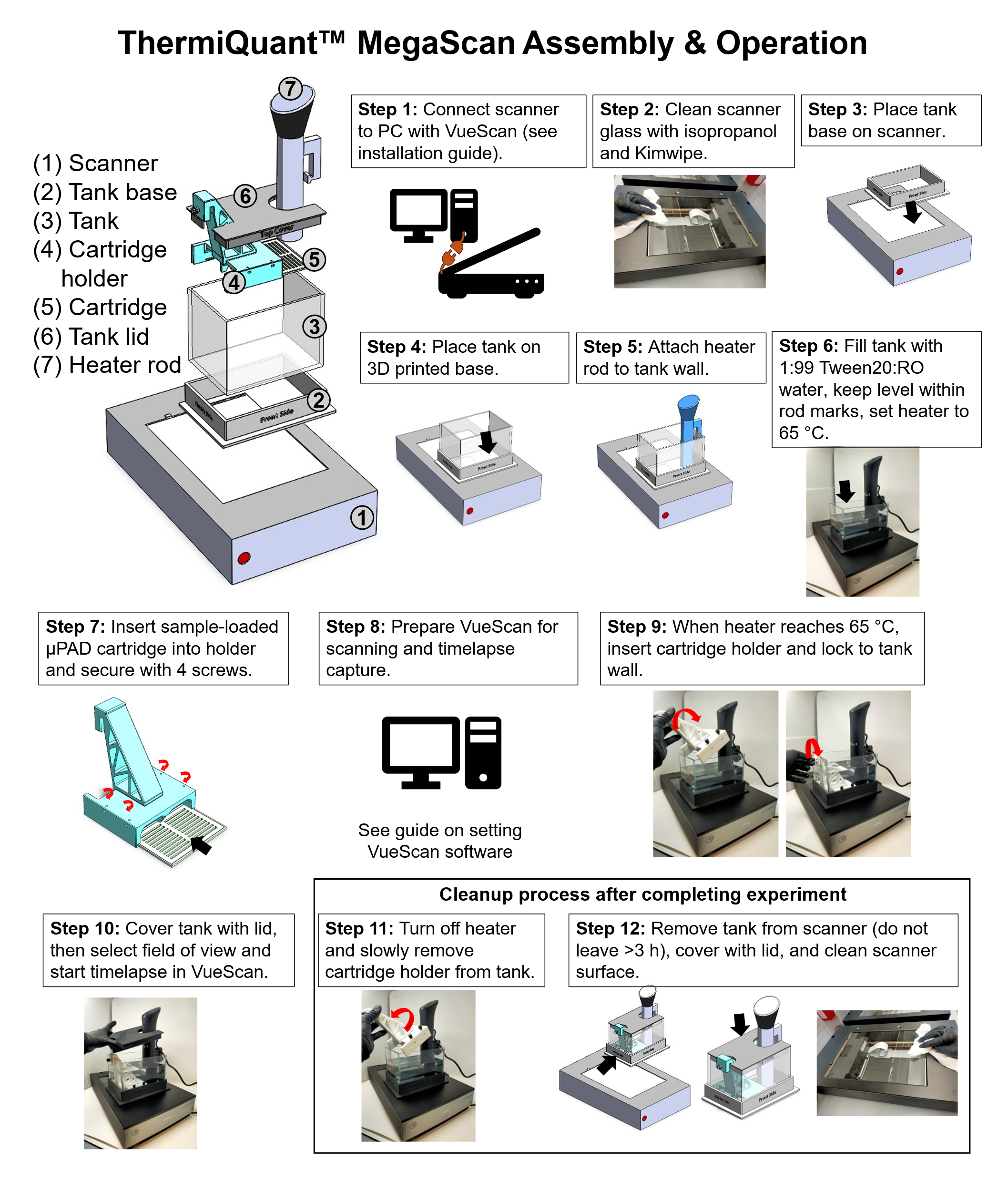

**Figure S1.** Step-by-step guide to ThermiQuant™ MegaScan assembly and operation. See Supplementary Notes 1.1 for details.

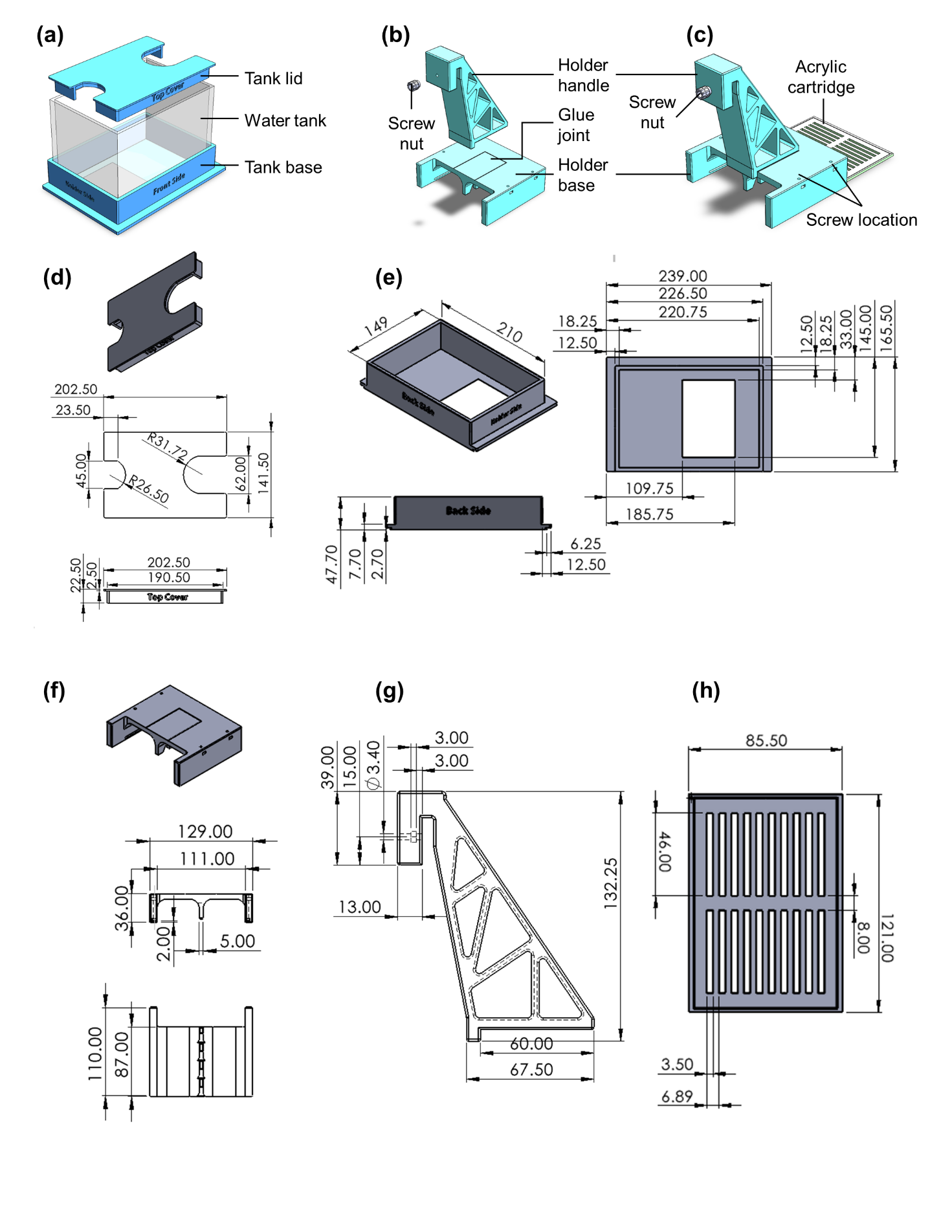
**Figure S2.** Drawings of 3D printed parts. (a) Schematics of tank, tank lid, and tank base. (b) Tank base. Holder handle and holder base were attached using superglue. (c) Cartridge in holder. Drawings showing (d) tank lid, (e) tank base, (f) holder base, (g) holder handle, and (h) cartridge. Dimensions are in mm.

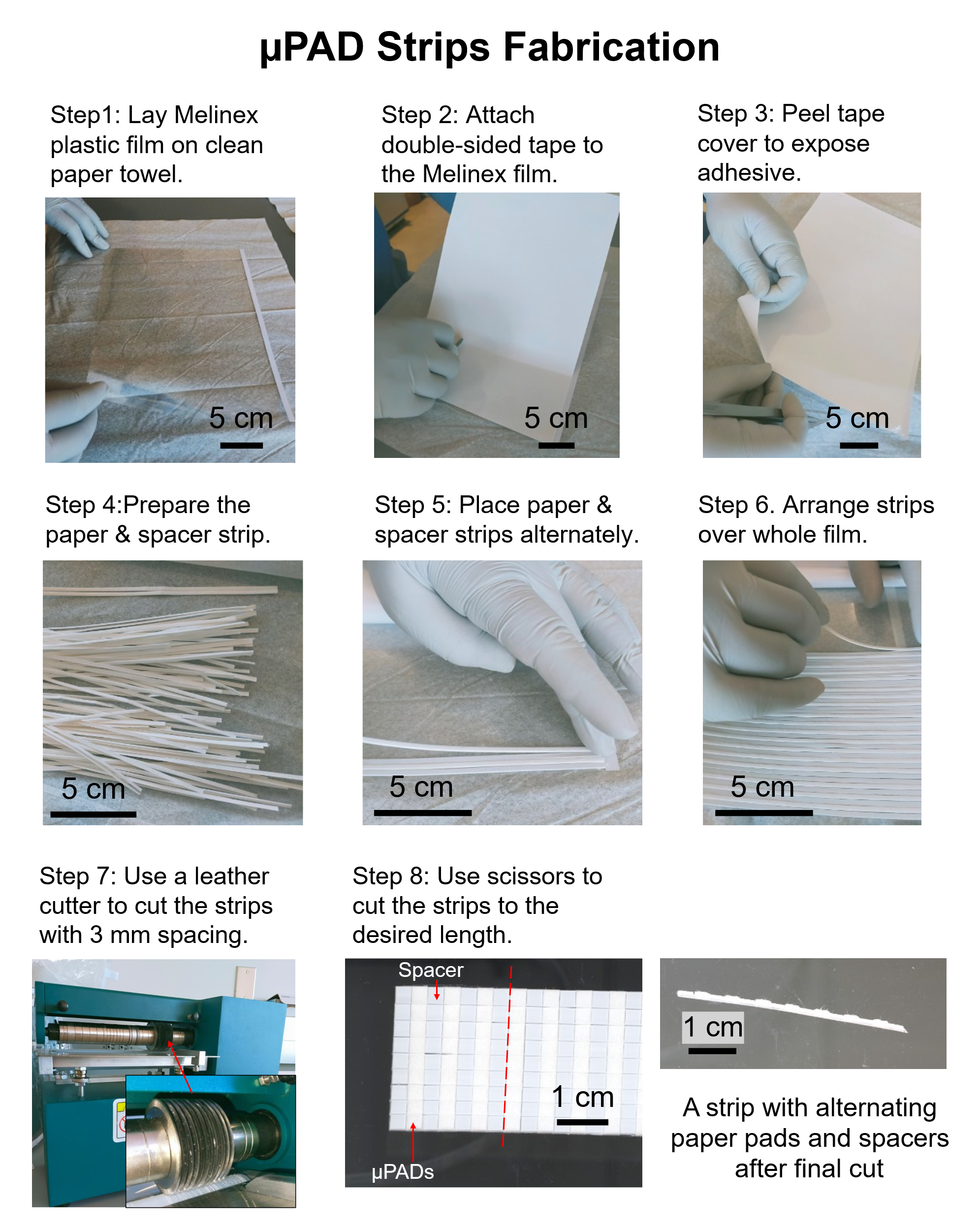

**Figure S3.** Step-by-step µPAD strip fabrication.

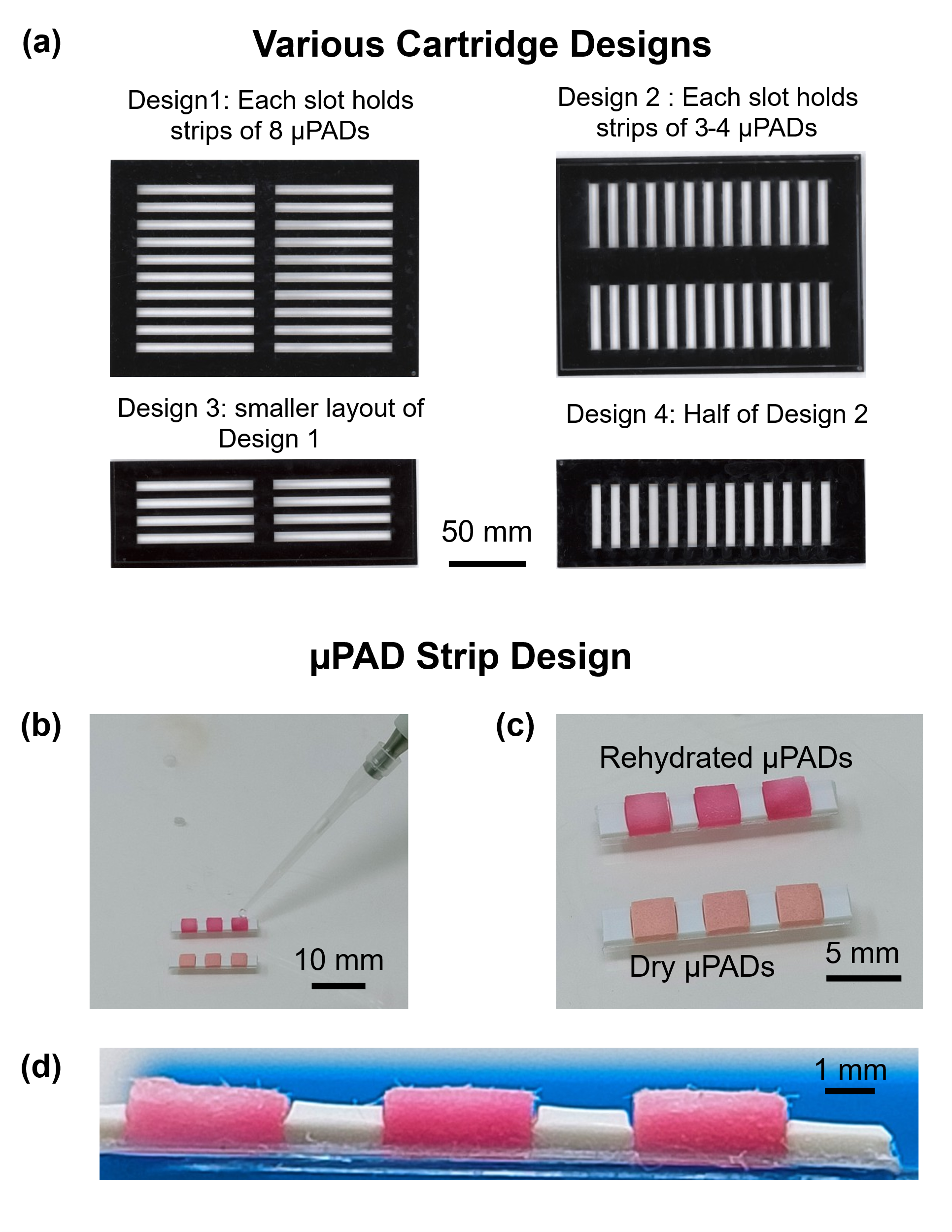

**Figure S4.** Cartridge and µPAD strip design. (A) Acrylic cartridge layouts for different numbers of µPADs and spacers, showing modularity for small- or large-scale assays. (B) Rehydrating a 3-strip µPAD. (C) Color µPADs before and after rehydration with predried reagents. (D) Side view of a µPAD strip with alternating paper pad and spacer.

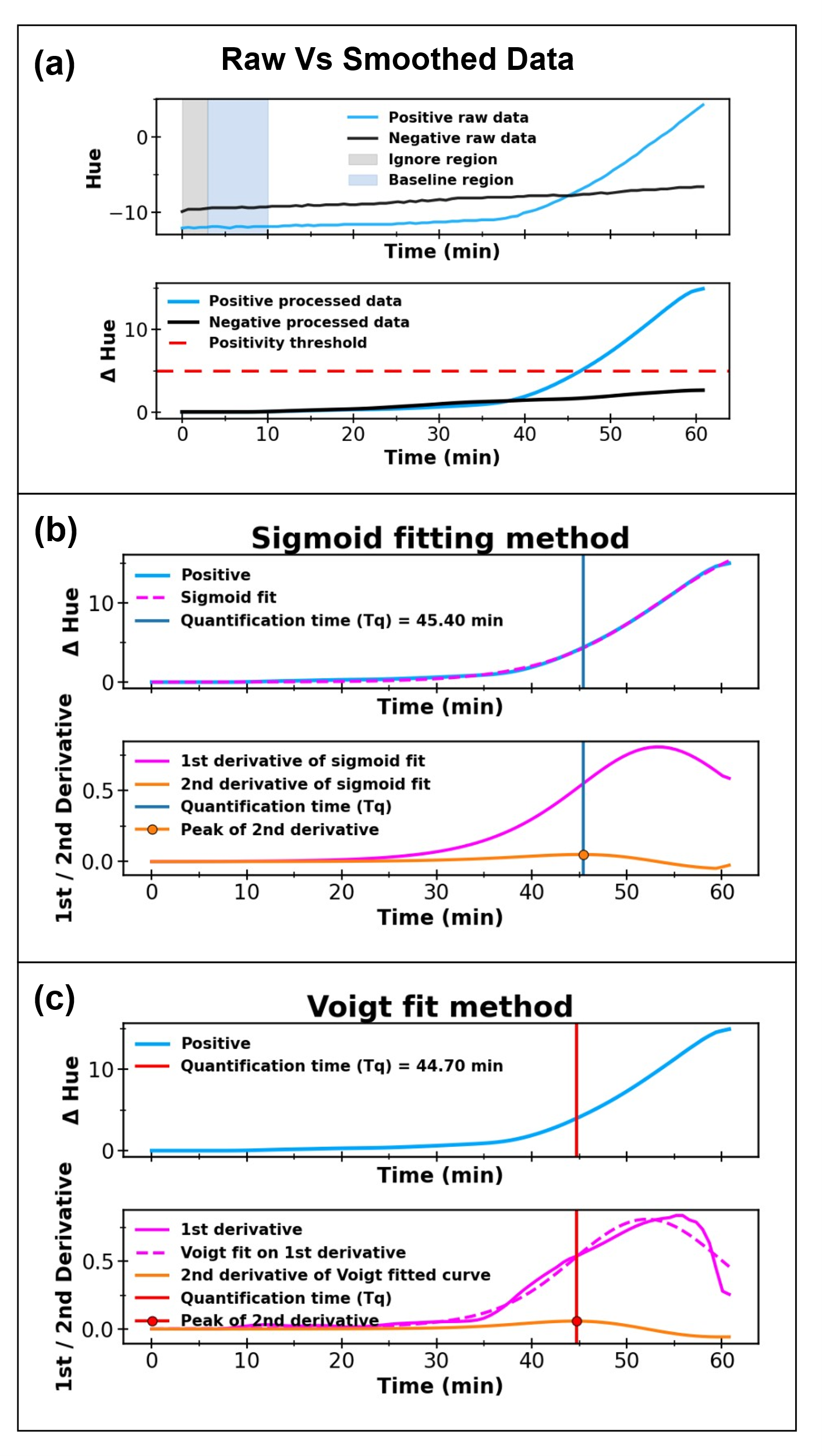

**Figure S5.** Amplimetrics workflow for quantification time (Tq) estimation. For step-by-step text description please refer to Supplementary Notes 1.4 (A) Raw and processed hue-time traces from a positive and a negative LAMP reaction. The top panel shows absolute raw hue values with shaded regions marking the ignore (gray) and baseline (blue) windows used during preprocessing. The bottom panel shows net hue change after zero-referencing and baseline correction, including the applied positivity threshold (red dashed line). (B) Sigmoid fitting method. Top panel: baseline-corrected amplification curve (blue) with the best-fit sigmoid model (green dashed line). Bottom panel: first derivative (magenta) and second derivative (orange) of the sigmoid fit. Tq is defined as the time at the maximum of the second derivative (orange dot), shown with a vertical blue reference line. (C) Voigt fitting method. Top panel: processed positive curve with Tq marked (red vertical line). Bottom panel: first derivative of the signal (magenta), Voigt fit to the derivative (green dashed line), and its second derivative (orange). Tq is defined as the time of the second-derivative peak (orange dot).

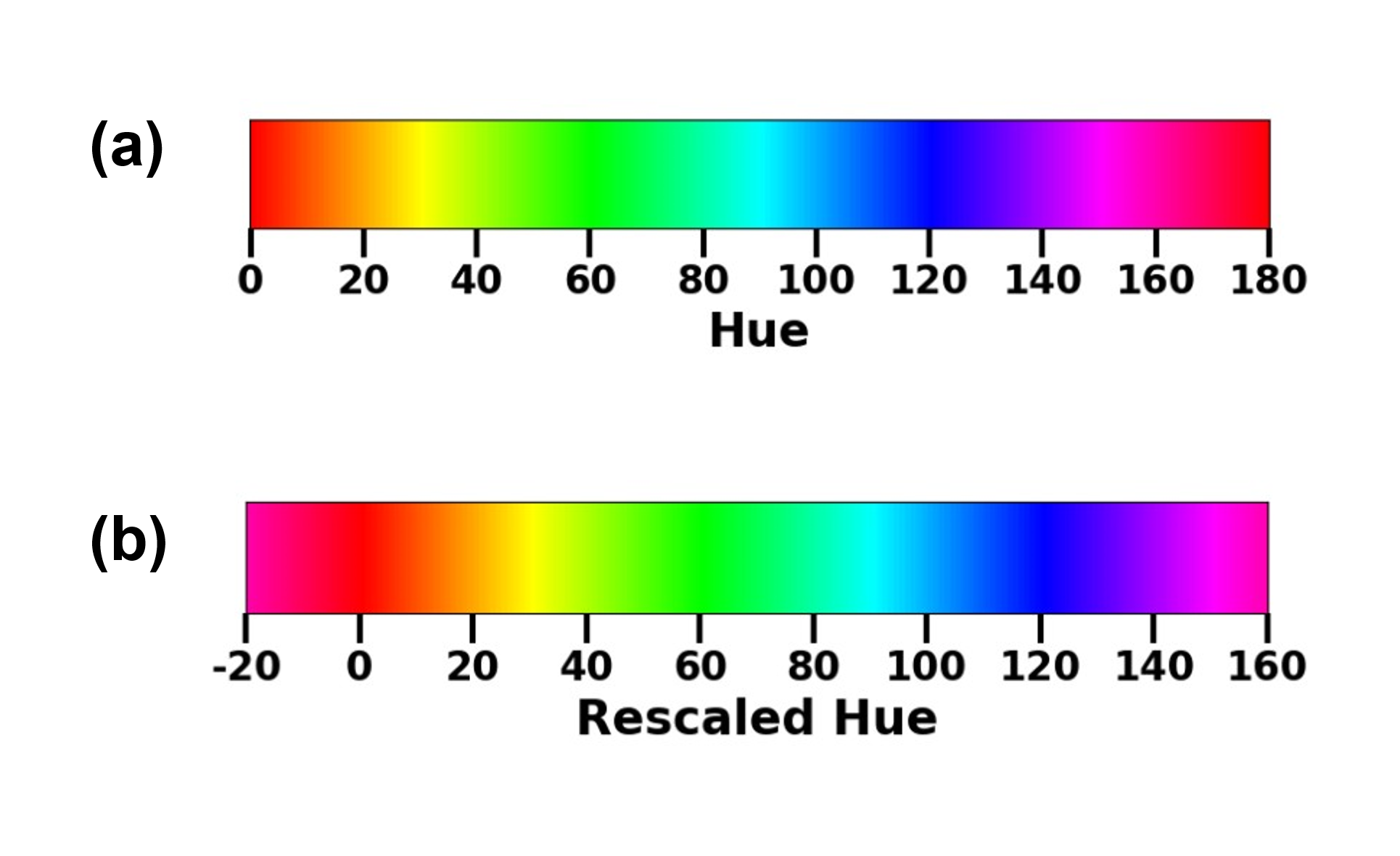

**Figure S6.** Linear representation of the hue color wheel showing (A) the 0-180° hue scale as defined in the OpenCV library and (B) a rescaled version where the red region (160-180°) is shifted to −20 to -1°, producing a continuous red-to-yellow range suitable for hue analysis in colorimetric LAMP reactions.

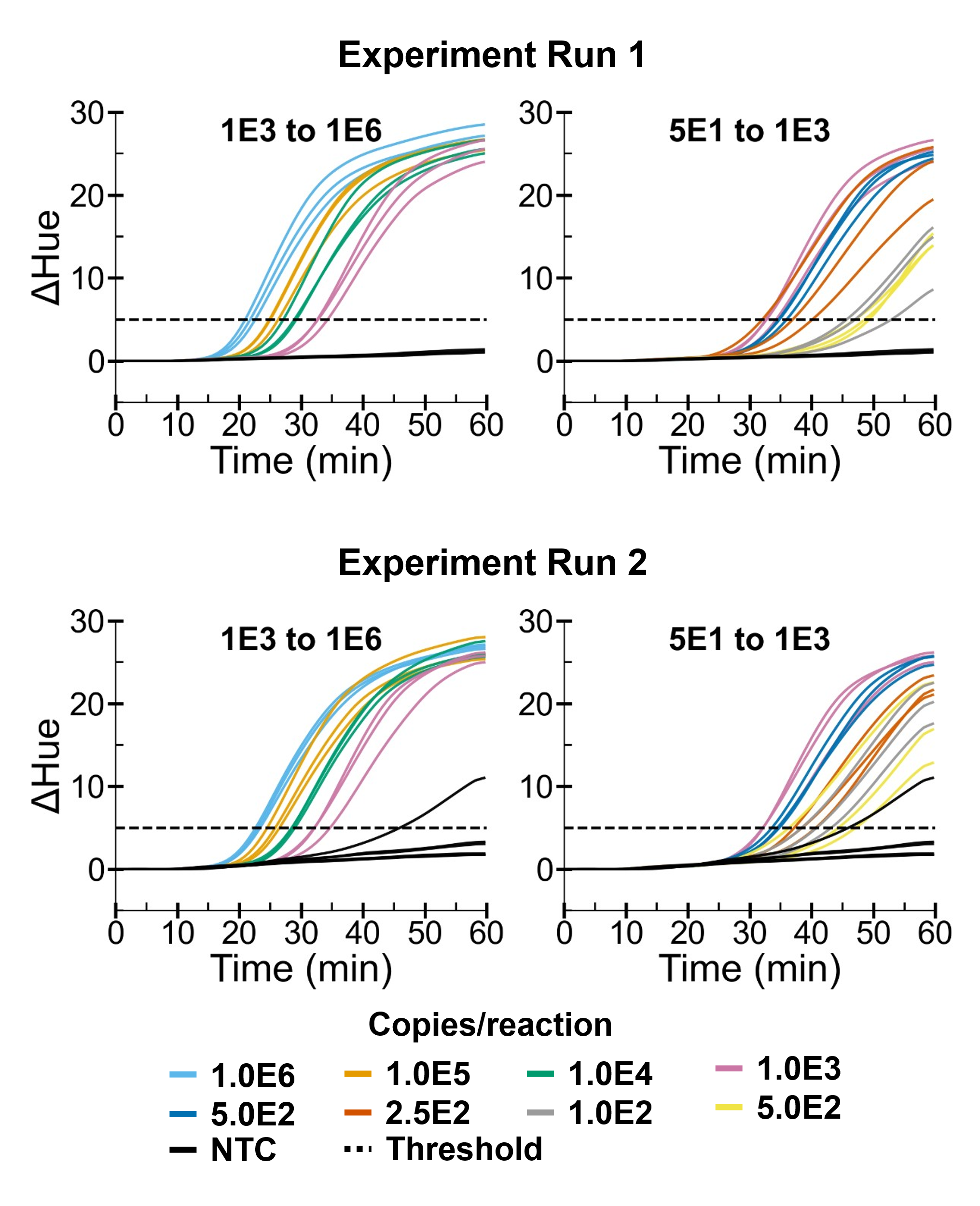

**Figure S7.** ΔHue versus time for color LAMP on uPADs, showing Run 1 (top) and Run 2 (bottom). Each run is split into two panels: Segment 1 (10^3^ to 10^6^ copies/reaction; left) and Segment 2 (50 to 10^3^ copies/reaction; right). Curves are colored by concentration groups with 3 technical triplicates. NTC reactions are included in both segments; in Run 2, one NTC trace crosses the threshold (false positive). The horizontal dashed line indicates the positivity threshold.

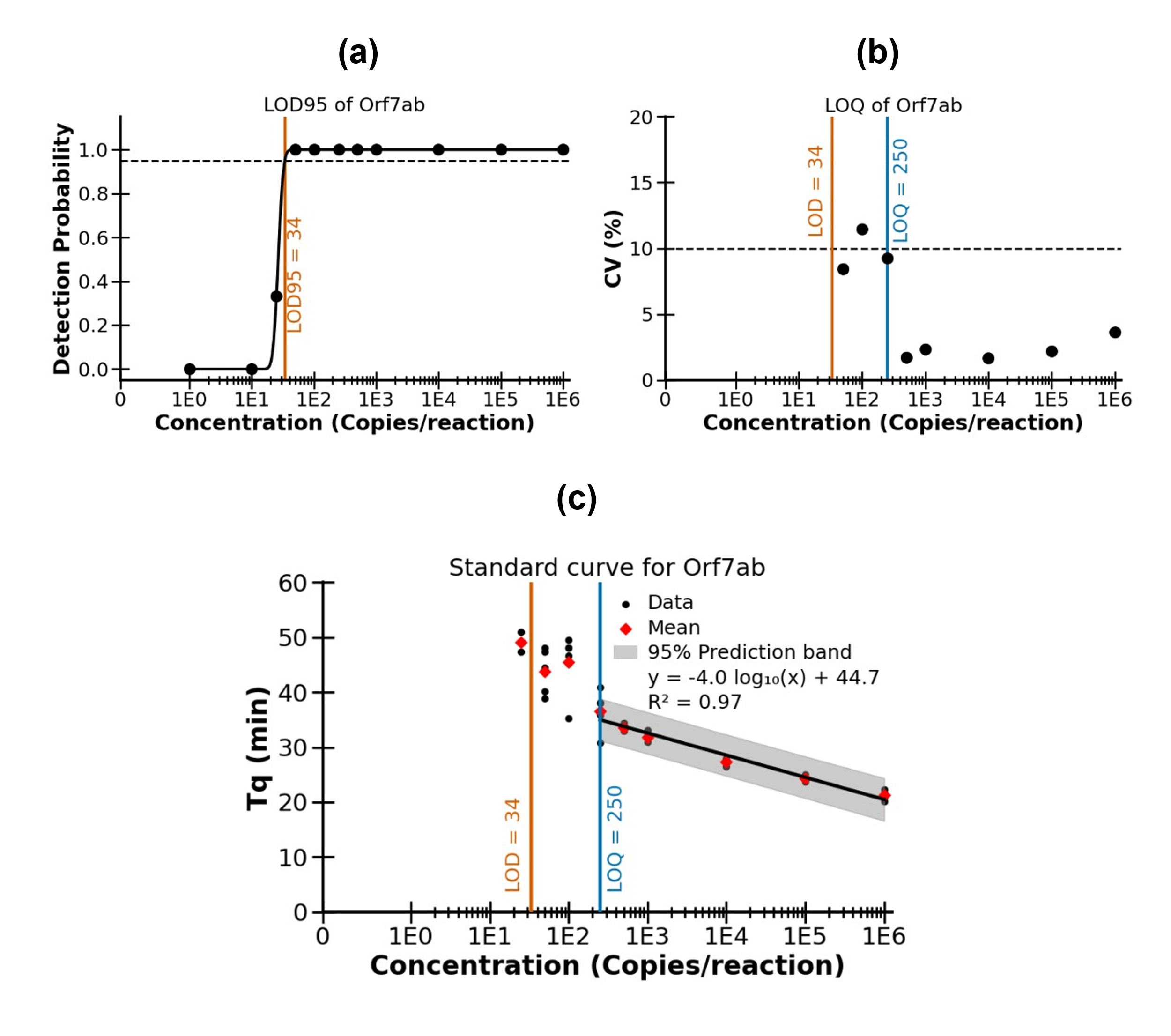

**Figure S8.** Determination of limit of detection (LOD) with 95% detection probability (LOD95) and limit of quantification (LOQ). (a) Probit regression of detection probability versus log₁₀ concentration was used to estimate LOD95 (95% detection probability). (b) LOQ was defined as the lowest concentration at or above LOD95 with CV ≤ 10% across replicates (n = 6 per concentration). (c) Standard curve (Tq vs log_10_ concentration) fitted using data ≥ LOQ; shaded region denotes the 95% prediction interval. Concentrations between LOD95 and LOQ are detectable but not reliably quantifiable.

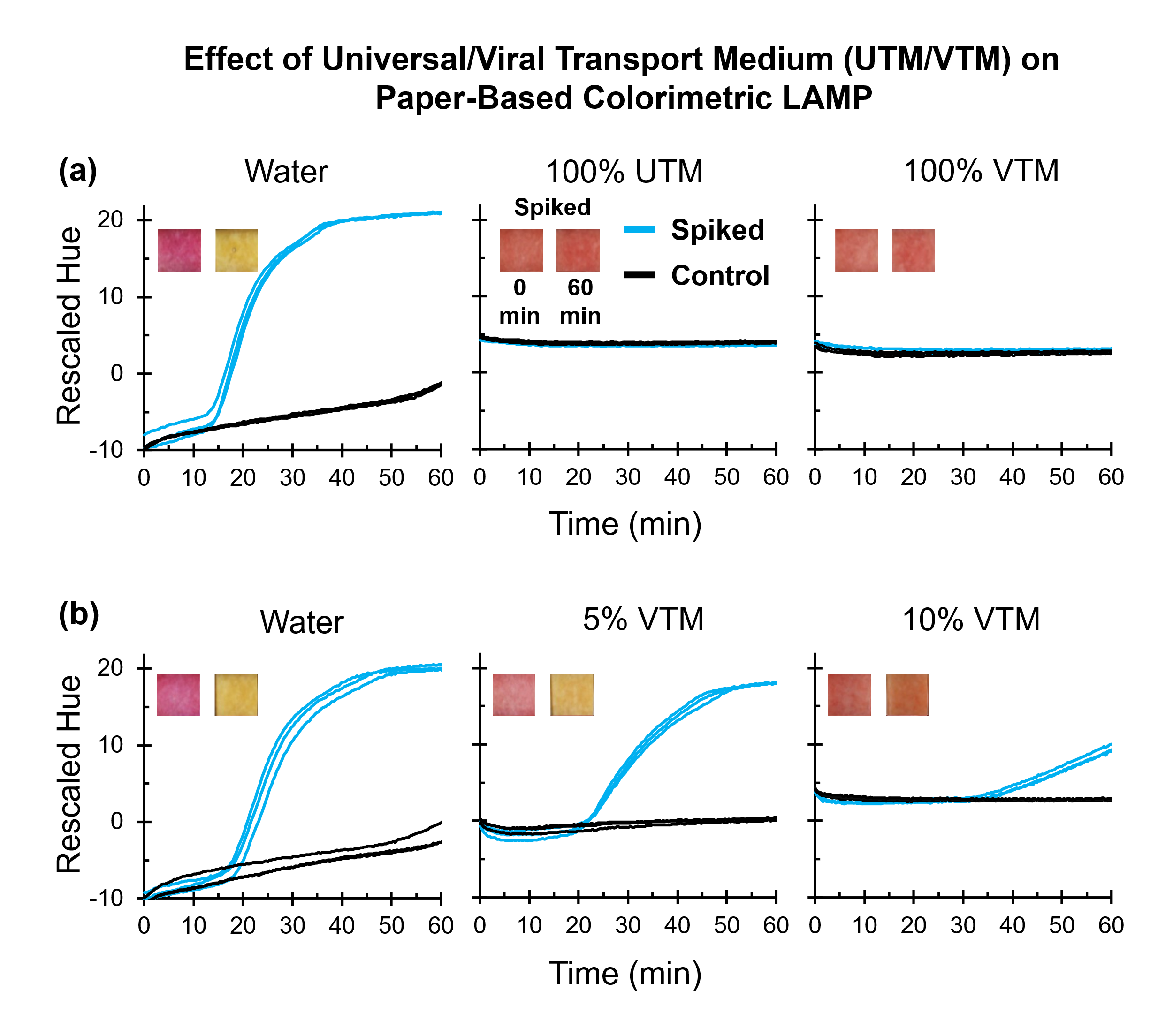

**Figure S9.** Effect of transport media on paper-based colorimetric LAMP. (A) Rescaled hue-time trajectories (see Figure S6 for hue rescaling) for LAMP reactions with 10^5^ copies/reaction synthetic *orf7ab* DNA target spiked (blue, n = 3) or no-template controls (NTC, black, n = 3) prepared in nuclease-free water and in undiluted transport media (100% UTM and 100% VTM). (B) Extension of A, showing rescaled hue–time trajectories for the same spiked and NTC conditions in nuclease-free water and after diluting VTM in nuclease-free water to 5% and 10% (v/v). Inset show representative µPAD images at 0 and 60 min for the spiked sample. Inset image were post-processed in Microsoft PowerPoint by adjusting brightness (-20%) and contrast (+40%) for improved visual representation; however, all image analyses presented were generated from raw image data using custom Python-based software, Amplimetrics™.

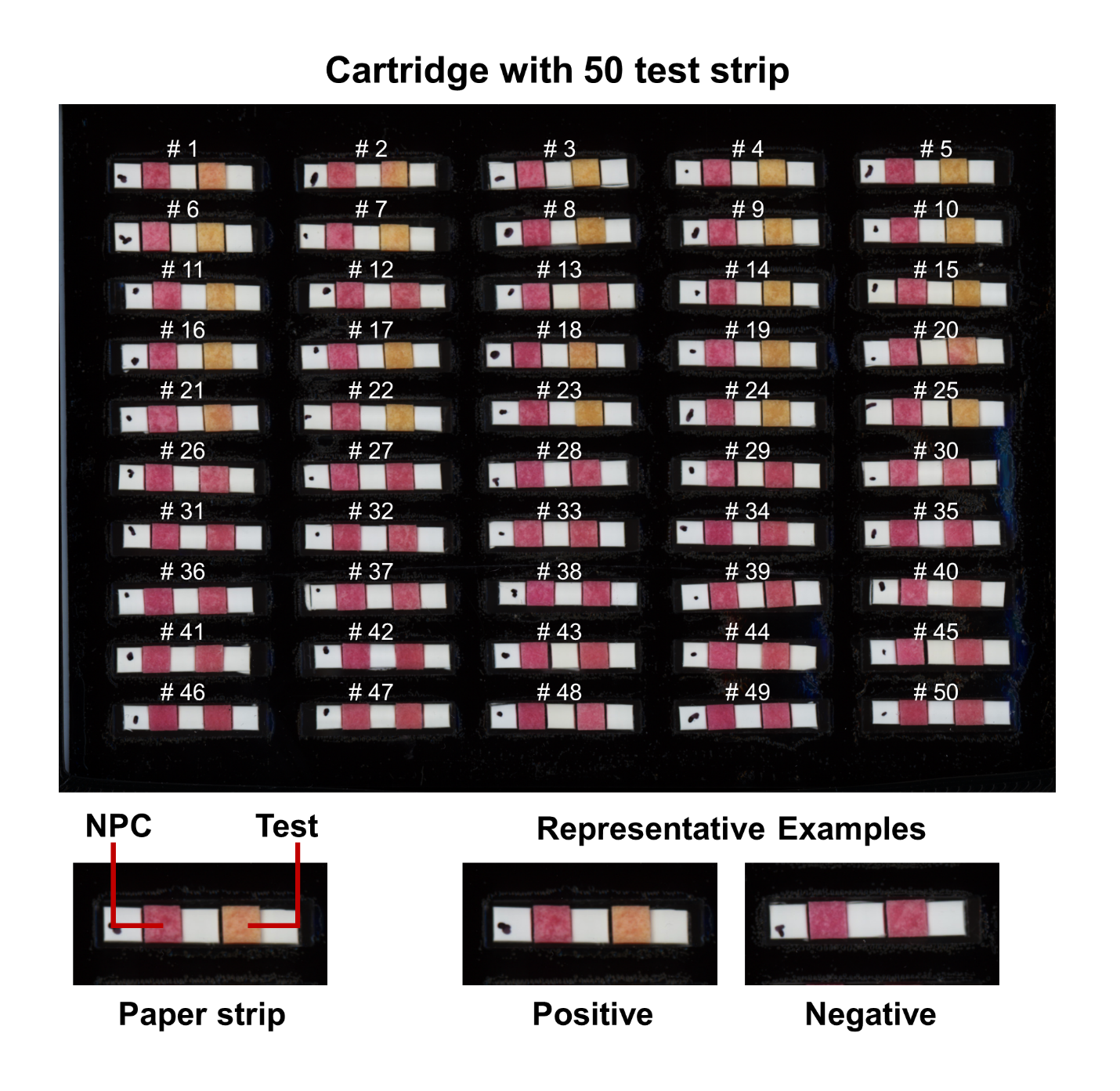

**Figure S10.** Representative full test cartridge containing 50 paper-based RT-LAMP paper strips after 60 minutes incubation at 65 °C. Each paper strip consists of two µPADs: an RT-LAMP test µPAD and a paired no-primer control (NPC) µPAD. A positive test is indicated by a color change of the test µPAD from red to yellow, while the NPC remains unchanged. Negative tests show no color change in either µPAD. An invalid test occurs when the NPC µPADs changes color (not shown). Image for this figure was post-processed in Microsoft PowerPoint by adjusting contrast (+40%) for improved visual representation; however, all image analyses presented were generated from raw image data using custom Python-based software, Amplimetrics™.

### 3. Supplementary Tables

Table S1. Key settings for the VueScan version 9.8.01. software. Settings not listed here should be left at their default values. Please refer to Supplementary Notes 1.2 for detail on the step-by-step operation of the VueScan software.

| **Menu** | **Scanner Settings** |
| --- | --- |
| **Input** | **Options:** Professional  **Tasks:** Scan to file  **Source:** PerfectionV800  **Mode:** Flatbed  **Media:** Color  **Media size:** 8.5 x 11 in.  **Bits per pixel:** Auto  **Scan from preview:** unchecked  **Preview resolution:** 600 dpi  **Scan resolution:** 600 dpi  **Rotation:** either None or left  **Auto Save:** Scan  **Auto repeat:** For first scan choose None, then switch to a 15 seconds for timelapse  **Number of samples:** 2  **Default folder:** Create/select a folder where images will be saved.  **JPG filename:** Leave default (e.g., Scan0001+.jpg). The “0001+.jpg” format is required for filename incrementing during timelapse capture. Do not delete files mid-timelapse, as this will reset numbering. |
| **Crop** | **Crop size:** Manual. Leave all other parameters at default. |
| **Filter** | Do not change any settings. |
| **Color** | **Color Balance:** Auto levels  **Printer color space:** sRGB  Leave all other parameters at default. |
| **Output** | **Default folder:** Same as set in Input so either select here or on in Input section.  **JPEG File:** Check only JPG (do not check other formats)  **JPEG file name:** Leave default (e.g., Scan0001+.jpg)  **JPEG size reduction:** 1  **JPEG multi page:** Off  **JPEG quality:** 90-95%  **Printed size:** Scan size  **Magnification:** 100% |
| **Prefs** | Leave all settings at default. |
| **PREVIEW** | Press Preview to display the scan in the right window. Use the mouse cursor to manually crop the region of interest. |
| **SCAN** | Before scanning, confirm folder and filenames are correct. Filenames must end with “-0001+.jpg” for incrementing to work during timelapse. While the first image is scanning, go to Input and set Auto repeat to 15 s (or chosen interval). To stop timelapse, reset Auto repeat to None. Always check that the first few timelapse images are saved correctly. |

Table S2. BOM of ThermiQuant™ MegaScan (Fixed cost)

| Bill of materials for the ThermiQuant™ MegaScan | | | | | | | |
| --- | --- | --- | --- | --- | --- | --- | --- |
| S.N. | **Parts name** | **Vendor** | **Units** | **Price/**  **Unit** | **Total Price (USD)** | | **Notes** |
| 1 | [Epson Perfection V800 Photo scanner](https://www.amazon.com/Epson-Perfection-V800-Photo-scanner/dp/B00OCEJM9K/ref=sr_1_3?crid=3FCJA51PTTYBX&dib=eyJ2IjoiMSJ9.6SekCOAzIOhxz7g0gk11a9PT3N5nejd2aCwBVsP8PQy2O2UHOp4b9XDAVHQWStH5RC5w_pAyHTWNOE9PcG7ZtHNuO_sz54SnSGsKwhI2--J3gN5kmQnGGKEKzPQ0lb-S_He8u81FEiVPqrZ8F6nVqvHtt0Agu1qhkReH9zk8mK3PG9tfvaNaVvIaqIJJk4h4xaJwdM1tdrNJVosPKoJpD6OMhxLwl2GdUSHEDwHnOb8.K2xHjGfcTVrb2edlP-8-aUo0DrqKPs0eznG16WowvSY&dib_tag=se&keywords=epson+v800+pro+scanner&qid=1756157910&sprefix=epson+v800+pro+scanner%2Caps%2C91&sr=8-3) | Epson | 1 | 1,298 | 1298 | | Can use either V800 or V850. |
| 2 | [Small Nano Rimless Tank (1.1 gallon)](https://www.amazon.com/AWXZOM-Aquarium-1-1Gallon-7-8x5-5x5-9inch-20x14x15cm/dp/B0D3F36CTY/ref=sr_1_17?crid=1G7HG2F401D0K&dib=eyJ2IjoiMSJ9.1knzeLNahx5xWryGogltnRYeJEGlfmagIzNOZpqIvQde6-uoyhsEUGw08waAaNAmpHOKyMjUbpngktBQHHASgK8EBtJbJ69WJa2StVpp8bY9FwfhDWY6j6lDPdf7yxOQwghE2GSmiwuiTtysTDDLlme-0sZQTLOnf_JjbJSOaNsdaNCMkvQrKtiFuJ362ANO2J_dvxbBVZsZ7lA9q7AyFELtW4M1QxbgqpYqp_Gu9GxCLz2S9zRt4RuUDc2dvRCssNU4jllKxdZ92JwBnYxYJsjR1df7IgkgEgZuhwptv9g.IvKT0pxAvhK6LGYEahaSqpZIg7PoYeef3SYZgcfpI68&dib_tag=se&keywords=1.1%2Bgallon%2Bmini%2Bfish%2Btank&qid=1756157973&sprefix=1.1%2Bgallon%2Bmini%2Bfish%2Btank%2Caps%2C99&sr=8-17&th=1) (AWXZOM, ASIN: B0D3F36CTY) | Amazon USA | 1 | 29.99 | 29.99 | | Brand name: AWXZOM and tank external size: 0.8 x 5.5 x 5.9 inch/ 20 x14 x15cm (1.1-gallon capacity) |
| 3 | [Anova Culinary Sous Vide Precision Cooker Nano 3.0, 800 watts](https://www.amazon.com/Anova-Precision-Cooker-Nano-3-0/dp/B0BQ93XGWC/ref=sr_1_1?crid=1ZKVAOE4MG4R8&dib=eyJ2IjoiMSJ9.3Ev3hZ8hS4NJ4Nx_kkaYQNHvoR0BFeGiacyuwI35HfVpXFFu4OwWu9f1zZzV3aEMyQDeeTu9S2lvi_Nb5hKZCBohVdXT5yl6gdioox4i-HiMCskYQ7F90EWK5l08QujvMefhrgQ2-aIzKBkCAzCF_4vJ0bx1hSRmaBI1YxclkRtgi8Z8laI8KEQYB3nVTI-Ac4zFC27RyBUzt6jkXLa_XCqNxfPw3v_GZFngrNpFbvE.mMLIIKNzX6XQqW_zN921GXNxFkC43VbpATUyY4tTvs0&dib_tag=se&keywords=Anova%2Bnano%2Bheater%2Brod&qid=1757527452&sprefix=anova%2Bnano%2Bheater%2Brod%2Caps%2C113&sr=8-1&th=1) | Anova | 1 | 84.11 | 84.11 | | Anova has recently released a cheaper "mini" version that costs USD 40 |
| 4 | [VueScan Software for timelapse imaging](https://www.hamrick.com/purchase-vuescan.html) | Hamrick USA | 1 | 131.21 | 131.21 | | This is price for one time purchase but 1 year subscription or bulk order is one third this price. Please note price may change over time. |
| 5 | 3D Printed Parts | Amazon USA | 1 | 30.00 | 30.00 | | Price estimate based on 1 Kg 3D printed PETG/PC filament cost. |
| Total*  *Cost of Laptop/PC not included as this is generally a shared resource in a lab | | | | | | **1573.30** | |

Table S3. BOM of 160 µPADs cartridge (Consumable cost)

| Consumable Cost (estimated for total of 160 µPAD reaction) | | | | | | |
| --- | --- | --- | --- | --- | --- | --- |
| S.N. | **Parts Name** | **Vendor** | **Units** | **Price/ Unit** | **Total Price (USD)** | **Notes** |
| 1 | [1.5 mm Acrylic for cartridge](https://www.amazon.com/Pieces-Acrylic-Sheet-Transparent-Project/dp/B09MRFHFTR/ref=sr_1_3?crid=3FYCOEH450GSN&dib=eyJ2IjoiMSJ9.ihSLCOsV10Aqr9Z5-Uye32nv9oSz0R-J5qUYbwjWWMQKYl1RgDehqXhzLbU7mufas7CLMOpU2WFHRfz94CUyVGi14zTMnIo5BeQKWOlkLHeVmG_6tsV7LUgD_7xzA6hRAMBOZljm_7sD73SNH3t86W8CtePxxN_u9X_Zn_P7ZMLPCiZ_OFJcvoyUShRyZRQDPykD06s-K0KOe1wCPLy_jfd7g_HYzriw5n3z3lNHrBs.Uh2vt7sTtHesFMpOyfvu4WRSIXqEJfbe9EZxcD4kRqk&dib_tag=se&keywords=1.5%2Bmm%2Bblack%2Bacrylic&qid=1756158730&sprefix=1.5%2Bmm%2Bblack%2Bacrylic%2Caps%2C111&sr=8-3&th=1) (Outus, ASIN: B09MRFHFTR) | Amazon | 0.17 | 4.3 | 0.72 | One 12-inch x 12-inch acrylic panel can be used for fabricating 6 full size Acrylic Cartridge |
| 2 | [Adhesive PCR Plate Seals](https://www.thermofisher.com/order/catalog/product/AB0558) (Catalog number: AB0558) | ThermoFisher Scientific | 0.02 | 188.65 | 3.77 | 2 full PCR tape is required (both sides) and it can be bought at a bulk of 100 units. |
| 3 | [LAMP assay for µPADs](https://www.sciencedirect.com/science/article/pii/S0956566325005640) | Multiple vendors | 160 | 0.74 | 118.4 | Cost estimate from Ahmed, et al. (2025).^1^ |
| Total*  Total cost for running 160 reactions in a run | | | | | **122.89** | |

***Synthetic DNA, LAMP Primers, and qPCR probes and primers***

dPCR was used for absolute quantification, and LAMP was used for validation of the µPAD-based assay, as described in the main Methods section. This study builds on a previously reported design of µPAD-based LAMP reactions targeting the orf7ab region of SARS-CoV-2^2,3^.

**Table S4. *Orf7ab* synthetic DNA sequence**

*orf7ab* Synthetic DNA sequence (5’-3’)

ATGAAAATTATTCTTTTCTTGGCACTGATAACACTCGCTACTTGTGAGCTTTATCACTACCAAGAGTGTGTTAGAGGTACAACAGTACTTTTAAAAGAACCTTGCTCTTCTGGAACATACGAGGGCAATTCACCATTTCATCCTCTAGCTGATAACAAATTTGCACTGACTTGCTTTAGCACTCAATTTGCTTTTGCTTGTCCTGACGGCGTAAAACACGTCTATCAGTTACGTGCCAGATCAGTTTCACCTAAACTGTTCATCAGACAAGAGGAAGTTCAAGAACTTTACTCTCCAATTTTTCTTATTGTTGCGGCAATAGTGTTTATAACACTTTGCTTCACACTCAAAAGAAAGACAGAATGATTGAACTTTCATTAATTGACTTCTATTTGTGCTTTTTAGCCTTTCTGCTATTCCTTGTTTTAATTATGCTTATTATCTTTTGGTTCTCACTTGAACTGCAAGATCATAATGAAACTTGTCACGCCTAA

**Table S5. dPCR primers and probes**

| **qPCR Primers** | **Sequence** | **Vendor** |
| --- | --- | --- |
| orf7ab_PCR2_FWD | GAGGGCAATTCACCATTTCATC | Life Science Tech |
| orf7ab_PCR2_REV | AAACTGATCTGGCACGTAACT | Life Science Tech |
| Probe | /56-FAM/TT TGC TTG T/ZEN/C CTG ACG GCG TAA A/3IABkFQ/ | IDT |

**Table S6. LAMP primers**

| Primers | Sequence | Vendor |
| --- | --- | --- |
| SC2.orf7ab.1_F3 | CGGCGTAAAACACGTCTA | IDT |
| SC2.orf7ab.1_B3 | GCTAAAAAGCACAAATAGAAG TC | IDT |
| SC2.orf7ab.1_FIP | GGAGAGTAAAGTTCTTGAACTT CCTAGTTACGTGCCAGATCAG | IDT |
| SC2.orf7ab.1_BIP | TGCGGCAATAGTGTTTATAAC ACTATGAAAGTTCAATCATTCT GTCT | IDT |
| SC2.orf7ab.1_LF | TGTCTGATGAACAGTTTAGGT GAAA | IDT |
| SC2.orf7ab.1_LB | TTGCTTCACACTCAAAAGAA | IDT |

### 4. Supplementary movie 1: Amplimetrics software user guide.
