## Supplementary Software for "ThermiQuant™ MegaScan: High-throughput isothermal reactor with quantitative colorimetric readout for paper-based nucleic acid amplification tests": data_analysis.py

```
import os
import csv
import pandas as pd
import numpy as np
from math import ceil
from PyQt5.QtWidgets import (
    QWidget, QVBoxLayout, QPushButton, QHBoxLayout, QLabel,
    QComboBox, QLineEdit, QFileDialog, QTextEdit,QGroupBox
)
from PyQt5.QtGui import QStandardItemModel, QStandardItem, QFont
from PyQt5.QtCore import Qt
from matplotlib.lines import Line2D
from scipy.ndimage import gaussian_filter1d
from scipy.optimize import curve_fit
from scipy.special import wofz
from scipy.stats import t as t_dist

def voigt(x, amplitude, center, sigma, gamma, offset):
    z = ((x - center) + 1j * gamma) / (sigma * np.sqrt(2))
    return amplitude * np.real(wofz(z)) / (sigma * np.sqrt(2 * np.pi)) + offset

def voigt_fit(x, amplitude, center, sigma, gamma):
    """
    Fit function for the derivative that enforces f(0)=0.
    This is achieved by subtracting the function value at 0.
    """
    z = ((x - center) + 1j * gamma) / (sigma * np.sqrt(2))
    z0 = ((0 - center) + 1j * gamma) / (sigma * np.sqrt(2))
    return amplitude * (np.real(wofz(z))/(sigma*np.sqrt(2*np.pi)) - np.real(wofz(z0))/(sigma*np.sqrt(2*np.pi)))

def sigmoid_func(x, L, k, x0):
    """
    Sigmoid function that satisfies F(0)=0 by subtracting the offset.
    F(x) = L/(1+exp(-k*(x-x0))) - L/(1+exp(k*x0))
    
    Here:
      - L is the amplitude,
      - k is the steepness,
      - x0 is the inflection point.
    """
    return L / (1 + np.exp(-k * (x - x0))) - L / (1 + np.exp(k * x0))

class DataAnalysisWidget(QWidget):
    def __init__(self, display_widget, parent=None):
        super().__init__(parent)
        self.display_widget = display_widget
        # Data storage.
        self.df = None               # Raw CSV data.
        self.df_smoothed = None      # Smoothed (and rebased) data.
        self.df_corrected = None     # Baseline-corrected data.
        self.df_to_plot = None       # Processed data to be plotted.
        self.threshold = None
        self.global_ignore_time = None
        self.global_baseline_end_time = None
        self.common_sensor_ylim = None
        self.common_deriv_ylim = None
        self.inBatchMode = False
        self.current_batch = 0
        self.total_batches = 0
        self.tq_df = None
        self.input_filename = None
        self.experiment_duration = None  # Holds the experiment duration
        self.initUI()

    def initUI(self):
        self.setFont(QFont("Arial", 11))
        style_sheet = """
        QPushButton {
            background-color: #555555;
            color: white;
            border: 1px solid #444444;
            border-radius: 4px;
            padding: 5px 10px;
            font: 11pt "Arial";
        }
        QPushButton:disabled {
            background-color: #cccccc;
            color: #666666;
            border: 1px solid #aaaaaa;
        }
        QLineEdit, QComboBox {
            background-color: #ffffff;
            color: #333333;
            border: 1px solid #888888;
            border-radius: 4px;
            padding: 3px 5px;
            font: 10pt "Arial";
        }
        QLabel {
            font: 10pt "Arial";
            color: #333333;
        }
        QTabBar::tab {
            background-color: #e0e0e0;
            border: 1px solid #aaa;
            border-radius: 5px;
            padding: 8px;
        }
        QTabBar::tab:selected {
            background-color: #d0d0d0;
            border: 1px solid #888;
        }
        """
        self.setStyleSheet(style_sheet)
        main_layout = QVBoxLayout()

        # Load CSV button.
        self.load_csv_btn = QPushButton("Load CSV")
        self.load_csv_btn.clicked.connect(self.load_csv)
        main_layout.addWidget(self.load_csv_btn)

        # Baseline selection description and dropdowns.
        desc_label = QLabel("Select Start and End Baseline Value")
        desc_label.setFont(QFont("Arial", 12, QFont.Bold))
        main_layout.addWidget(desc_label)
        baseline_layout = QHBoxLayout()
        baseline_layout.addWidget(QLabel("Start:"))
        self.start_combo = QComboBox()
        baseline_layout.addWidget(self.start_combo)
        baseline_layout.addWidget(QLabel("End:"))
        self.stop_combo = QComboBox()
        baseline_layout.addWidget(self.stop_combo)
        main_layout.addLayout(baseline_layout)

        # "Smooth Normalize" button.
        self.smooth_btn = QPushButton("Smooth Normalize")
        self.smooth_btn.clicked.connect(self.update_baseline_plot)
        main_layout.addWidget(self.smooth_btn)

        # Experiment Duration widget (shortened).
        duration_layout = QHBoxLayout()
        duration_layout.addWidget(QLabel("Exp Duration:"))
        self.duration_edit = QLineEdit()
        self.duration_edit.setPlaceholderText("Enter experiment duration")
        duration_layout.addWidget(self.duration_edit)
        self.apply_duration_btn = QPushButton("Apply")
        self.apply_duration_btn.clicked.connect(self.apply_experiment_duration)
        duration_layout.addWidget(self.apply_duration_btn)
        main_layout.addLayout(duration_layout)

        # Threshold entry.
        thresh_layout = QHBoxLayout()
        thresh_layout.addWidget(QLabel("Adjust Threshold:"))
        self.thresh_edit = QLineEdit()
        self.thresh_edit.setPlaceholderText("Enter threshold value")
        thresh_layout.addWidget(self.thresh_edit)
        self.apply_thresh_btn = QPushButton("Apply")
        self.apply_thresh_btn.clicked.connect(self.apply_threshold)
        thresh_layout.addWidget(self.apply_thresh_btn)
        main_layout.addLayout(thresh_layout)
       
        # Process Derivative button.
        self.process_deriv_btn = QPushButton("Process Derivative")
        self.process_deriv_btn.clicked.connect(self.process_derivative)
        main_layout.addWidget(self.process_deriv_btn)

        # Batch Navigation container.
        self.nav_widget = QWidget()
        nav_layout = QHBoxLayout(self.nav_widget)
        self.prev_btn = QPushButton("Previous")
        self.prev_btn.clicked.connect(self.prev_batch)
        nav_layout.addWidget(self.prev_btn)
        self.batch_label = QLabel("Batch 0/0")
        nav_layout.addWidget(self.batch_label, alignment=Qt.AlignCenter)
        self.next_btn = QPushButton("Next")
        self.next_btn.clicked.connect(self.next_batch)
        nav_layout.addWidget(self.next_btn)
        main_layout.addWidget(self.nav_widget)

        # Compute & Save Tq buttons.
        tq_layout = QHBoxLayout()
        self.compute_tq_btn = QPushButton("Compute Tq")
        self.compute_tq_btn.clicked.connect(self.compute_tq)
        tq_layout.addWidget(self.compute_tq_btn)
        self.save_tq_btn = QPushButton("Save Tq Data")
        self.save_tq_btn.clicked.connect(self.save_tq_data)
        tq_layout.addWidget(self.save_tq_btn)
        main_layout.addLayout(tq_layout)

        # Save Processed Data button.
        self.save_processed_btn = QPushButton("Save Processed Data")
        self.save_processed_btn.clicked.connect(self.save_processed_data)
        main_layout.addWidget(self.save_processed_btn)

        # Log text widget for status messages.
        self.log_text = QTextEdit()
        self.log_text.setReadOnly(True)
        self.log_text.setMinimumHeight(100)
        main_layout.addWidget(self.log_text)

        main_layout.addStretch(1)
        self.setLayout(main_layout)

        # Initially disable controls and hide the navigation widget.
        self.smooth_btn.setEnabled(False)
        self.start_combo.setEnabled(False)
        self.stop_combo.setEnabled(False)
        self.duration_edit.setEnabled(False)
        self.apply_duration_btn.setEnabled(False)
        self.thresh_edit.setEnabled(False)
        self.apply_thresh_btn.setEnabled(False)
        self.process_deriv_btn.setEnabled(False)
        self.compute_tq_btn.setEnabled(False)
        self.save_tq_btn.setEnabled(False)
        self.save_processed_btn.setEnabled(False)
        self.nav_widget.hide()

    def log_message(self, message):
        self.log_text.append(message)
        print(message)

    def load_csv(self):
        options = QFileDialog.Options()
        file_name, _ = QFileDialog.getOpenFileName(self, "Select CSV File", "", "CSV Files (*.csv)", options=options)
        self.log_message(f"File name returned: {file_name}")
        if not file_name:
            self.log_message("No file selected.")
            return
        try:
            self.df = pd.read_csv(file_name)
            self.log_message(f"DataFrame head:\n{self.df.head()}")
            if self.df is None or self.df.shape[1] < 2:
                raise ValueError("CSV must have at least two columns.")
            self.input_filename = file_name
            # Enable controls.
            self.start_combo.setEnabled(True)
            self.stop_combo.setEnabled(True)
            self.smooth_btn.setEnabled(True)
            self.duration_edit.setEnabled(True)
            self.apply_duration_btn.setEnabled(True)
            self.thresh_edit.setEnabled(True)
            self.apply_thresh_btn.setEnabled(True)
            # Process the CSV.
            self.smooth_normalize()
            # Set default experiment duration as the full length.
            time_col = self.df_smoothed.columns[0]
            max_time = self.df_smoothed[time_col].values[-1]
            self.experiment_duration = max_time
            self.duration_edit.setText(str(max_time))
            # Update table with processed data (with metadata).
            self.populate_table(self.get_processed_data_with_metadata())
            self.inBatchMode = False
            self.process_deriv_btn.setEnabled(False)
            self.compute_tq_btn.setEnabled(False)
            self.save_tq_btn.setEnabled(False)
            self.save_processed_btn.setEnabled(True)
            self.nav_widget.hide()
            self.log_message("CSV loaded and processed successfully.")
        except Exception as e:
            self.log_message(f"Error loading CSV: {e}")

    def smooth_normalize(self):
        if self.df is None:
            self.log_message("No CSV data loaded.")
            return
        time_col = self.df.columns[0]
        df_processed = self.df.copy()
        df_processed.reset_index(drop=True, inplace=True)
        t = df_processed[time_col].values.astype(float)
        t = t - t[0]  # Rebase time.
        df_processed[time_col] = t
        for col in df_processed.columns[1:]:
            series_ma = df_processed[col].rolling(window=5, center=True, min_periods=1).mean()
            series_smoothed = gaussian_filter1d(series_ma.values, sigma=2, mode='reflect')
            df_processed[col] = series_smoothed

        sensor_columns = df_processed.columns[1:]
        ignore_times = []
        for col in sensor_columns:
            y = df_processed[col].values
            dy = np.gradient(y, t)
            threshold_val = np.percentile(np.abs(dy), 90) * 0.5
            ignore_idx = np.argmax(np.abs(dy) < threshold_val)
            ignore_times.append(t[ignore_idx])
        global_ignore_time = int(round(min(max(np.median(ignore_times), 3), 10)))
        self.global_ignore_time = global_ignore_time

        baseline_times = []
        window_size = 5
        for col in sensor_columns:
            y = df_processed[col].values
            start_idx = np.argmax(t >= global_ignore_time)
            rolling_mean = pd.Series(y[start_idx:]).rolling(window=window_size, center=True, min_periods=1).mean()
            rolling_std = pd.Series(y[start_idx:]).rolling(window=window_size, center=True, min_periods=1).std()
            lower_bound = rolling_mean - 1.5 * rolling_std
            upper_bound = rolling_mean + 1.5 * rolling_std
            in_baseline_range = (y[start_idx:] >= lower_bound) & (y[start_idx:] <= upper_bound)
            if np.any(~in_baseline_range):
                baseline_end_idx = start_idx + np.where(~in_baseline_range)[0][0]
            else:
                baseline_end_idx = start_idx + 10
            baseline_times.append(t[baseline_end_idx])
        baseline_duration = int(round(np.median(baseline_times) - global_ignore_time))
        baseline_duration = max(5, min(baseline_duration, 15))
        global_baseline_end_time = global_ignore_time + baseline_duration
        self.global_baseline_end_time = global_baseline_end_time

        self.start_combo.blockSignals(True)
        self.start_combo.clear()
        for i in range(3, 11):
            self.start_combo.addItem(str(i))
        self.start_combo.setCurrentText(str(global_ignore_time))
        self.stop_combo.blockSignals(True)
        self.stop_combo.clear()
        for j in range(global_ignore_time + 1, global_ignore_time + 17):
            self.stop_combo.addItem(str(j))
        self.stop_combo.setCurrentText(str(global_baseline_end_time))
        self.start_combo.blockSignals(False)
        self.stop_combo.blockSignals(False)

        df_corrected = df_processed.copy()
        sigma_gaussian = 2
        for col in sensor_columns:
            y = df_processed[col].values
            baseline_mask = (t >= global_ignore_time) & (t <= global_baseline_end_time)
            baseline_values = y[baseline_mask]
            smoothed_baseline = gaussian_filter1d(baseline_values, sigma=sigma_gaussian, mode='reflect')
            full_smoothed_baseline = np.interp(t, t[baseline_mask], smoothed_baseline)
            corrected = y - full_smoothed_baseline
            corrected[t < global_ignore_time] = 0
            corrected = pd.Series(corrected).rolling(window=5, center=True, min_periods=1).mean().values
            corrected = np.maximum(corrected, 0)
            df_corrected[col] = corrected

        self.df_smoothed = df_processed
        self.df_corrected = df_corrected
        self.df_to_plot = df_corrected

        self.plot_two_subplots(t, df_processed, df_corrected, global_ignore_time, global_baseline_end_time, apply_threshold=False)
        self.log_message("Data smoothing and baseline correction applied.")

    def apply_experiment_duration(self):
        if self.df_smoothed is None:
            self.log_message("No data loaded to apply experiment duration.")
            return
        try:
            duration = float(self.duration_edit.text())
        except ValueError:
            self.log_message("Please enter a valid numeric experiment duration.")
            return

        self.experiment_duration = duration
        time_col = self.df_smoothed.columns[0]
        mask = self.df_smoothed[time_col] <= duration
        self.df_smoothed = self.df_smoothed.loc[mask].reset_index(drop=True)
        self.df_corrected = self.df_corrected.loc[mask].reset_index(drop=True)
        self.df_to_plot = self.df_to_plot.loc[mask].reset_index(drop=True)
        t = self.df_smoothed[time_col].values

        self.plot_two_subplots(t, self.df_smoothed, self.df_corrected,
                            self.global_ignore_time,
                            self.global_baseline_end_time,
                            apply_threshold=(self.threshold is not None))

        self.populate_table(self.get_processed_data_with_metadata())
        self.log_message(f"Experiment duration set to {duration}. Data truncated accordingly.")

        self.setup_threshold_method_ui()

    def setup_threshold_method_ui(self):
        from PyQt5.QtWidgets import QGroupBox, QRadioButton, QButtonGroup, QGridLayout

        if hasattr(self, 'threshold_group'):
            self.threshold_group.setParent(None)

        self.threshold_group = QGroupBox("Threshold Method")
        layout = QVBoxLayout()

        self.manual_radio = QRadioButton("Manual Threshold")
        self.manual_radio.setChecked(True)
        self.suggest_radio = QRadioButton("Suggest Threshold")
        self.radio_group = QButtonGroup()
        self.radio_group.addButton(self.manual_radio)
        self.radio_group.addButton(self.suggest_radio)

        layout.addWidget(self.manual_radio)
        layout.addWidget(self.suggest_radio)
        self.threshold_group.setLayout(layout)
        self.layout().insertWidget(5, self.threshold_group)

        if hasattr(self, 'control_panel'):
            self.control_panel.setParent(None)
        if hasattr(self, 'suggest_box'):
            self.suggest_box.setParent(None)

        self.suggest_radio.toggled.connect(self.setup_control_buttons_ui)

    def setup_control_buttons_ui(self):
        from PyQt5.QtWidgets import QGridLayout, QPushButton, QHBoxLayout

        if hasattr(self, 'control_panel'):
            self.control_panel.setParent(None)

        if not self.suggest_radio.isChecked():
            return

        self.control_panel = QGroupBox("Select Control Labels")
        layout = QVBoxLayout()
        self.control_buttons_layout = QGridLayout()
        self.control_buttons = []
        self.control_page = 0

        labels = list(self.df_corrected.columns[1:])
        self.all_control_labels = labels
        self.selected_controls = set()
        self.control_button_refs = {}

        self.color_palette = ['red', 'green', 'blue', 'orange', 'purple', 'brown', 'magenta', 'cyan', 'lime', 'navy']
        self.control_colors = {}

        def make_button(label):
            btn = QPushButton(label)
            btn.setCheckable(True)
            btn.clicked.connect(lambda: self.toggle_control_label(label))
            return btn

        for label in labels:
            btn = make_button(label)
            self.control_button_refs[label] = btn
            self.control_buttons.append(btn)

        nav_layout = QHBoxLayout()
        self.prev_btn = QPushButton("Prev")
        self.next_btn = QPushButton("Next")
        self.prev_btn.clicked.connect(self.prev_control_page)
        self.next_btn.clicked.connect(self.next_control_page)
        nav_layout.addWidget(self.prev_btn)
        nav_layout.addWidget(self.next_btn)

        layout.addLayout(self.control_buttons_layout)
        layout.addLayout(nav_layout)
        self.control_panel.setLayout(layout)
        self.layout().insertWidget(6, self.control_panel)

        self.update_control_buttons_page()

        # Remove previous suggestion box if it exists
        if hasattr(self, 'suggest_box'):
            self.suggest_box.setParent(None)

        self.suggest_box = QGroupBox("Threshold Suggestion Settings")
        sugg_layout = QHBoxLayout()
        self.multiplier_edit = QLineEdit("5")
        self.multiplier_edit.setPlaceholderText("Std Dev Multiplier")
        self.suggest_thresh_btn = QPushButton("Suggest Threshold")
        self.suggest_thresh_btn.clicked.connect(self.suggest_threshold)

        sugg_layout.addWidget(QLabel("Multiplier:"))
        sugg_layout.addWidget(self.multiplier_edit)
        sugg_layout.addWidget(self.suggest_thresh_btn)
        self.suggest_box.setLayout(sugg_layout)

        self.layout().insertWidget(7, self.suggest_box)


    def update_control_buttons_page(self):
        for i in reversed(range(self.control_buttons_layout.count())):
            self.control_buttons_layout.itemAt(i).widget().setParent(None)

        start = self.control_page * 6
        end = start + 6
        subset = self.control_buttons[start:end]

        for i, btn in enumerate(subset):
            row, col = divmod(i, 3)
            self.control_buttons_layout.addWidget(btn, row, col)

    def prev_control_page(self):
        if self.control_page > 0:
            self.control_page -= 1
            self.update_control_buttons_page()

    def next_control_page(self):
        max_page = (len(self.control_buttons) - 1) // 6
        if self.control_page < max_page:
            self.control_page += 1
            self.update_control_buttons_page()

    def toggle_control_label(self, label):
        btn = self.control_button_refs[label]
        if label in self.selected_controls:
            self.selected_controls.remove(label)
            btn.setStyleSheet("")
            self.control_colors.pop(label, None)
        else:
            self.selected_controls.add(label)
            color = self.color_palette[len(self.control_colors) % len(self.color_palette)]
            self.control_colors[label] = color
            btn.setStyleSheet(f"background-color: {color}; color: white;")

        self.update_control_plot()

    def update_control_plot(self):
        if self.df_corrected is None:
            return

        time_col = self.df_corrected.columns[0]
        t = self.df_corrected[time_col].values
        ax = self.display_widget.canvas.fig.axes[1]  # processed data plot
        ax.clear()

        for col in self.df_corrected.columns[1:]:
            y = self.df_corrected[col].values
            if col in self.selected_controls:
                color = self.control_colors.get(col, 'red')
                ax.plot(t, y, linestyle=':', color=color, label=col)
            else:
                ax.plot(t, y, linestyle='-', color='black', alpha=0.3)

        ax.set_title("Processed Data")
        ax.set_xlabel("Time (min)")
        ax.set_ylabel("Sensor Value")
        ax.legend(loc="upper left", fontsize=8, frameon=False)
        self.display_widget.canvas.draw()
    
    
    def suggest_threshold(self):
        if self.df_corrected is None:
            self.log_message("No data to compute threshold from.")
            return

        try:
            duration = float(self.duration_edit.text())
            multiplier = float(self.multiplier_edit.text())
        except ValueError:
            self.log_message("Invalid numeric values for duration or multiplier.")
            return

        time_col = self.df_corrected.columns[0]
        t = self.df_corrected[time_col].values
        control_labels = list(self.selected_controls)

        if not control_labels:
            self.log_message("No controls selected.")
            return

        values_at_duration = []
        for label in control_labels:
            y = self.df_corrected[label].values
            y_interp = np.interp(duration, t, y)
            values_at_duration.append(y_interp)

        values = np.array(values_at_duration)
        self.log_message(f"Raw control values at experiment duration: {values.tolist()}")

        # ----------- basic‑dropout (n<=4)  or  IQR (>4) -----------
        n = len(values)
        if n < 2:
            self.log_message("Not enough control values to filter outliers.")
            filtered = values
        elif n <= 4:
            mu  = np.mean(values)
            sd  = np.std(values, ddof=1)
            if sd == 0:
                filtered = values                    # all identical
            else:
                # find the point farthest from the mean
                idx_max = np.argmax(np.abs(values - mu))
                z_max   = np.abs(values[idx_max] - mu) / sd
                if z_max > 1:                      # basic “>2 SD” rule
                    self.log_message(f"Drop‑out: removed {values[idx_max]:.4f} (z={z_max:.2f})")
                    filtered = np.delete(values, idx_max)
                else:
                    filtered = values
        else:   # n >= 5 → IQR fences
            q1 = np.percentile(values, 25)
            q3 = np.percentile(values, 75)
            iqr = q3 - q1
            lo, hi = q1 - 1.5 * iqr, q3 + 1.5 * iqr
            mask = (values >= lo) & (values <= hi)
            filtered = values[mask]
            dropped  = values[~mask]
            if dropped.size:
                self.log_message(f"IQR: removed {dropped.tolist()}")

        new_mean = np.mean(filtered)
        new_std = np.std(filtered)
        suggested_threshold = new_mean + multiplier * new_std

        self.thresh_edit.setText(f"{suggested_threshold:.2f}")
        self.log_message(f"Suggested threshold: {suggested_threshold:.2f} (mean={new_mean:.2f}, std={new_std:.2f})")


    def update_baseline_plot(self):
        if self.df_smoothed is None:
            return
        time_col = self.df_smoothed.columns[0]
        t = self.df_smoothed[time_col].values
        try:
            new_ignore_time = float(self.start_combo.currentText())
            new_baseline_end_time = float(self.stop_combo.currentText())
        except ValueError:
            return

        self.global_ignore_time = new_ignore_time
        self.global_baseline_end_time = new_baseline_end_time

        sensor_columns = self.df_smoothed.columns[1:]
        df_corrected_new = self.df_smoothed.copy()
        sigma_gaussian = 2
        for col in sensor_columns:
            y = self.df_smoothed[col].values
            baseline_mask = (t >= new_ignore_time) & (t <= new_baseline_end_time)
            if np.sum(baseline_mask) < 2:
                corrected = y.copy()
            else:
                baseline_values = y[baseline_mask]
                smoothed_baseline = gaussian_filter1d(baseline_values, sigma=sigma_gaussian, mode='reflect')
                full_smoothed_baseline = np.interp(t, t[baseline_mask], smoothed_baseline)
                corrected = y - full_smoothed_baseline
                corrected[t < new_ignore_time] = 0
                corrected = pd.Series(corrected).rolling(window=5, center=True, min_periods=1).mean().values
                corrected = np.maximum(corrected, 0)
            df_corrected_new[col] = corrected
        self.df_corrected = df_corrected_new
        self.df_to_plot = df_corrected_new
        self.plot_two_subplots(t, self.df_smoothed, df_corrected_new, new_ignore_time, new_baseline_end_time, apply_threshold=True)
        self.populate_table(self.get_processed_data_with_metadata())
        self.log_message("Baseline parameters updated.")

    def plot_two_subplots(self, t, df_raw, df_corrected, ignore_time, baseline_end_time, apply_threshold=False):
        self.display_widget.canvas.fig.clf()
        axes = self.display_widget.canvas.fig.subplots(1, 2)
        ax_left, ax_right = axes

        if t is not None:
            sensor_columns = df_raw.columns[1:]
            for col in sensor_columns:
                ax_left.plot(t, df_raw[col].values, color='black', alpha=0.8)
            ax_left.axvspan(t[0], ignore_time, color='gray', alpha=0.3, label="Ignored Region")
            ax_left.axvspan(ignore_time, baseline_end_time, color='blue', alpha=0.3, label="Baseline")
            ax_left.set_title("Raw Data")
            ax_left.set_xlabel("Time (min)")
            ax_left.set_ylabel("Sensor Value")
            ax_left.legend(loc="upper left", frameon=False)

            sensor_columns = df_corrected.columns[1:]
            positive_color = "skyblue"
            negative_color = "black"
            for col in sensor_columns:
                y = df_corrected[col].values
                if apply_threshold and self.threshold is not None:
                    color = positive_color if np.any(y >= self.threshold) else negative_color
                    ax_right.plot(t, y, linestyle='-', color=color)
                else:
                    ax_right.plot(t, y, linestyle='-', color='black')
            if self.threshold is not None:
                ax_right.axhline(y=self.threshold, color='red', linestyle=':', linewidth=1)
            ax_right.set_title("Processed Data")
            ax_right.set_xlabel("Time (min)")
            ax_right.set_ylabel("Processed Sensor Value")
            if apply_threshold and self.threshold is not None:
                pos_handle = Line2D([0], [0], color=positive_color, lw=2, label='Positive')
                neg_handle = Line2D([0], [0], color=negative_color, lw=2, label='Negative')
                thresh_handle = Line2D([0], [0], color='red', lw=1, linestyle=':', label='Threshold')
                ax_right.legend(handles=[pos_handle, neg_handle, thresh_handle], loc="upper left", frameon=False)
        else:
            ax_left.text(0.5, 0.5, "No Data", ha="center", va="center")
            ax_right.text(0.5, 0.5, "No Data", ha="center", va="center")
        self.display_widget.canvas.fig.tight_layout()
        self.display_widget.canvas.draw()

    def apply_threshold(self):
        if self.df_smoothed is None:
            self.log_message("No data loaded to apply threshold.")
            return
        try:
            self.threshold = float(self.thresh_edit.text())
        except ValueError:
            self.log_message("Please enter a valid numeric threshold.")
            return
        time_col = self.df_smoothed.columns[0]
        t = self.df_smoothed[time_col].values
        self.plot_two_subplots(t, self.df_smoothed, self.df_corrected,
                                self.global_ignore_time, self.global_baseline_end_time,
                                apply_threshold=True)
        self.populate_table(self.get_processed_data_with_metadata())
        self.process_deriv_btn.setEnabled(True)
        self.compute_tq_btn.setEnabled(True)
        self.save_tq_btn.setEnabled(False)
        self.log_message("Threshold applied.")

    def process_derivative(self):
        if self.df_to_plot is None or self.threshold is None:
            self.log_message("Please load data and apply threshold before processing derivative.")
            return
        self.inBatchMode = True
        self.nav_widget.show()
        self.current_batch = 0
        self.prepare_batches()
        self.plot_batches()
        self.compute_tq_btn.setEnabled(True)
        self.save_tq_btn.setEnabled(True)
        self.log_message("Derivative processing completed.")

    def prepare_batches(self):
        if self.df_to_plot is None:
            self.total_batches = 0
        else:
            num_sensors = self.df_to_plot.shape[1] - 1
            self.total_batches = ceil(num_sensors / 6)
        time_col = self.df_to_plot.columns[0]
        x = self.df_to_plot[time_col].values
        sensor_min, sensor_max = np.inf, -np.inf
        deriv_min, deriv_max = np.inf, -np.inf
        for col in self.df_to_plot.columns[1:]:
            y = self.df_to_plot[col].values
            sensor_min = min(sensor_min, np.min(y))
            sensor_max = max(sensor_max, np.max(y))
            if np.any(y >= self.threshold):
                dy = np.gradient(y, x)
                deriv_min = min(deriv_min, np.min(dy))
                deriv_max = max(deriv_max, np.max(dy))
                try:
                    p0 = [np.max(dy)-np.min(dy), x[np.argmax(dy)], 1, 1, np.min(dy)]
                    popt, _ = curve_fit(voigt, x, dy, p0=p0)
                    fitted1 = voigt(x, *popt)
                    second_deriv = np.gradient(fitted1, x)
                    deriv_min = min(deriv_min, np.min(second_deriv))
                    deriv_max = max(deriv_max, np.max(second_deriv))
                except:
                    pass
        pad1 = 0.05 * (sensor_max - sensor_min) if (sensor_max - sensor_min) != 0 else 1
        self.common_sensor_ylim = (sensor_min - pad1, sensor_max + pad1)
        pad2 = 0.05 * (deriv_max - deriv_min) if (deriv_max - deriv_min) != 0 else 1
        self.common_deriv_ylim = (deriv_min - pad2, deriv_max + pad2)

    def plot_batches(self):
        self.display_widget.canvas.fig.clf()
        # Create 6 subplots per batch (2 rows x 3 columns).
        axes = self.display_widget.canvas.fig.subplots(2, 3, sharex=True, sharey=False)
        axes = axes.flatten()
        time_col = self.df_to_plot.columns[0]
        x = self.df_to_plot[time_col].values
        start_idx = self.current_batch * 6 + 1
        end_idx = min(start_idx + 6, self.df_to_plot.shape[1])
        
        for i, col in enumerate(self.df_to_plot.columns[start_idx:end_idx]):
            ax_left = axes[i]
            y = self.df_to_plot[col].values
            # Use original color format: Pos sensors are skyblue; Neg are black.
            if np.any(y >= self.threshold):
                raw_color = 'skyblue'
            else:
                raw_color = 'black'
            ax_left.plot(x, y, linestyle='-', color=raw_color)
            ax_left.set_title(col, fontsize=12)
            ax_left.set_ylim(self.common_sensor_ylim)
            
            # Only process fits for positive sensors.
            if np.any(y >= self.threshold):
                # --- Sigmoid Strategy (applied to sensor data) ---
                fitted_sigmoid = None
                try:
                    L_guess = np.max(y) - np.min(y)
                    half_val = np.min(y) + L_guess / 2
                    indices = np.where(y >= half_val)[0]
                    x0_guess = x[indices[0]] if len(indices) > 0 else np.median(x)
                    p0_sigmoid = [L_guess, 1, x0_guess]
                    bounds_sigmoid = ([0, 0, min(x)], [np.inf, np.inf, max(x)])
                    popt_sigmoid, _ = curve_fit(sigmoid_func, x, y, p0=p0_sigmoid, bounds=bounds_sigmoid)
                    fitted_sigmoid = sigmoid_func(x, *popt_sigmoid)
                except Exception as e:
                    self.log_message(f"Sigmoid fit failed for {col}: {e}")
                    fitted_sigmoid = None
                if fitted_sigmoid is not None:
                    ax_left.plot(x, fitted_sigmoid, linestyle=':', color='blue', label="Sigmoid fit")
                    residuals_sig = y - fitted_sigmoid
                    ss_res_sig = np.sum(residuals_sig**2)
                    ss_tot_sig = np.sum((y - np.mean(y))**2)
                    r2_sig = 1 - ss_res_sig/ss_tot_sig if ss_tot_sig != 0 else 0
                    if r2_sig >= 0.75:
                        second_deriv_sig = np.gradient(np.gradient(fitted_sigmoid, x), x)
                        max_idx_sig = np.argmax(second_deriv_sig)
                        tq_sigmoid = x[max_idx_sig]
                        ax_left.axvline(x=tq_sigmoid, color='blue', linestyle='-', linewidth=1, label="Tq_Sigmoid")
                    else:
                        ax_left.text(0.1, 0.8, "Tq_Sigmoid failed", transform=ax_left.transAxes, fontsize=8, color='blue')
                else:
                    ax_left.text(0.1, 0.8, "Tq_Sigmoid failed", transform=ax_left.transAxes, fontsize=8, color='blue')
                
                # --- Voigt Strategy (applied to derivative) ---
                ax_right = ax_left.twinx()
                dy = np.gradient(y, x)
                ax_right.plot(x, dy, linestyle='-', color='magenta', label="df(x)")
                fitted_voigt = None
                try:
                    bounds = ([0, min(x), 0, 0], [np.inf, max(x), np.inf, np.inf])
                    p0 = [np.max(dy)-np.min(dy), x[np.argmax(dy)], 2, 1]
                    popt_voigt, _ = curve_fit(voigt_fit, x, dy, p0=p0, bounds=bounds)
                    fitted_voigt = voigt_fit(x, *popt_voigt)
                except Exception as e:
                    self.log_message(f"Voigt fit failed for {col}: {e}")
                    fitted_voigt = None
                if fitted_voigt is not None:
                    ax_right.plot(x, fitted_voigt, linestyle=':', color='magenta', label="Voigt fit")
                    residuals = dy - fitted_voigt
                    ss_res = np.sum(residuals**2)
                    ss_tot = np.sum((dy - np.mean(dy))**2)
                    r2_voigt = 1 - ss_res/ss_tot if ss_tot != 0 else 0
                    if r2_voigt >= 0.75:
                        second_deriv = np.gradient(fitted_voigt, x)
                        ax_right.plot(x, second_deriv, linestyle='--', color='red', label="d2f(x) Voigt")
                        max_idx = np.argmax(second_deriv)
                        tq_voigt = x[max_idx]
                        ax_right.axvline(x=tq_voigt, color='red', linestyle='-', linewidth=1, label="Tq_Voigt")   
                    else:
                        ax_right.text(0.1, 0.9, "Tq_Voigt failed", transform=ax_right.transAxes, fontsize=8, color='red')
                else:
                    ax_right.text(0.1, 0.9, "Tq_Voigt failed", transform=ax_right.transAxes, fontsize=8, color='red')
                ax_right.set_ylim(self.common_deriv_ylim)
                ax_right.set_ylabel("Derivatives", fontsize=12, fontweight='bold')
            if i == 0:
                ax_left.set_xlabel("Time (min)", fontsize=12, fontweight='bold')
                ax_left.set_ylabel("Sensor Value", fontsize=12, fontweight='bold')
            else:
                ax_left.set_xlabel("")
                ax_left.set_ylabel("")
        for j in range(end_idx - start_idx, 6):
            axes[j].axis('off')
        
        # Legend handles.
        handles = [
            Line2D([0], [0], color='skyblue', lw=2, label='Pos'),
            Line2D([0], [0], color='black', lw=2, label='Neg'),
            Line2D([0], [0], color='magenta', lw=2, label='df(x)'),
            Line2D([0], [0], color='magenta', lw=2, linestyle=':', label='Voigt fit'),
            Line2D([0], [0], color='red', lw=2, linestyle='--', label='d2f(x) Voigt'),
            Line2D([0], [0], color='red', lw=1, linestyle='-', label='Tq_Voigt'),
            Line2D([0], [0], color='blue', lw=2, linestyle=':', label='Sigmoid fit'),
            Line2D([0], [0], color='blue', lw=2, linestyle='-', label='Tq_Sigmoid')
        ]
        if len(axes) > 0:
            axes[0].legend(handles=handles, loc='upper left', fontsize=8, frameon=False)
        self.display_widget.canvas.fig.tight_layout()
        self.display_widget.canvas.draw()
        self.batch_label.setText(f"{self.current_batch+1} / {self.total_batches}")

    def next_batch(self):
        if self.current_batch < self.total_batches - 1:
            self.current_batch += 1
            self.plot_batches()

    def prev_batch(self):
        if self.current_batch > 0:
            self.current_batch -= 1
            self.plot_batches()

    def compute_tq(self):
        """
        Compute Tq for each sensor column using two separate strategies for positive sensors:

        1. Voigt Fit (applied to the derivative of the sensor data):
        - Tq_Voigt is determined as the time at the maximum of the second derivative 
            of the Voigt-fitted derivative.
        - R2_Voigt is computed, and if below 0.75, tq_Voigt is marked as "tq_failed".
        
        2. Sigmoid Fit (applied directly to the sensor data):
        - The sigmoid is fitted to the sensor data, its second derivative is computed,
            and the time at its maximum is taken as tq_Sigmoid.
        - R2_Sigmoid is computed, and if below 0.75, tq_Sigmoid is marked as "Tq_failed".
        
        For sensors not classified as positive, both Tq values are "No Tq".
        """
        if self.df_to_plot is None or self.threshold is None:
            self.log_message("Data or threshold not available for computing Tq.")
            return
        time_col = self.df_to_plot.columns[0]
        x = self.df_to_plot[time_col].values
        results = []
        for col in self.df_to_plot.columns[1:]:
            y = self.df_to_plot[col].values
            label = col
            # Initialize outputs.
            tq_voigt = "No Tq"
            r2_voigt = "N/A"
            tq_sigmoid = "No Tq"
            r2_sigmoid = "N/A"
            
            if np.any(y >= self.threshold):  # Only process positive sensors.
                # --- Voigt Strategy on derivative ---
                dy = np.gradient(y, x)
                fitted_voigt = None
                try:
                    bounds = ([0, min(x), 0, 0], [np.inf, max(x), np.inf, np.inf])
                    p0 = [np.max(dy)-np.min(dy), x[np.argmax(dy)], 2, 1]
                    popt_voigt, _ = curve_fit(voigt_fit, x, dy, p0=p0, bounds=bounds)
                    fitted_voigt = voigt_fit(x, *popt_voigt)
                except Exception as e:
                    self.log_message(f"Voigt fit failed for {col} in compute_tq: {e}")
                    fitted_voigt = None
                if fitted_voigt is not None:
                    residuals = dy - fitted_voigt
                    ss_res = np.sum(residuals**2)
                    ss_tot = np.sum((dy - np.mean(dy))**2)
                    r2 = 1 - ss_res / ss_tot if ss_tot != 0 else 0
                    r2_voigt = r2
                    if r2 >= 0.75:
                        second_deriv = np.gradient(fitted_voigt, x)
                        max_idx = np.argmax(second_deriv)
                        tq_voigt = x[max_idx]
                    else:
                        tq_voigt = "Tq_failed"
                else:
                    tq_voigt = "Tq_failed"
                    r2_voigt = "N/A"
                
                # --- Sigmoid Strategy on sensor data ---
                fitted_sigmoid = None
                try:
                    L_guess = np.max(y) - np.min(y)
                    half_val = np.min(y) + L_guess / 2
                    indices = np.where(y >= half_val)[0]
                    x0_guess = x[indices[0]] if len(indices) > 0 else np.median(x)
                    p0_sigmoid = [L_guess, 1, x0_guess]
                    bounds_sigmoid = ([0, 0, min(x)], [np.inf, np.inf, max(x)])
                    popt_sigmoid, _ = curve_fit(sigmoid_func, x, y, p0=p0_sigmoid, bounds=bounds_sigmoid)
                    fitted_sigmoid = sigmoid_func(x, *popt_sigmoid)
                except Exception as e:
                    self.log_message(f"Sigmoid fit failed for {col} in compute_tq: {e}")
                    fitted_sigmoid = None
                if fitted_sigmoid is not None:
                    residuals_sig = y - fitted_sigmoid
                    ss_res_sig = np.sum(residuals_sig**2)
                    ss_tot_sig = np.sum((y - np.mean(y))**2)
                    r2_sig = 1 - ss_res_sig / ss_tot_sig if ss_tot_sig != 0 else 0
                    r2_sigmoid = r2_sig
                    if r2_sig >= 0.75:
                        # For the sigmoid strategy, compute the second derivative.
                        second_deriv_sig = np.gradient(np.gradient(fitted_sigmoid, x), x)
                        max_idx_sig = np.argmax(second_deriv_sig)
                        tq_sigmoid = x[max_idx_sig]
                    else:
                        tq_sigmoid = "Tq_failed"
                else:
                    tq_sigmoid = "Tq_failed"
                    r2_sigmoid = "N/A"
            results.append((label, tq_voigt, r2_voigt, tq_sigmoid, r2_sigmoid))
            
        sensor_results = pd.DataFrame(results, columns=["Label", "Tq_Voigt", "R2_Voigt", "Tq_Sigmoid", "R2_Sigmoid"])
        metadata = pd.DataFrame({
            "Label": ["Metadata_BaselineStart", "Metadata_BaselineEnd", "Metadata_Threshold", 
                    "Metadata_ExpDuration", "Metadata_Tq-Unit"],
            "Tq_Voigt": [self.global_ignore_time, self.global_baseline_end_time, self.threshold, self.experiment_duration, "minutes"],
            "R2_Voigt": ["N/A"] * 5,
            "Tq_Sigmoid": [self.global_ignore_time, self.global_baseline_end_time, self.threshold, self.experiment_duration, "minutes"],
            "R2_Sigmoid": ["N/A"] * 5
        })
        final_df = pd.concat([metadata, sensor_results], ignore_index=True)
        final_df.columns = list(map(str, final_df.columns))
        self.tq_df = final_df
        self.populate_table(self.tq_df)
        self.log_message("Tq computed.")
        
    def save_tq_data(self):
        if self.tq_df is None:
            self.log_message("No Tq data available. Please compute Tq first.")
            return
        if not self.input_filename:
            self.log_message("Input filename not available.")
            return

        base, ext = os.path.splitext(self.input_filename)
        default_name = base + "_Tq.csv"
        save_path, _ = QFileDialog.getSaveFileName(self, "Save Tq Data", default_name, "CSV Files (*.csv)")
        
        if save_path:
            try:
                with open(save_path, 'w', newline='') as f:
                    writer = csv.writer(f)
                    
                    # Write metadata rows FIRST (these are the first 5 rows of tq_df)
                    for idx in range(5):
                        writer.writerow(self.tq_df.iloc[idx].tolist())
                    
                    # Write column headers (once, after metadata)
                    writer.writerow(self.tq_df.columns.tolist())
                    
                    # Write actual sensor data
                    for idx in range(5, len(self.tq_df)):
                        writer.writerow(self.tq_df.iloc[idx].tolist())

                self.log_message(f"Tq data saved to {save_path}")
            except Exception as e:
                self.log_message(f"Error saving Tq data: {e}")


    def save_processed_data(self):
        if self.df_corrected is None:
            self.log_message("No processed data available.")
            return
        if not self.input_filename:
            self.log_message("Input filename not available.")
            return
        base, ext = os.path.splitext(self.input_filename)
        default_name = base + "_processed.csv"
        save_path, _ = QFileDialog.getSaveFileName(self, "Save Processed Data", default_name, "CSV Files (*.csv)")
        if save_path:
            try:
                with open(save_path, 'w', newline='') as f:
                    writer = csv.writer(f)
                    meta_start = ["Metadata_BaselineStart"] + [self.global_ignore_time] * (len(self.df_corrected.columns) - 1)
                    meta_end = ["Metadata_BaselineStop"] + [self.global_baseline_end_time] * (len(self.df_corrected.columns) - 1)
                    meta_thresh = ["Metadata_Threshold"] + [self.threshold] * (len(self.df_corrected.columns) - 1)
                    meta_duration = ["Metadata_ExpDuration"] + [self.experiment_duration] * (len(self.df_corrected.columns) - 1)
                    writer.writerow(meta_start)
                    writer.writerow(meta_end)
                    writer.writerow(meta_thresh)
                    writer.writerow(meta_duration)
                    writer.writerow(self.df_corrected.columns.tolist())
                    for index, row in self.df_corrected.iterrows():
                        writer.writerow(row.tolist())
                self.log_message(f"Processed data saved to {save_path}")
            except Exception as e:
                self.log_message(f"Error saving processed data: {e}")

    def get_processed_data_with_metadata(self):
        if self.df_corrected is None:
            return None
        df_proc = self.df_corrected.copy()
        df_proc.columns = df_proc.columns.astype(str)
        cols = df_proc.columns.tolist()
        meta_start = {cols[0]: "Metadata_BaselineStart"}
        meta_end = {cols[0]: "Metadata_BaselineEnd"}
        meta_thresh = {cols[0]: "Metadata_Threshold"}
        meta_duration = {cols[0]: "Metadata_ExpDuration"}
        for col in cols[1:]:
            meta_start[col] = self.global_ignore_time
            meta_end[col] = self.global_baseline_end_time
            meta_thresh[col] = self.threshold
            meta_duration[col] = self.experiment_duration
        metadata_df = pd.DataFrame([meta_start, meta_end, meta_thresh, meta_duration])
        new_df = pd.concat([metadata_df, df_proc], ignore_index=True)
        return new_df

    def populate_table(self, df_to_show):
        model = QStandardItemModel(df_to_show.shape[0], df_to_show.shape[1])
        header_labels = list(map(str, df_to_show.columns.tolist()))
        model.setHorizontalHeaderLabels(header_labels)
        cell_font = QFont("Arial", 10)
        for row in range(df_to_show.shape[0]):
            for col in range(df_to_show.shape[1]):
                value = df_to_show.iat[row, col]
                text = f"{value:.2f}" if isinstance(value, float) else str(value)
                item = QStandardItem(text)
                item.setFont(cell_font)
                model.setItem(row, col, item)
        self.populate_table_model(model)

    def populate_table_model(self, model):
        self.display_widget.table_view.setModel(model)
        self.display_widget.table_view.resizeColumnsToContents()
        for row in range(model.rowCount()):
            self.display_widget.table_view.setRowHeight(row, 40)
```
