## Supplementary Software for "ThermiQuant™ MegaScan: High-throughput isothermal reactor with quantitative colorimetric readout for paper-based nucleic acid amplification tests": display_widget.py

```
# display_widget.py
from PyQt5.QtWidgets import QWidget, QVBoxLayout, QTableView, QHeaderView
from matplotlib.backends.backend_qt5agg import FigureCanvasQTAgg as FigureCanvas
from matplotlib.figure import Figure

class MplCanvas(FigureCanvas):
    def __init__(self, parent=None, width=8, height=6, dpi=100):
        fig = Figure(figsize=(width, height), dpi=dpi)
        self.fig = fig
        self.axes = fig.add_subplot(111)
        super().__init__(fig)

class DisplayWidget(QWidget):
    def __init__(self, parent=None):
        super().__init__(parent)
        layout = QVBoxLayout()
        self.canvas = MplCanvas(self, width=8, height=6, dpi=100)
        layout.addWidget(self.canvas)
        self.table_view = QTableView()
        layout.addWidget(self.table_view)
        self.setLayout(layout)
        self.table_view.horizontalHeader().setSectionResizeMode(QHeaderView.Stretch)
        self.table_view.horizontalHeader().setDefaultSectionSize(150)
        self.table_view.verticalHeader().setDefaultSectionSize(40)
```
