## Supplementary Software for "ThermiQuant™ MegaScan: High-throughput isothermal reactor with quantitative colorimetric readout for paper-based nucleic acid amplification tests": image_analysis.py

```
#!/usr/bin/env python
# Module for image processing
# Bibek Raut
# Purdue University

import sys, os, math, json, re, csv
import cv2
import numpy as np
import matplotlib
import time
matplotlib.use("Qt5Agg")
import matplotlib.pyplot as plt
from matplotlib.backends.backend_qt5agg import FigureCanvasQTAgg as FigureCanvas

from datetime import datetime
from ultralytics import YOLO
from PIL import Image
from PIL.ExifTags import TAGS

from PyQt5.QtWidgets import (
    QApplication, QWidget, QVBoxLayout, QHBoxLayout, QPushButton, QLabel,
    QComboBox, QProgressBar, QTextEdit, QFileDialog
)
from PyQt5.QtGui import QFont
from PyQt5.QtCore import Qt

###############################################################################
#                           Helper Functions
###############################################################################
def natural_keys(text):
    return [int(c) if c.isdigit() else c for c in re.split(r'(\d+)', text)]

def get_exif(filename):
    try:
        img = Image.open(filename)
        img.verify()
        return img._getexif()
    except Exception:
        return None

def get_file_dates(filepath):
    created = datetime.fromtimestamp(os.path.getctime(filepath))
    modified = datetime.fromtimestamp(os.path.getmtime(filepath))
    return created, modified

def get_original_date(filename, logger=None):
    exif = get_exif(filename)
    if not exif:
        if logger: logger(f"No EXIF for {os.path.basename(filename)}; using file creation time.")
        return get_file_dates(filename)[0].timestamp()
    labeled = {TAGS.get(k, k): v for k, v in exif.items()}
    date_str = labeled.get("DateTimeDigitized") or labeled.get("DateTimeOriginal")
    if not date_str:
        if logger: logger(f"No EXIF DateTime* for {os.path.basename(filename)}; using file creation time.")
        return get_file_dates(filename)[0].timestamp()
    try:
        return datetime.strptime(date_str, "%Y:%m:%d %H:%M:%S").timestamp()
    except Exception:
        if logger: logger(f"Bad EXIF date in {os.path.basename(filename)}; using file creation time.")
        return get_file_dates(filename)[0].timestamp()

def populate_table(self, df):
    from PyQt5.QtGui import QStandardItemModel, QStandardItem
    model = QStandardItemModel(df.shape[0], df.shape[1])
    model.setHorizontalHeaderLabels(df.columns.tolist())
    for row in range(df.shape[0]):
        for col in range(df.shape[1]):
            value = df.iat[row, col]
            item = QStandardItem(str(value))
            model.setItem(row, col, item)
    self.display_widget.table_view.setModel(model)

def load_and_rotate_image(image_path, rotation_angle=0):
    img = cv2.imread(image_path)
    if img is None:
        print(f"Unable to load image: {image_path}")
        return None
    if rotation_angle == 0:
        return img
    (h, w) = img.shape[:2]
    center = (w//2, h//2)
    M = cv2.getRotationMatrix2D(center, rotation_angle, 1.0)
    cos_val = abs(M[0,0]); sin_val = abs(M[0,1])
    new_w = int((h * sin_val) + (w * cos_val))
    new_h = int((h * cos_val) + (w * sin_val))
    M[0,2] += (new_w//2) - center[0]
    M[1,2] += (new_h//2) - center[1]
    rotated = cv2.warpAffine(img, M, (new_w, new_h))
    return rotated

def run_yolo_detector(image, model_path, conf, iou):
    model = YOLO(model_path)
    results = model.predict(image, conf=conf, iou=iou)
    return results

def create_directories(base_directory, dir_name="combined_patches"):
    path = os.path.join(base_directory, dir_name)
    os.makedirs(path, exist_ok=True)
    return path

###############################################################################
#           Linear Grid Chip – Specific Functions
###############################################################################
LEFT_SHRINK = 0.05
RIGHT_SHRINK = 0.05
TOP_SHRINK = 0.05
BOTTOM_SHRINK = 0.05
HUE_MIN_CUTOFF = 50
HUE_MAX_CUTOFF = 130
FIXED_PATCH_SIZE = (64, 64)

def extract_bboxes_linear_grid(results):
    if not results:
        return []
    boxes_out = []
    for box in results[0].boxes:
        x1, y1, x2, y2 = box.xyxy[0].cpu().numpy()
        boxes_out.append((int(x1), int(y1), int(x2), int(y2)))
    return boxes_out

def filter_segmented_patches(bounding_boxes):
    if not bounding_boxes:
        return []
    patches = []
    for i, (x1, y1, x2, y2) in enumerate(bounding_boxes):
        w = x2 - x1
        h = y2 - y1
        x1 += int(0.1 * w)
        x2 -= int(0.1 * w)
        y1 += int(0.1 * h)
        y2 -= int(0.1 * h)
        w2 = x2 - x1
        h2 = y2 - y1
        cx = (x1 + x2) // 2
        cy = (y1 + y2) // 2
        area = w2 * h2
        patches.append({
            "index": i+1,
            "bounding_box": (x1, y1, x2, y2),
            "center": (cx, cy),
            "area": area
        })
    largest_patch = max(patches, key=lambda p: p["area"])
    threshold_area = 0.25 * largest_patch["area"]
    patches = [p for p in patches if p["area"] >= threshold_area]
    patches_sorted = sorted(patches, key=lambda p: p["area"])
    if not patches_sorted:
        return []
    return patches_sorted

def sort_and_label_patches(patches):
    if not patches:
        return []
    sorted_patches = []
    alpha_labels = list("ABCDEFGHIJKLMNOPQRSTUVWXYZ")
    row_label_counter = 0
    to_process = sorted(patches, key=lambda p: p["center"][1])
    while to_process and row_label_counter < len(alpha_labels):
        row_letter = alpha_labels[row_label_counter]
        top_patch = to_process[0]
        patch_h = top_patch["bounding_box"][3] - top_patch["bounding_box"][1]
        row_y_min = top_patch["center"][1] - patch_h
        row_y_max = top_patch["center"][1] + patch_h
        same_row = [p for p in to_process if row_y_min <= p["center"][1] <= row_y_max]
        same_row.sort(key=lambda p: p["center"][0])
        for i, p in enumerate(same_row, start=1):
            p["label"] = f"{row_letter}{i}"
        sorted_patches.extend(same_row)
        used_ids = set(id(p) for p in same_row)
        to_process = [p for p in to_process if id(p) not in used_ids]
        row_label_counter += 1
    return sorted_patches

###############################################################################
#           Drawing / Patch Extraction / Hue (Linear only)
###############################################################################
def draw_metadata_annotations(full_image, metadata, is_circular=False):
    ann = full_image.copy()
    h_img, w_img = ann.shape[:2]
    font = cv2.FONT_HERSHEY_SIMPLEX

    text_color = (255, 255, 255)
    bg_color = (0, 0, 0)

    # Linear Grid chips
    for pad in metadata.get("pads", []):
        bbox = pad["bbox"]
        label = pad["label"]
        (x1, y1, x2, y2) = map(int, bbox)
        cv2.rectangle(ann, (x1, y1), (x2, y2), (0, 255, 0), 3)
        pad_height = y2 - y1
        desired_text_height = pad_height * 0.6
        (base_w, base_h), _ = cv2.getTextSize("A", font, 1.0, 1)
        font_scale = desired_text_height / base_h if base_h else 1.0
        thickness = int(max(1, font_scale * 2))
        (tw, th), base = cv2.getTextSize(label, font, font_scale, thickness)
        cv2.rectangle(ann, (x1, y1 - th - base), (x1 + tw, y1), bg_color, -1)
        cv2.putText(ann, label, (x1, y1 - base), font, font_scale, text_color, thickness)
    return ann

def crop_and_store_patches_combined_from_metadata_linear_grid(image, metadata, output_directory, image_index, fixed_size=FIXED_PATCH_SIZE, save_labeled_first_image=False):
    patch_list = []
    for p in metadata.get("pads", []):
        patch_list.append({
            "label": p["label"],
            "bounding_box": tuple(p["bbox"])
        })
    if not patch_list:
        placeholder = np.zeros((fixed_size[1], fixed_size[0], 3), dtype=np.uint8)
        fname = f"ProcessedImage_{image_index+1}.png"
        cv2.imwrite(os.path.join(output_directory, fname), placeholder)
        return placeholder, []
    labels = [p["label"] for p in patch_list]
    row_letters = sorted(list({lbl[0] for lbl in labels}))
    col_nums = sorted(list({int(lbl[1:]) for lbl in labels if lbl[1:].isdigit()}))
    row_map = {letter: i for i, letter in enumerate(row_letters)}
    col_map = {num: i for i, num in enumerate(col_nums)}
    num_rows = len(row_letters)
    num_cols = len(col_nums)
    ph, pw = fixed_size
    combined_h = ph * num_rows
    combined_w = pw * num_cols
    combined = np.zeros((combined_h, combined_w, 3), dtype=np.uint8)
    positions = []
    for p in patch_list:
        x1, y1, x2, y2 = p["bounding_box"]
        sub = image[y1:y2, x1:x2]
        patch_resized = cv2.resize(sub, fixed_size)
        row_letter = p["label"][0]
        try:
            col_num = int(p["label"][1:])
        except:
            col_num = 1
        row_idx = row_map.get(row_letter, 0)
        col_idx = col_map.get(col_num, 0)
        sy = row_idx * ph
        sx = col_idx * pw
        combined[sy:sy+ph, sx:sx+pw] = patch_resized
        positions.append({"label": p["label"], "start_x": sx, "start_y": sy})
    outname = f"ProcessedImage_{image_index+1}.png"
    outpath = os.path.join(output_directory, outname)
    cv2.imwrite(outpath, combined)
    if save_labeled_first_image and (image_index == 0):
        labeled = combined.copy()
        for pos in positions:
            cv2.putText(labeled, pos["label"], (pos["start_x"]+5, pos["start_y"]+20),
                        cv2.FONT_HERSHEY_SIMPLEX, 0.6, (0,255,0), 2)
        proc_data = os.path.join(os.path.dirname(output_directory), "processed_data")
        os.makedirs(proc_data, exist_ok=True)
        cv2.imwrite(os.path.join(proc_data, "first_combined_image_with_labels.png"), labeled)
    return combined, positions

def calculate_hue_for_patches_in_image_combined(combined_image, positions, hue_min=HUE_MIN_CUTOFF, hue_max=HUE_MAX_CUTOFF, threshold=0.9):
    hue_results = {}
    ph, pw = 64, 64
    for patch in positions:
        label = patch['label']
        try:
            x, y = patch['start_x'], patch['start_y']
            subimg = combined_image[y:y+ph, x:x+pw]
            hsv = cv2.cvtColor(subimg, cv2.COLOR_BGR2HSV)
            hues = hsv[:,:,0].astype(np.int16).ravel()
            total = len(hues)
            filtered = [h for h in hues if h < hue_min or h > hue_max]
            if total > 0 and (len(filtered)/total) < threshold:
                avg = "N/A"
            elif filtered:
                mod_hues = [h - 179 if h > hue_max else h for h in filtered]
                avg = np.round(np.mean(mod_hues), 1)
            else:
                avg = "N/A"
            hue_results[label] = avg
        except Exception as e:
            print(f"Error processing hue for {label}: {e}")
            hue_results[label] = "N/A"
    return hue_results

def get_expected_labels_linear_grid(metadata):
    labels = []
    for p in metadata.get("pads", []):
        lab = p["label"]
        if lab not in labels:
            labels.append(lab)
    return labels

def save_hue_results_to_csv(hue_results, time_min, csv_path, expected_labels, overwrite=False, retries=5, delay=0.5):
    """
    Saves hue results to a CSV file with retry logic to handle file locks.
    """
    header = ["Time_min"] + expected_labels
    row = [time_min] + [hue_results.get(lbl, None) for lbl in expected_labels]
    mode = "w" if overwrite else "a"

    for attempt in range(1, retries + 1):
        try:
            with open(csv_path, mode, newline="") as f:
                writer = csv.writer(f)
                if overwrite:
                    writer.writerow(header)
                writer.writerow(row)
            return
        except PermissionError as e:
            if attempt < retries:
                print(f"⚠️ CSV file locked (attempt {attempt}/{retries}), retrying in {delay} sec...")
                time.sleep(delay)
            else:
                raise e

###############################################################################
#                          Matplotlib Canvas
###############################################################################
class MplCanvas(FigureCanvas):
    def __init__(self, parent=None):
        fig, ax = plt.subplots()
        super().__init__(fig)
        self.fig = fig
        self.ax = ax
        self.fig.tight_layout()

###############################################################################
#                       Image Analysis Widget
###############################################################################
class ImageAnalysisWidget(QWidget):
    def showEvent(self, event):
        # (Line ~350) Clear the shared canvas (figure) and table view when the widget is shown.
        self.display_widget.canvas.fig.clf()
        self.display_widget.canvas.axes = self.display_widget.canvas.fig.add_subplot(111)
        self.display_widget.canvas.draw()
        if hasattr(self.display_widget, "table_view"):
            self.display_widget.table_view.setModel(None)
        super().showEvent(event)

    def __init__(self, display_widget, parent=None):
        super().__init__(parent)
        self.setFont(QFont("Arial", 11))
        style_sheet = """
        QPushButton {
            background-color: #555555;
            color: white;
            border: 1px solid #444444;
            border-radius: 4px;
            padding: 5px 10px;
            font: 11pt "Arial";
        }
        QPushButton:disabled {
            background-color: #cccccc;
            color: #666666;
            border: 1px solid #aaaaaa;
        }
        QLineEdit, QComboBox {
            background-color: #ffffff;
            color: #333333;
            border: 1px solid #888888;
            border-radius: 4px;
            padding: 3px 5px;
            font: 10pt "Arial";
        }
        QLabel {
            font: 10pt "Arial";
            color: #333333;
        }
        """
        self.setStyleSheet(style_sheet)

        self.current_folder = None
        self.current_image = None
        self.metadata = None
        self.annotated_image = None
        self.display_widget = display_widget

        main_layout = QVBoxLayout()

        # Folder Selection
        folder_layout = QHBoxLayout()
        self.btn_select_folder = QPushButton("Select Folder")
        self.btn_select_folder.setCheckable(True)
        self.btn_select_folder.clicked.connect(self.select_folder)
        folder_layout.addWidget(self.btn_select_folder)
        main_layout.addLayout(folder_layout)

        # Chip Type Selection (single option retained)
        chip_layout = QHBoxLayout()
        chip_layout.addWidget(QLabel("Chip Type:"))
        self.combo_chip_type = QComboBox()
        self.combo_chip_type.addItem("Linear Grid")
        self.combo_chip_type.setStyleSheet("""
            QComboBox {
                background-color: #d3d3d3;
                padding: 5px;
                border: 1px solid #808080;
                font-size: 14px;
            }
            QComboBox::drop-down {
                border: none;
            }
        """)
        chip_layout.addWidget(self.combo_chip_type)
        main_layout.addLayout(chip_layout)

        # Metadata Generation Button
        meta_layout = QHBoxLayout()
        self.btn_generate_metadata = QPushButton("Generate Metadata")
        self.btn_generate_metadata.setCheckable(True)
        self.btn_generate_metadata.clicked.connect(self.generate_metadata)
        meta_layout.addWidget(self.btn_generate_metadata)
        main_layout.addLayout(meta_layout)

        # Process Images Button
        proc_layout = QHBoxLayout()
        self.btn_process_images = QPushButton("Process Images")
        self.btn_process_images.setCheckable(True)
        self.btn_process_images.clicked.connect(self.process_images)
        proc_layout.addWidget(self.btn_process_images)
        main_layout.addLayout(proc_layout)

        # Progress Bar and Log
        self.progress_bar = QProgressBar()
        self.progress_bar.setValue(0)
        main_layout.addWidget(self.progress_bar)

        self.log = QTextEdit()
        self.log.setReadOnly(True)
        main_layout.addWidget(self.log)

        self.setLayout(main_layout)

    def log_message(self, msg):
        self.log.append(msg)
        QApplication.processEvents()

    def select_folder(self):
        folder = QFileDialog.getExistingDirectory(self, "Select Folder", "")
        if folder:
            self.current_folder = folder
            self.log_message(f"Folder selected: {folder}")
            valid_files = sorted(
                [f for f in os.listdir(folder) if f.lower().endswith(('.jpg','.jpeg','.png','.tif'))],
                key=natural_keys
            )
            if valid_files:
                first_img_path = os.path.join(folder, valid_files[0])
                img = load_and_rotate_image(first_img_path, 0)
                if img is not None:
                    self.current_image = img
                    self.display_widget.canvas.axes.clear()
                    self.display_widget.canvas.axes.imshow(cv2.cvtColor(img, cv2.COLOR_BGR2RGB))
                    self.display_widget.canvas.axes.set_xticks([])
                    self.display_widget.canvas.axes.set_yticks([])
                    self.display_widget.canvas.axes.set_title("First Image")
                    self.display_widget.canvas.draw()
                    self.log_message("First image displayed.")
            else:
                self.log_message("No image files found in the folder.")

    def generate_metadata(self):
        if self.current_image is None:
            self.log_message("Please select a folder with images first.")
            return

        # Create metadata folder
        metadata_folder = os.path.join(self.current_folder, "metadata")
        os.makedirs(metadata_folder, exist_ok=True)

        # Linear Grid Chip Analysis Branch
        lib_dir = os.path.dirname(os.path.abspath(__file__))
        base_dir = os.path.dirname(lib_dir)
        pad_model_path = os.path.join(base_dir, "model", "phenol_red_paper_pad_yolo.pt")
        pad_results = run_yolo_detector(self.current_image, pad_model_path, conf=0.5, iou=0.1)
        bboxes = extract_bboxes_linear_grid(pad_results)
        filtered = sort_and_label_patches(filter_segmented_patches(bboxes))
        metadata_dict = {
            "image_info": {
                "filename": "LoadedImage",
                "width": self.current_image.shape[1],
                "height": self.current_image.shape[0]
            },
            "pads": []
        }
        for p in filtered:
            (x1, y1, x2, y2) = p["bounding_box"]
            metadata_dict["pads"].append({
                "label": p["label"],
                "bbox": [x1, y1, x2, y2]
            })
        self.metadata = metadata_dict
        self.log_message("Linear Grid chip metadata generated.")
        annotated = draw_metadata_annotations(self.current_image, self.metadata, is_circular=False)

        # Save metadata + labeled image
        metadata_path = os.path.join(metadata_folder, "image_metadata.json")
        with open(metadata_path, "w") as jf:
            json.dump(self.metadata, jf, indent=4)
        self.log_message(f"Metadata saved to: {metadata_path}")

        labelled_image_path = os.path.join(metadata_folder, "AnnotatedImage_with_labels.png")
        cv2.imwrite(labelled_image_path, annotated)
        self.log_message(f"Labelled annotated image saved to: {labelled_image_path}")

        self.annotated_image = annotated
        self.display_widget.canvas.axes.clear()
        self.display_widget.canvas.axes.imshow(cv2.cvtColor(self.annotated_image, cv2.COLOR_BGR2RGB))
        self.display_widget.canvas.axes.set_xticks([])
        self.display_widget.canvas.axes.set_yticks([])
        self.display_widget.canvas.draw()
        self.log_message("Annotated image displayed.")

    def process_images(self):
        if not self.current_folder:
            self.log_message("No folder selected.")
            return
        if self.metadata is None:
            self.log_message("Please generate metadata first.")
            return

        # Clear the canvas before processing.
        self.display_widget.canvas.axes.clear()
        self.display_widget.canvas.draw()
        self.log_message("Processing images...")

        valid_files = sorted(
            [f for f in os.listdir(self.current_folder) if f.lower().endswith(('.jpg','.jpeg','.png','.tif'))],
            key=natural_keys
        )
        if len(valid_files) < 1:
            self.log_message("No images to process.")
            return

        first_path = os.path.join(self.current_folder, valid_files[0])
        init_time = get_original_date(first_path, logger=self.log_message)
        cum_time = 0.0

        patches_dir = create_directories(self.current_folder, "combined_patches")
        proc_data = os.path.join(self.current_folder, "processed_data")
        os.makedirs(proc_data, exist_ok=True)
        csv_path = os.path.join(proc_data, "csv_hue_mean.csv")
        self.progress_bar.setMaximum(len(valid_files))

        for i, fn in enumerate(valid_files, start=0):
            path = os.path.join(self.current_folder, fn)
            img = load_and_rotate_image(path, 0)
            if img is None:
                self.log_message(f"Skipping {fn}; could not load.")
                continue

            cur_time = get_original_date(path, logger=self.log_message)
            elapsed = 0.0 if i == 0 else (cur_time - init_time) / 60.0
            cum_time += elapsed
            init_time = cur_time

            combined, positions = crop_and_store_patches_combined_from_metadata_linear_grid(
                img, self.metadata, patches_dir, i, fixed_size=FIXED_PATCH_SIZE, save_labeled_first_image=(i==0)
            )

            hue_results = calculate_hue_for_patches_in_image_combined(
                combined, positions, hue_min=HUE_MIN_CUTOFF, hue_max=HUE_MAX_CUTOFF, threshold=0.9
            )

            expected_labels = get_expected_labels_linear_grid(self.metadata)
            overwrite = (i == 0)
            save_hue_results_to_csv(hue_results, round(cum_time,2), csv_path, expected_labels, overwrite=overwrite)
            self.log_message(f"Processed {fn} => t={round(cum_time,2)} min")
            self.progress_bar.setValue(i + 1)

        self.plot_hue_csv(csv_path, "Hue Data Over Time (Linear Grid)")

    def plot_hue_csv(self, csv_path, title_str):
        try:
            import pandas as pd
            df = pd.read_csv(csv_path)
        except Exception as e:
            self.log_message(f"Error reading CSV: {e}")
            return

        self.display_widget.canvas.fig.clf()
        self.display_widget.canvas.axes = self.display_widget.canvas.fig.add_subplot(111)

        for col in df.columns[1:]:
            df[col] = pd.to_numeric(df[col], errors='coerce')
            self.display_widget.canvas.axes.plot(df["Time_min"], df[col], label=col)
        self.display_widget.canvas.axes.set_xlabel("Time (min)")
        self.display_widget.canvas.axes.set_ylabel("Hue")
        self.display_widget.canvas.fig.tight_layout()
        self.display_widget.canvas.draw()
        self.log_message("Hue plot displayed.")

        populate_table(self, df)

if __name__ == "__main__":
    from PyQt5.QtWidgets import QApplication
    from display_widget import DisplayWidget
    app = QApplication(sys.argv)
    display = DisplayWidget()
    widget = ImageAnalysisWidget(display)
    widget.show()
    sys.exit(app.exec_())
```
