## Supplementary figures and images for "ThermiQuant™ MegaScan: High-throughput isothermal reactor with quantitative colorimetric readout for paper-based nucleic acid amplification tests"

### Timelapse_Image11.jpg

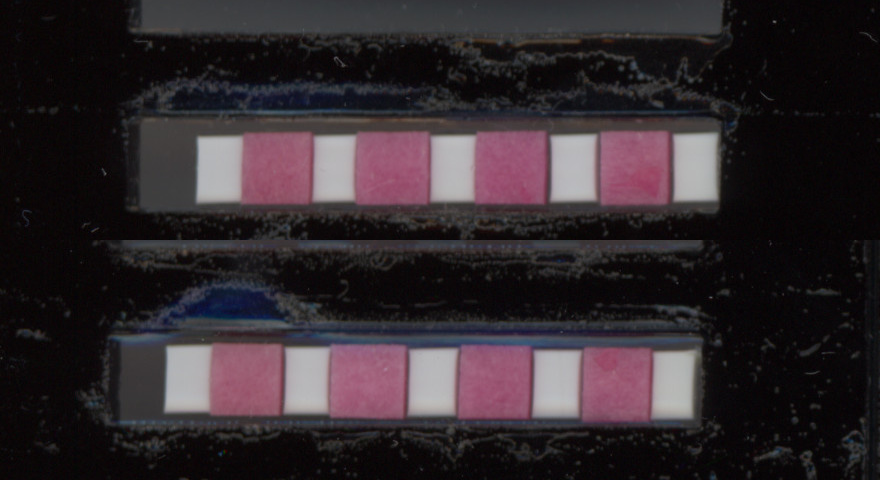

### Timelapse_Image12.jpg

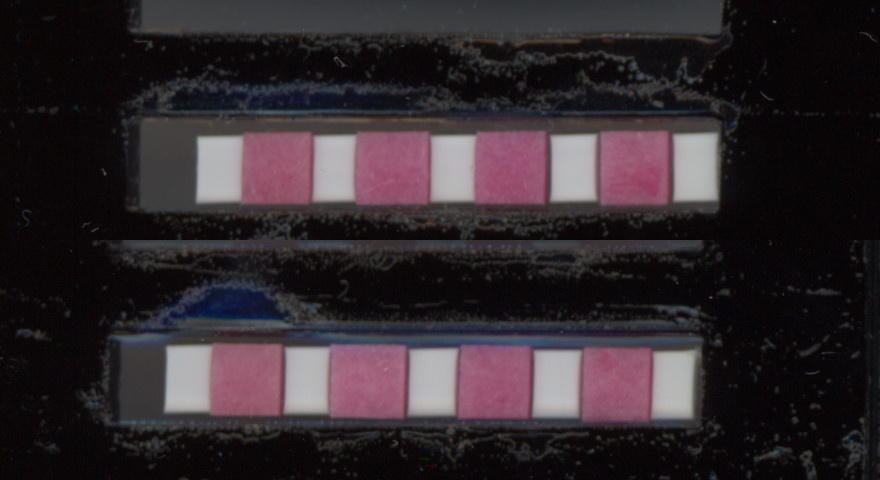

### Timelapse_Image13.jpg

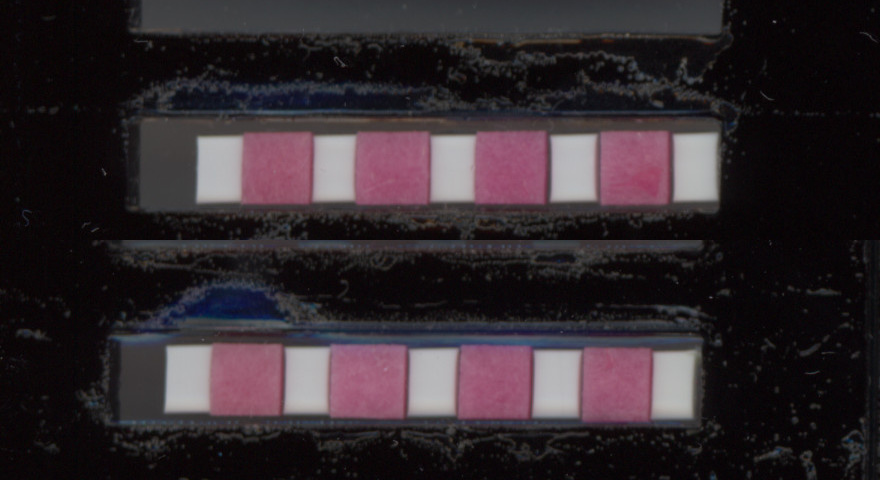

### Timelapse_Image14.jpg

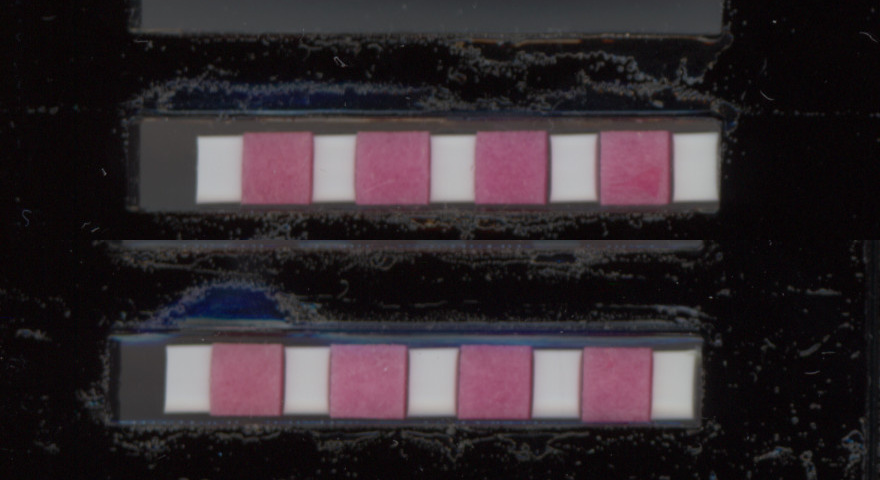

### Timelapse_Image15.jpg

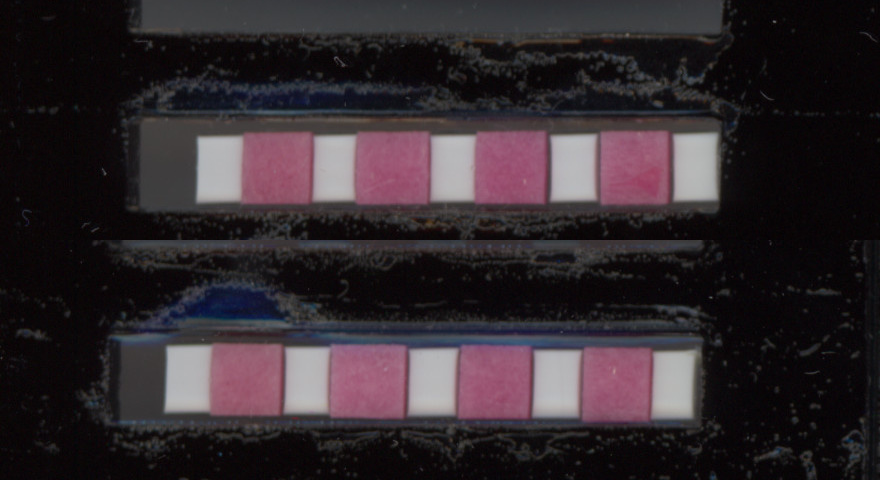

### Timelapse_Image16.jpg

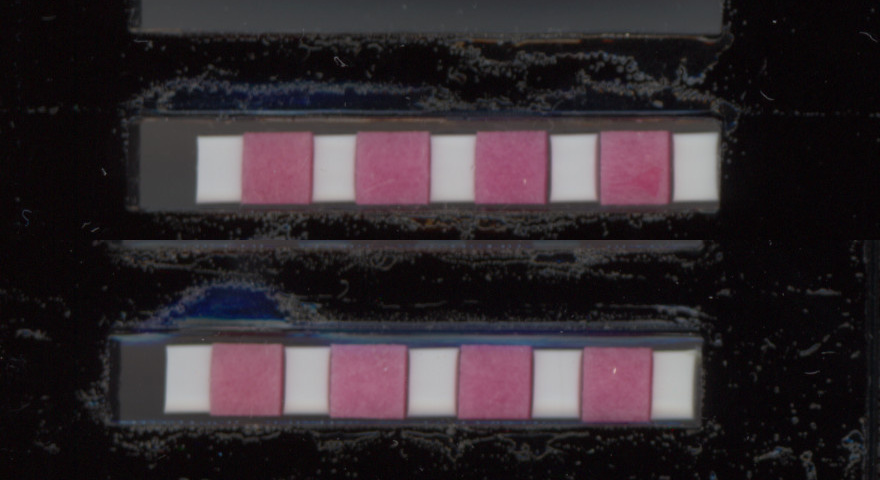

### Timelapse_Image17.jpg

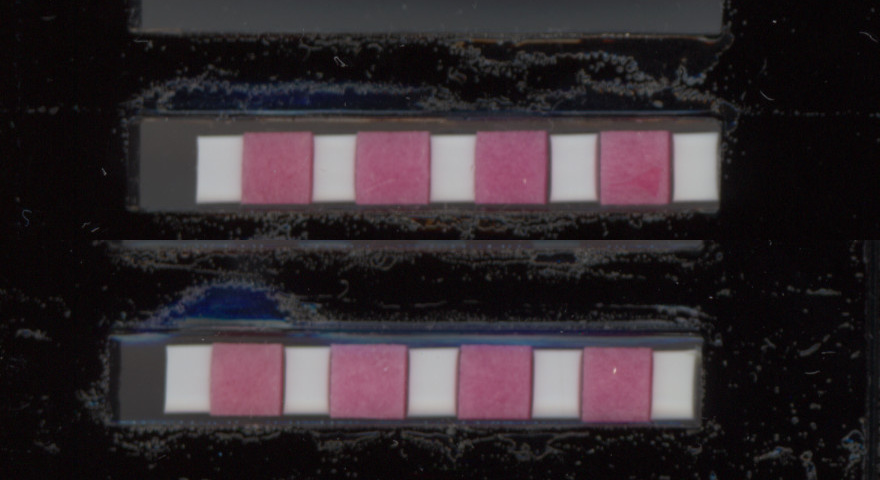

### Timelapse_Image18.jpg

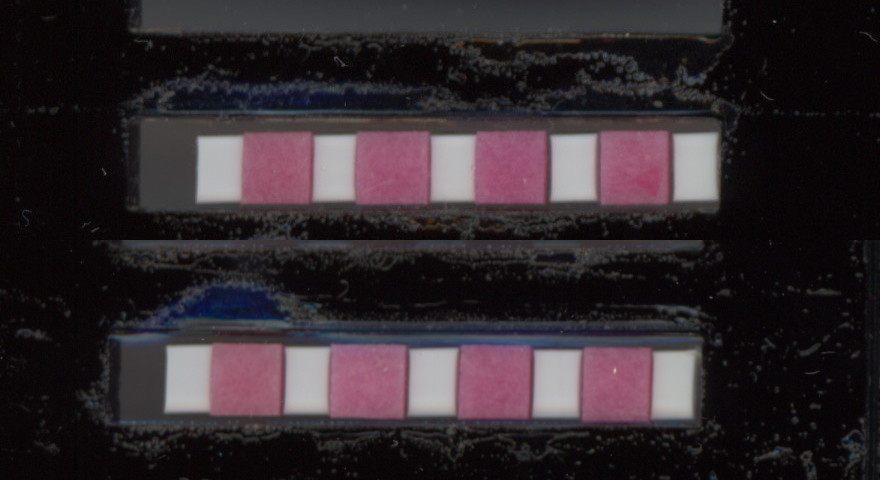

### Timelapse_Image19.jpg

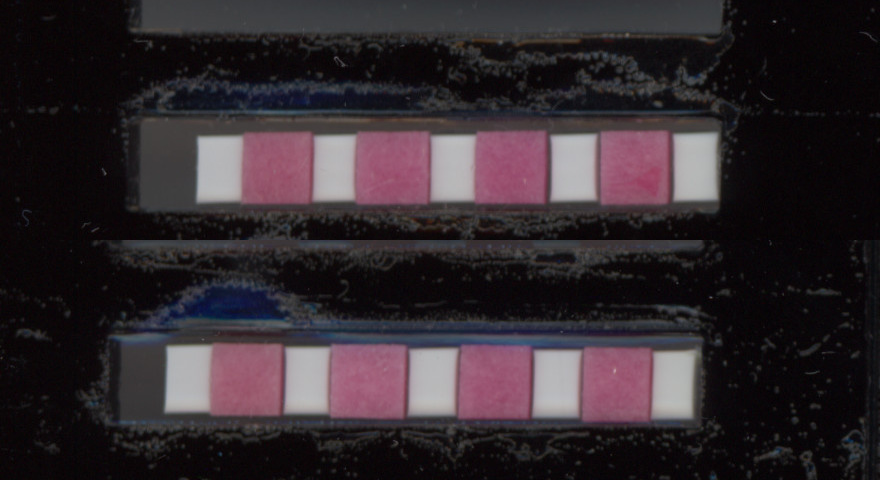

### Timelapse_Image20.jpg

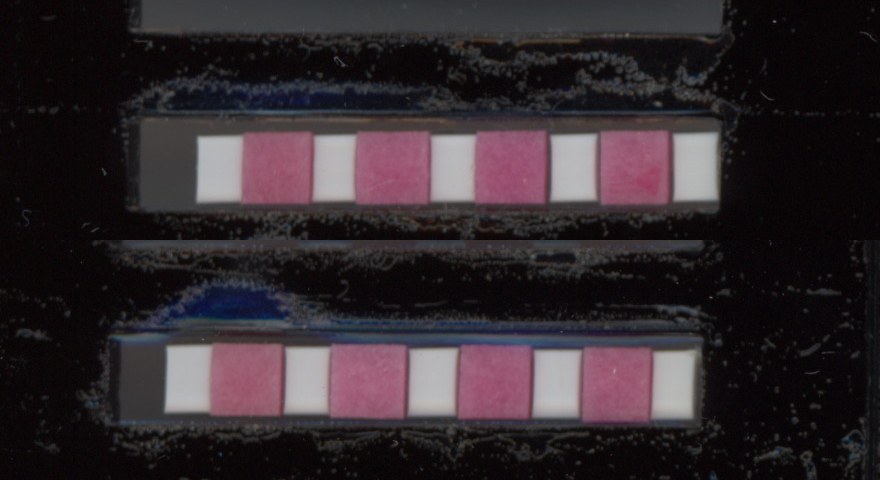

### Timelapse_Image21.jpg

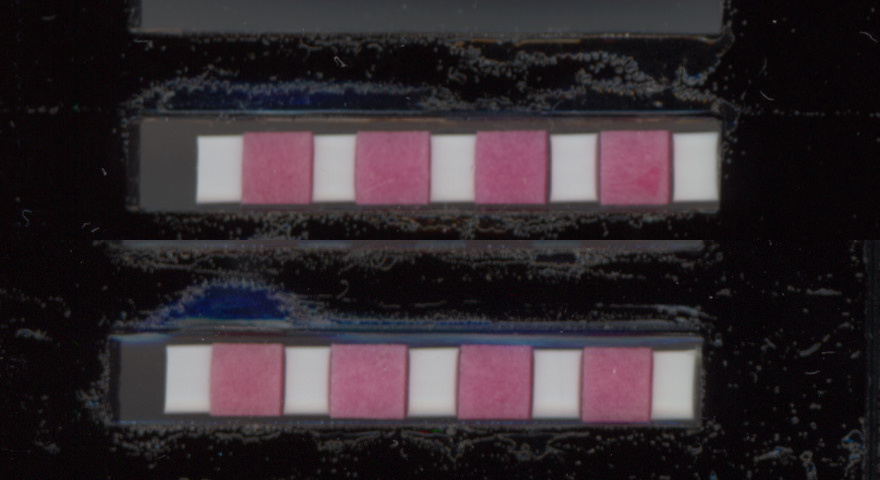

### Timelapse_Image22.jpg

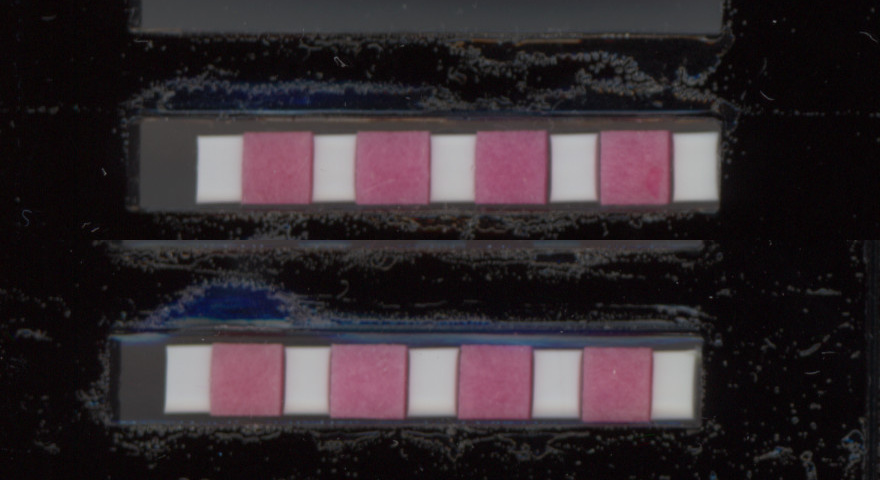

### Timelapse_Image23.jpg

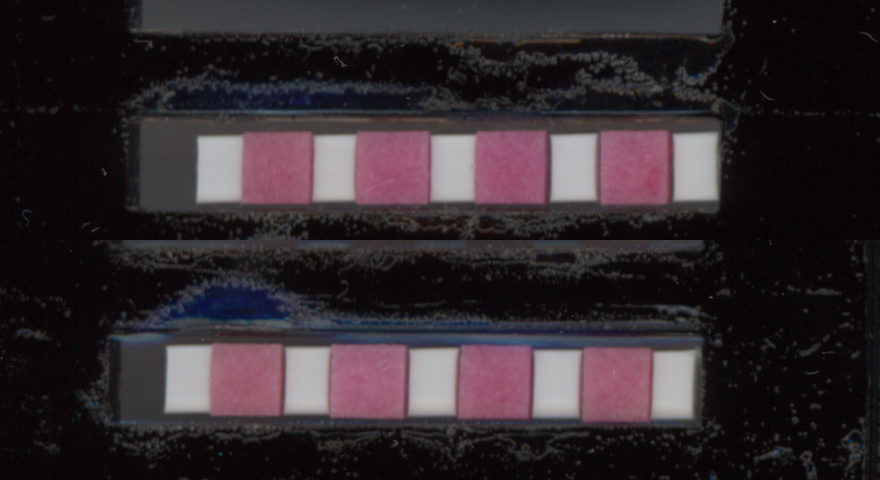

### Timelapse_Image24.jpg

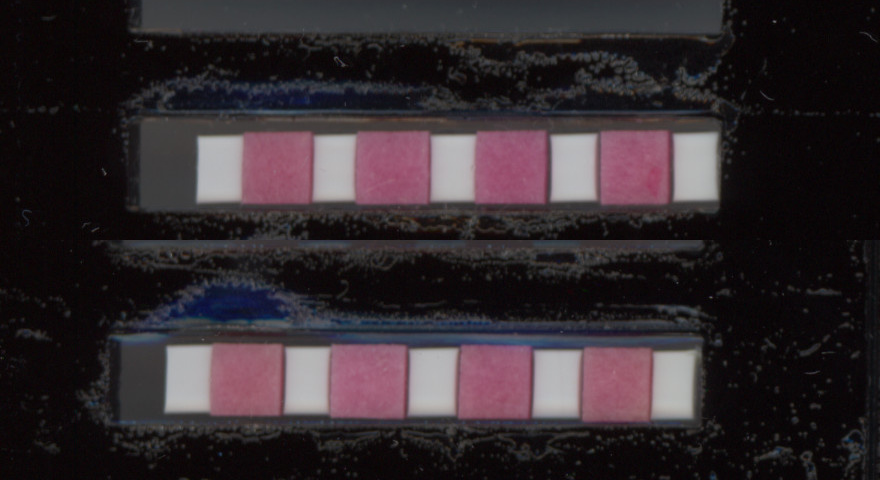

### Timelapse_Image25.jpg

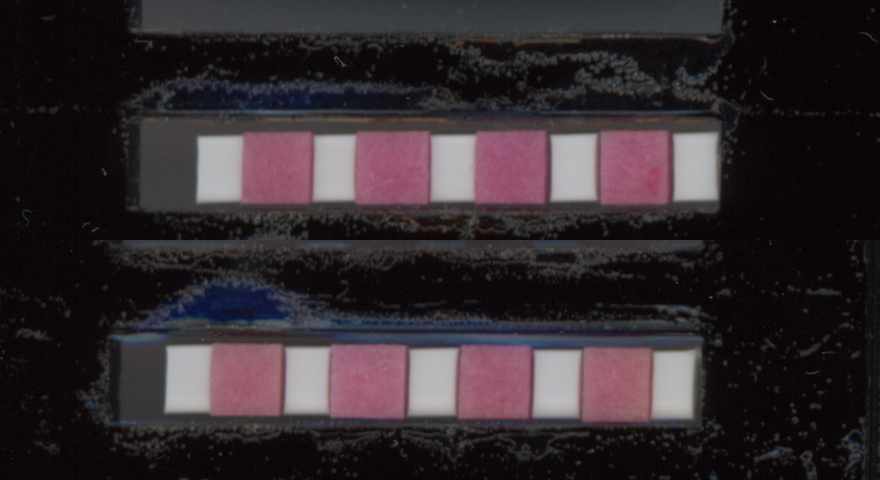

### Timelapse_Image26.jpg

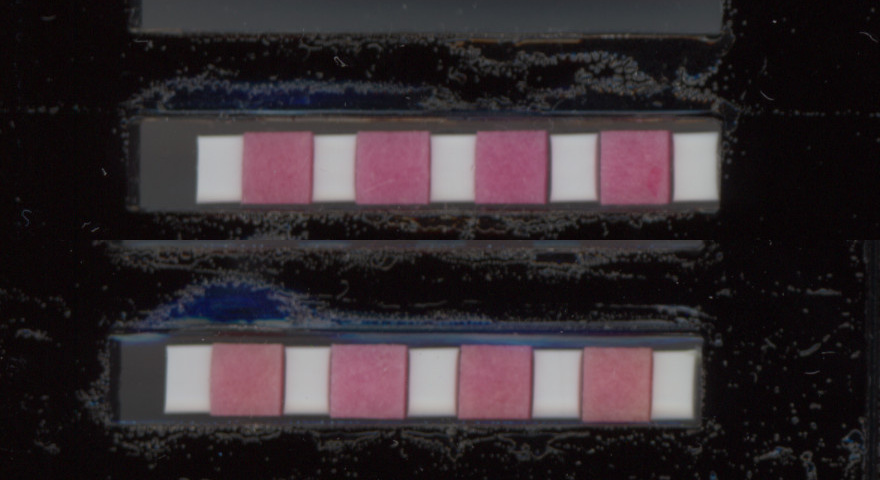

### Timelapse_Image27.jpg

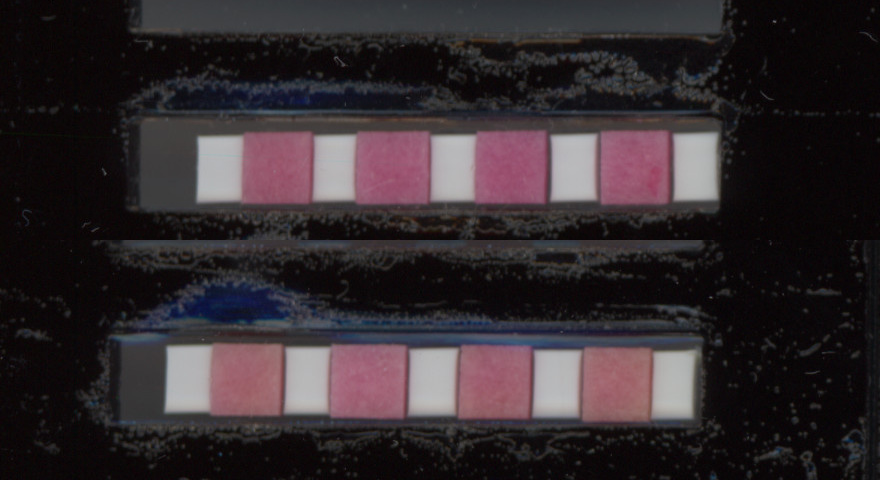

### Timelapse_Image28.jpg

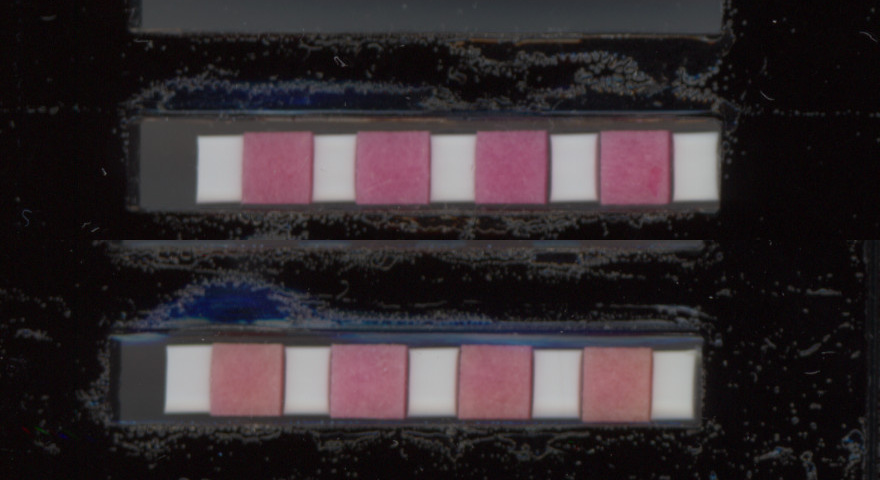

### Timelapse_Image29.jpg

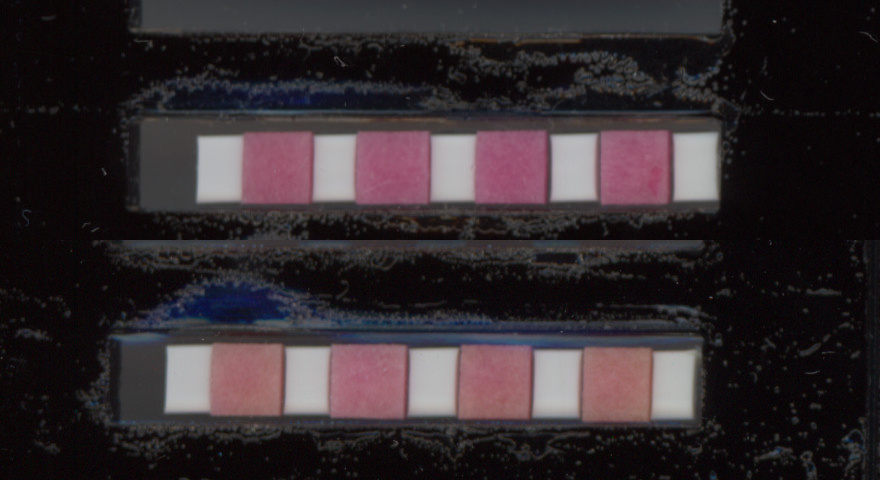

### Timelapse_Image30.jpg

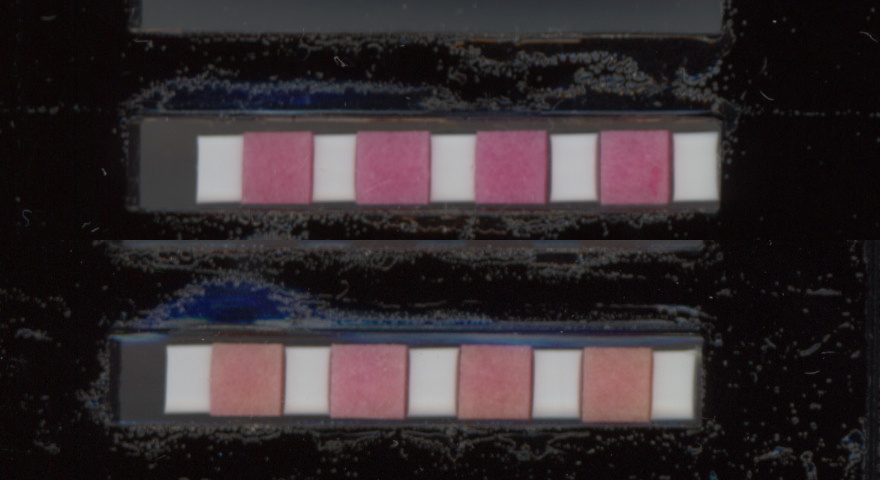

### Timelapse_Image31.jpg

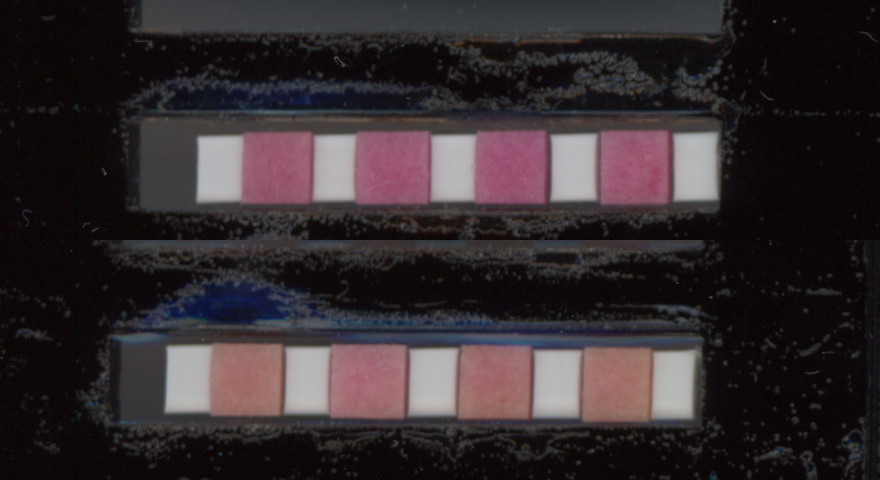

### Timelapse_Image32.jpg

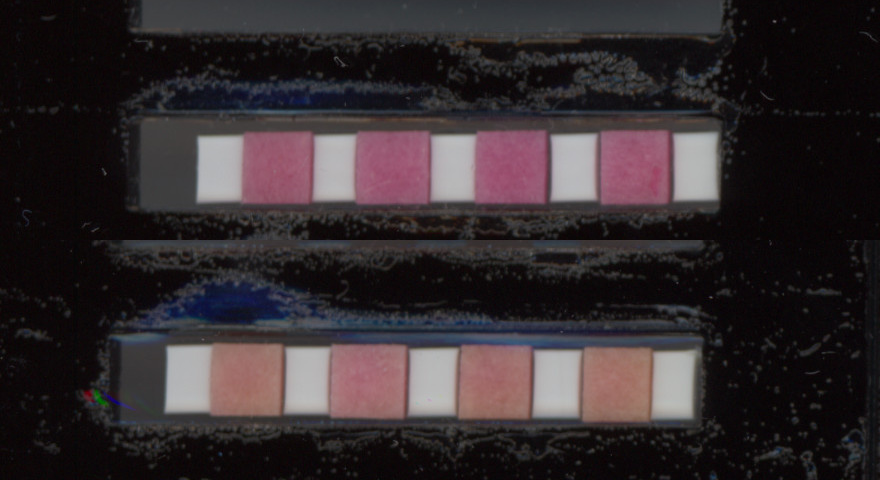

### Timelapse_Image33.jpg

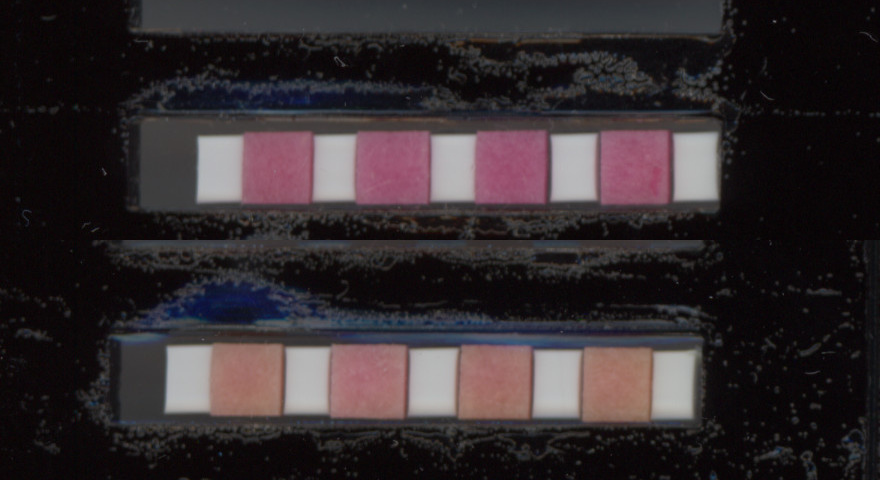

### Timelapse_Image34.jpg

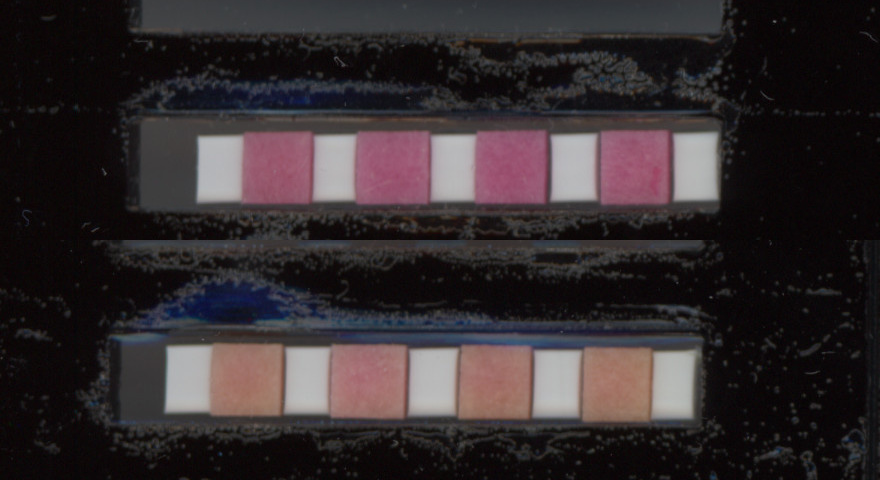

### Timelapse_Image35.jpg

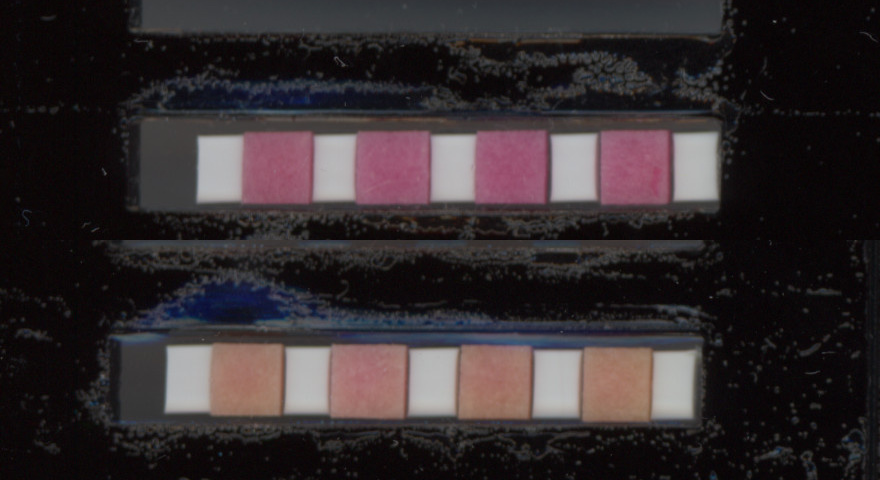

### Timelapse_Image36.jpg

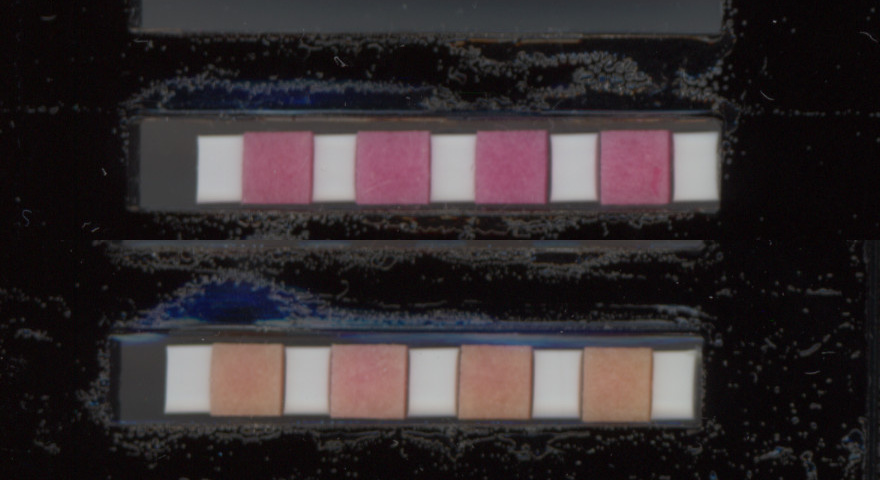

### Timelapse_Image37.jpg

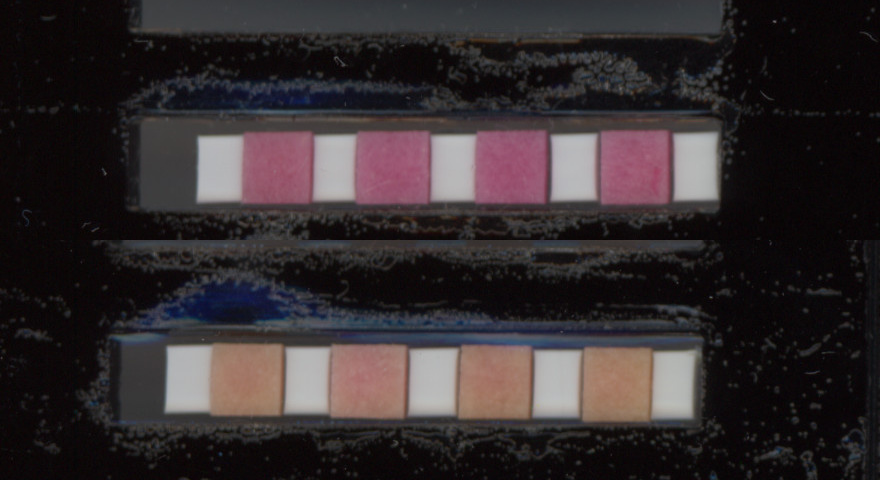

### Timelapse_Image38.jpg

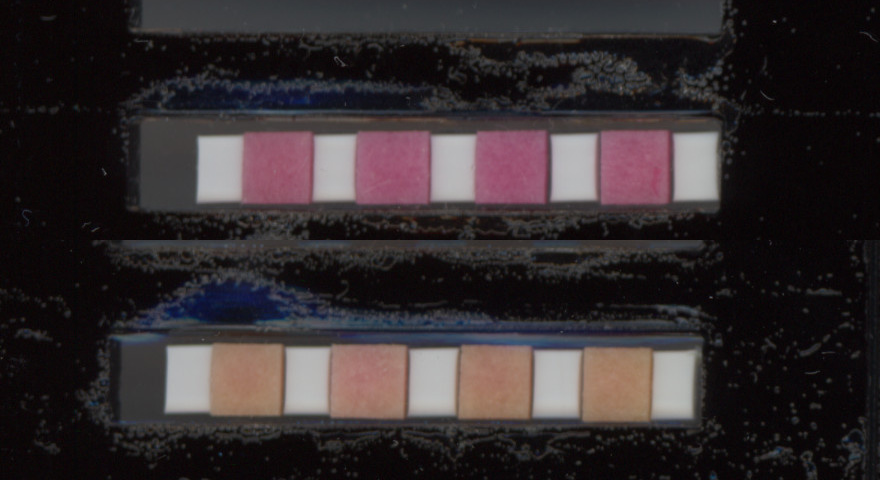

### Timelapse_Image39.jpg

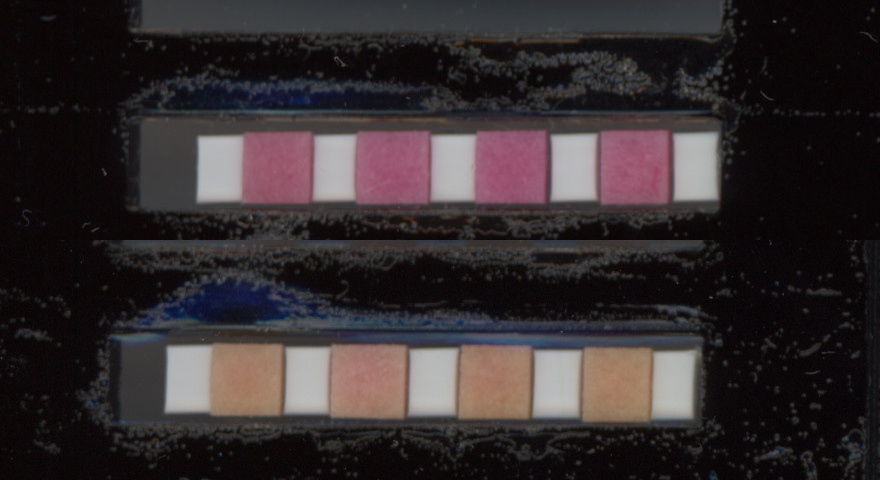

### Timelapse_Image40.jpg

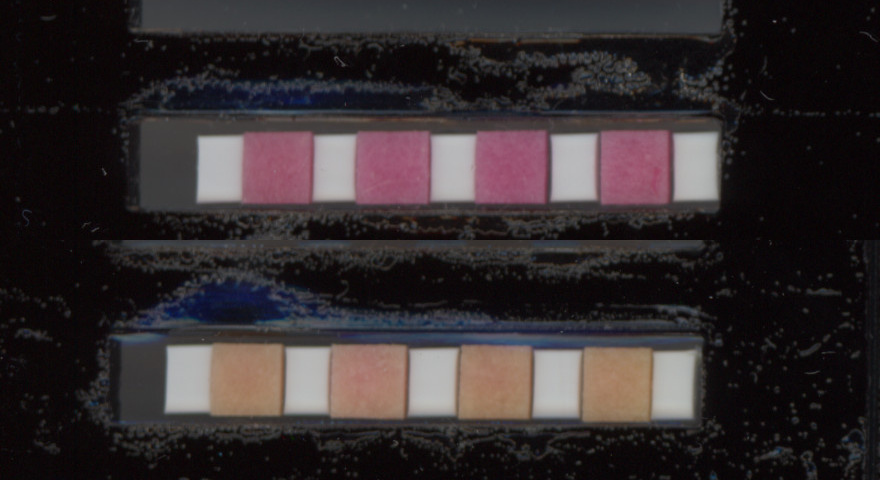

### Timelapse_Image41.jpg

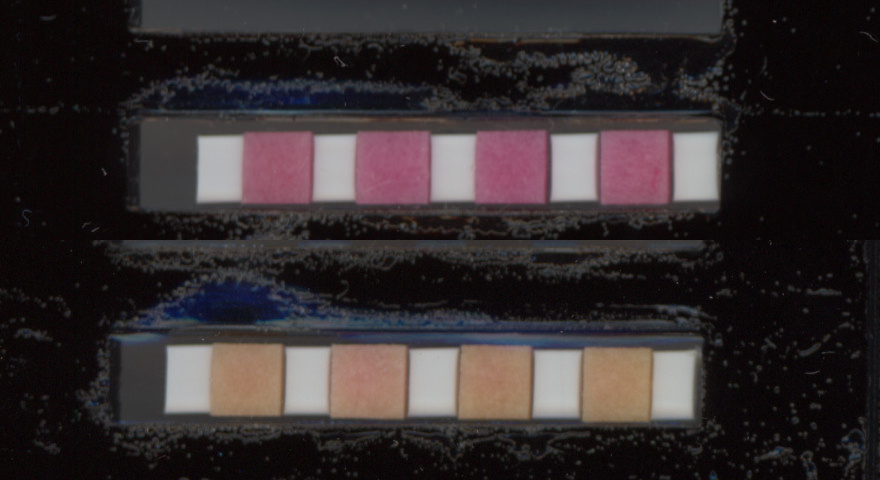

### Timelapse_Image42.jpg

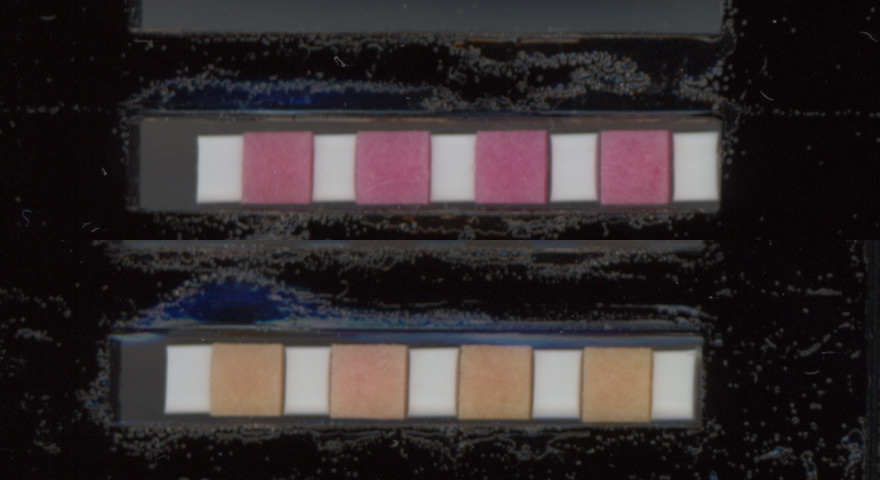

### Timelapse_Image43.jpg

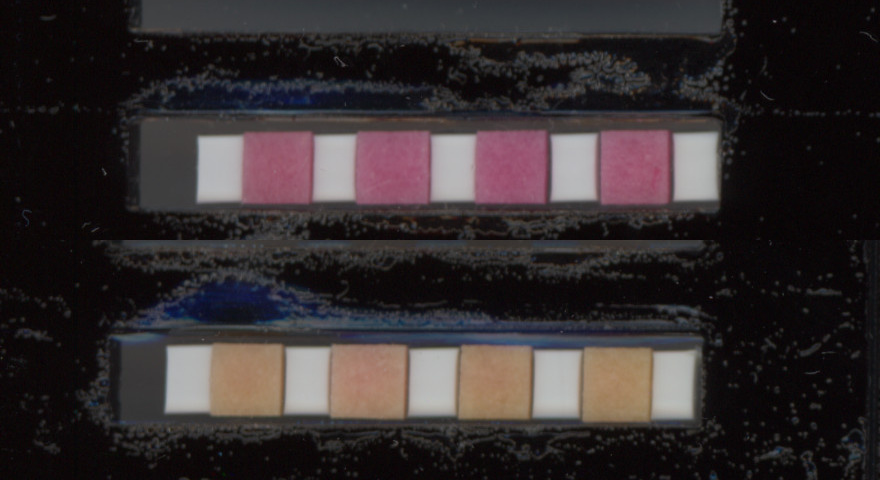

### Timelapse_Image44.jpg

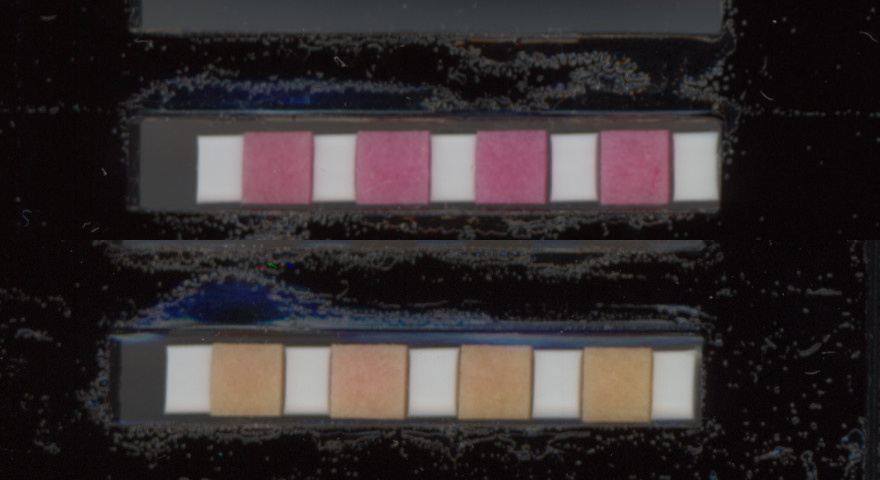

### Timelapse_Image45.jpg

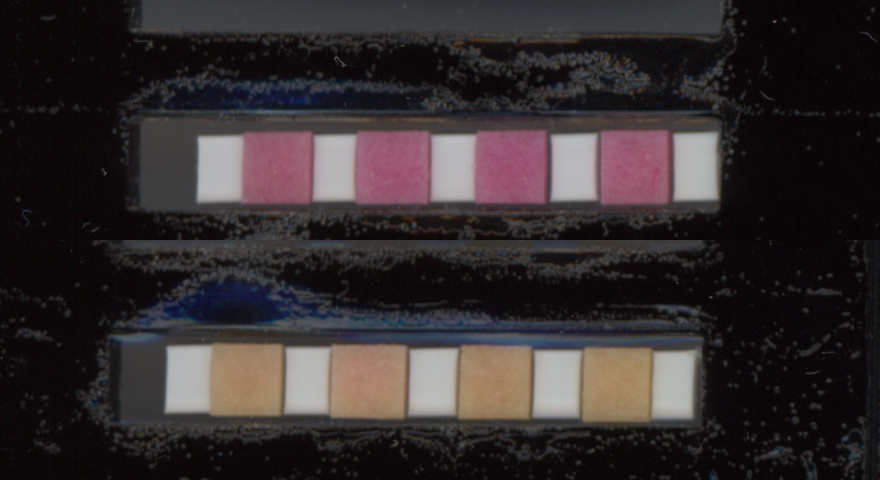

### Timelapse_Image46.jpg

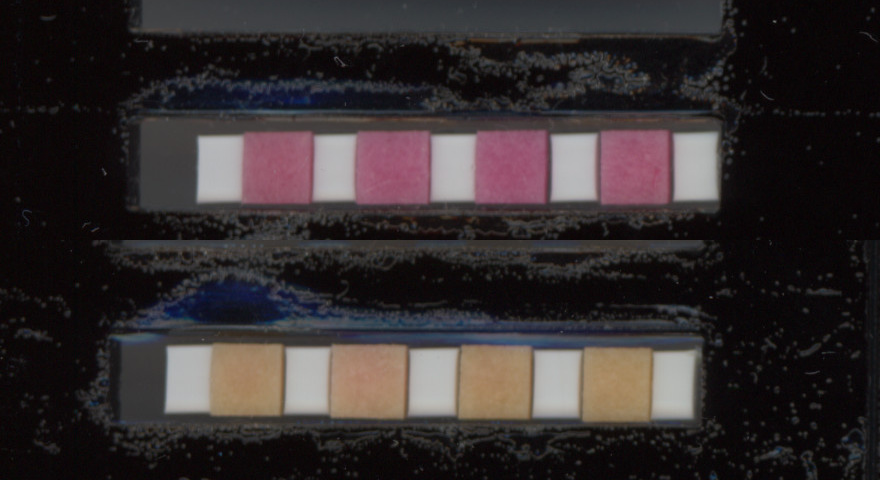

### Timelapse_Image47.jpg

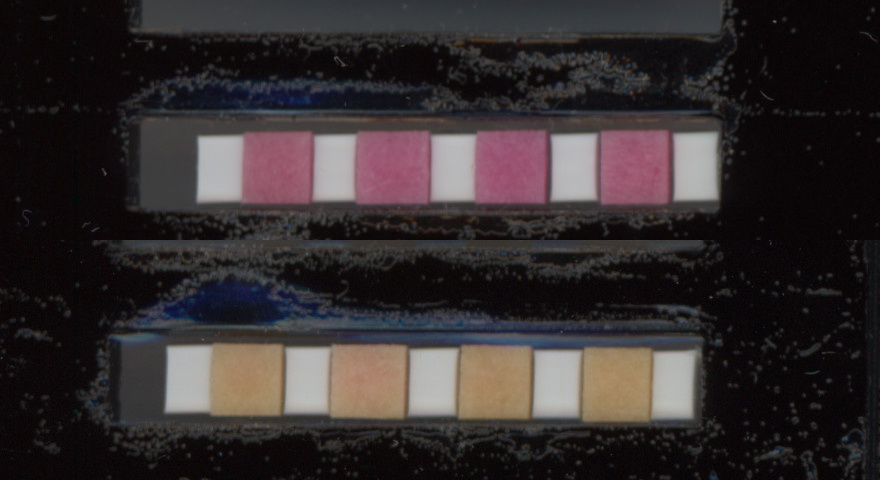

### Timelapse_Image48.jpg

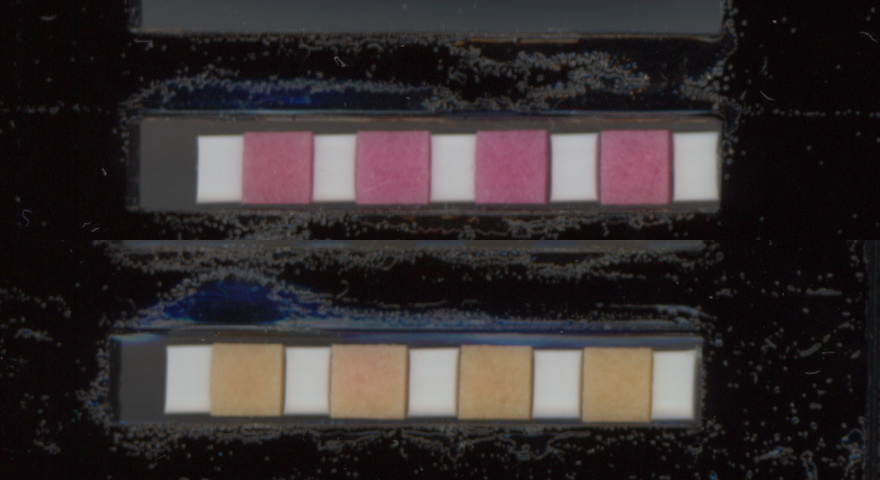

### Timelapse_Image49.jpg

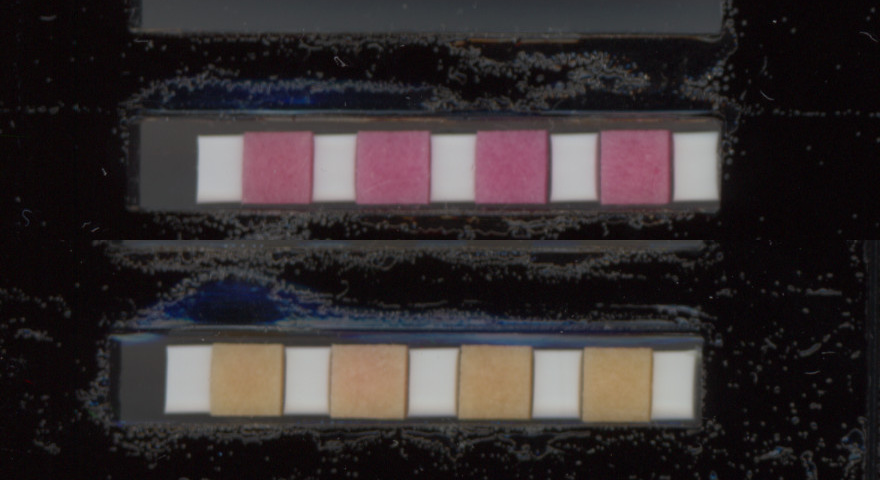

### Timelapse_Image50.jpg

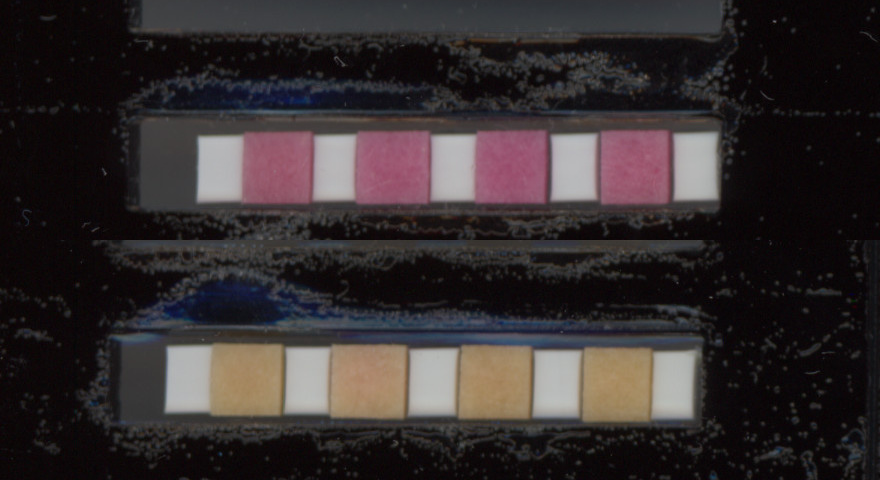

### Timelapse_Image51.jpg

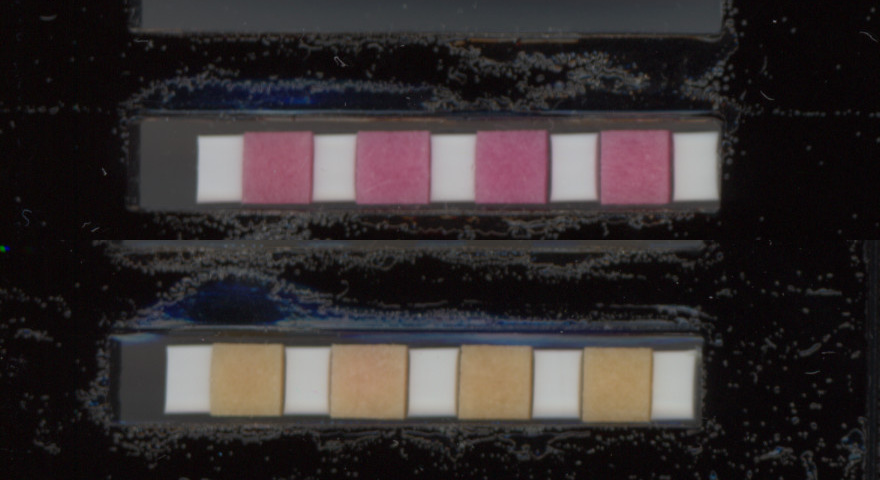

### Timelapse_Image52.jpg

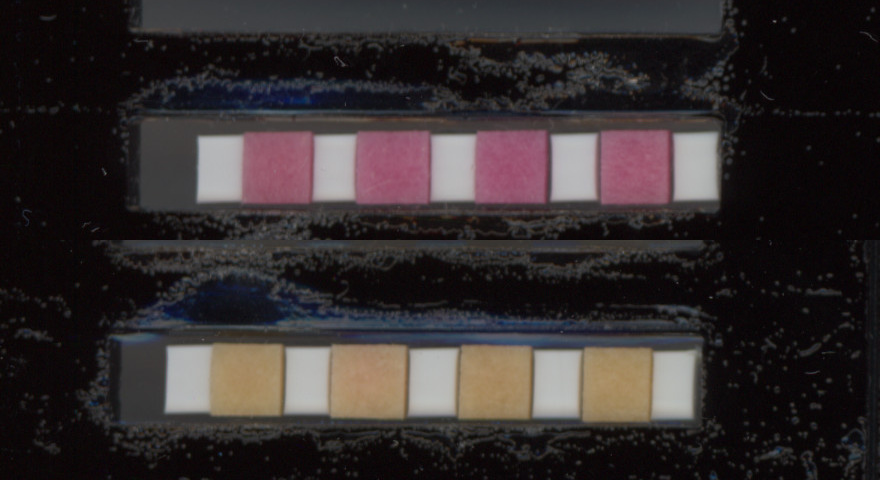

### Timelapse_Image53.jpg

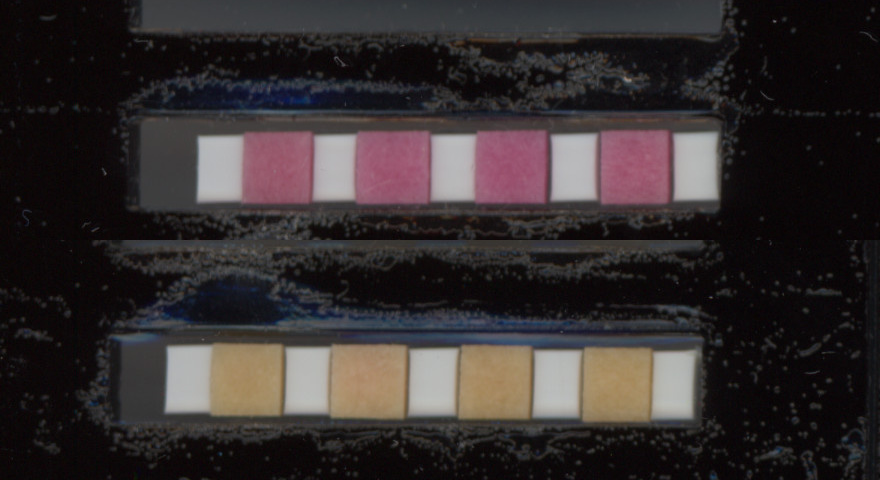

### Timelapse_Image54.jpg

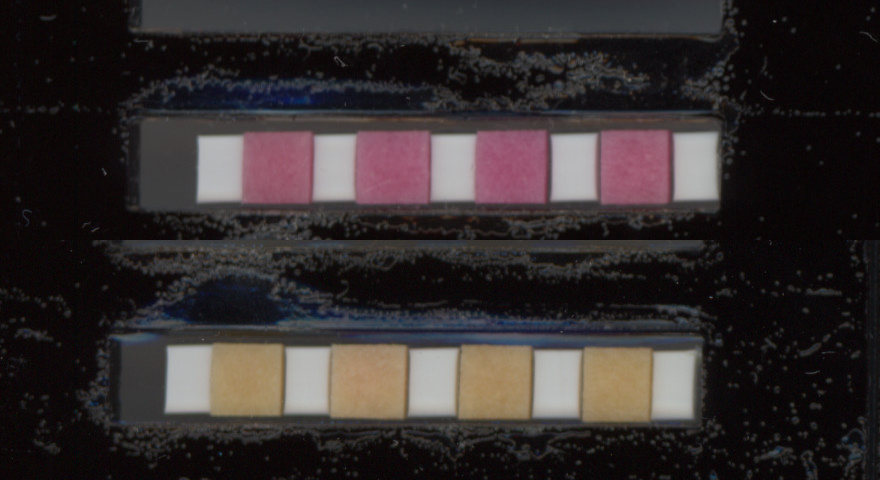

### Timelapse_Image55.jpg

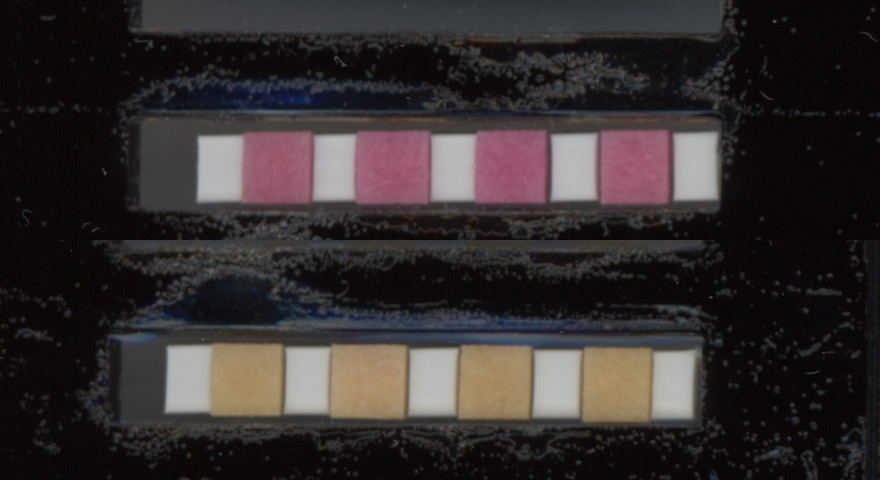

### Timelapse_Image56.jpg

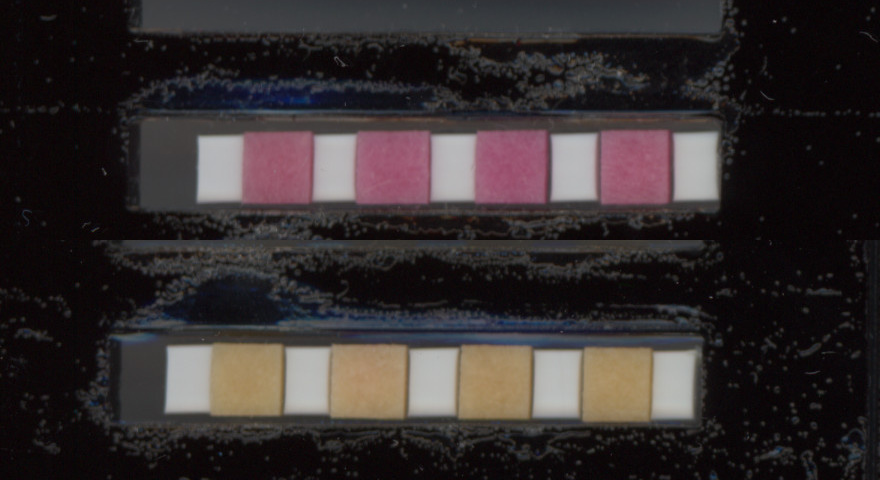

### Timelapse_Image57.jpg

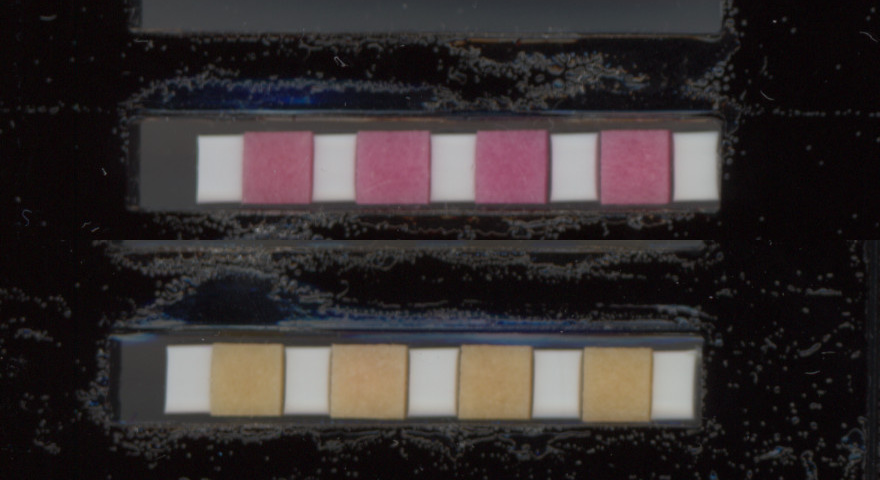

### Timelapse_Image58.jpg

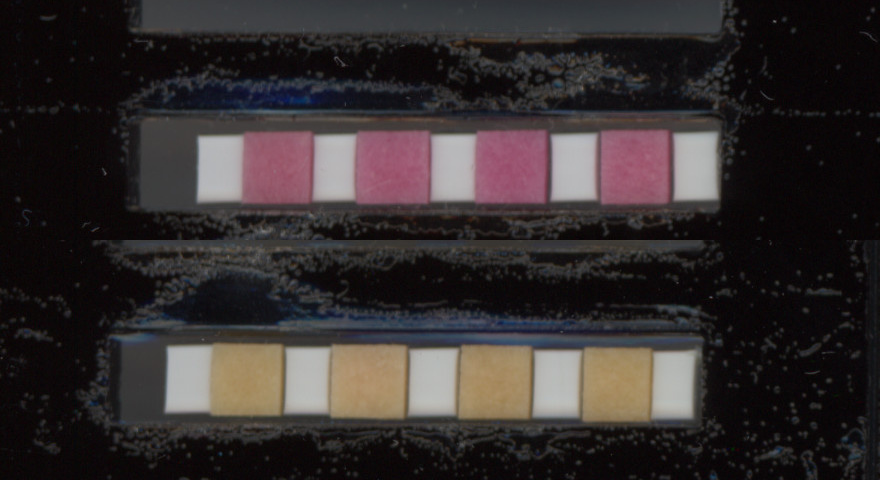

### Timelapse_Image59.jpg

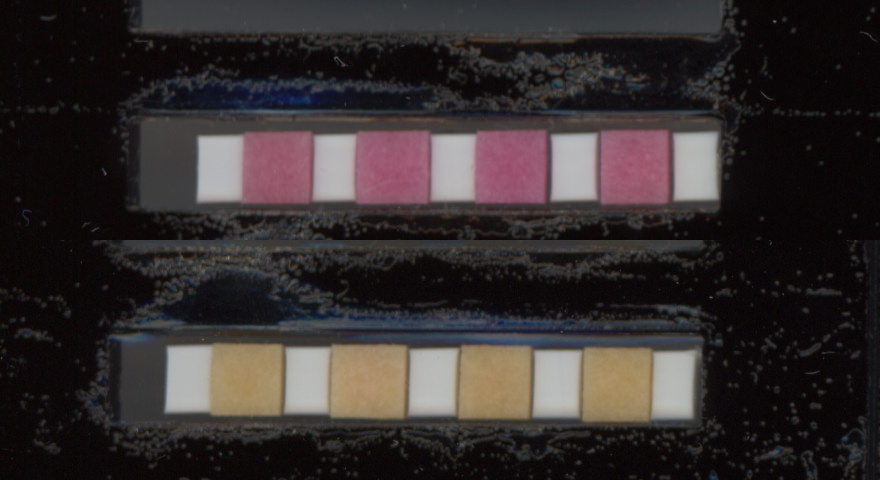

### Timelapse_Image60.jpg

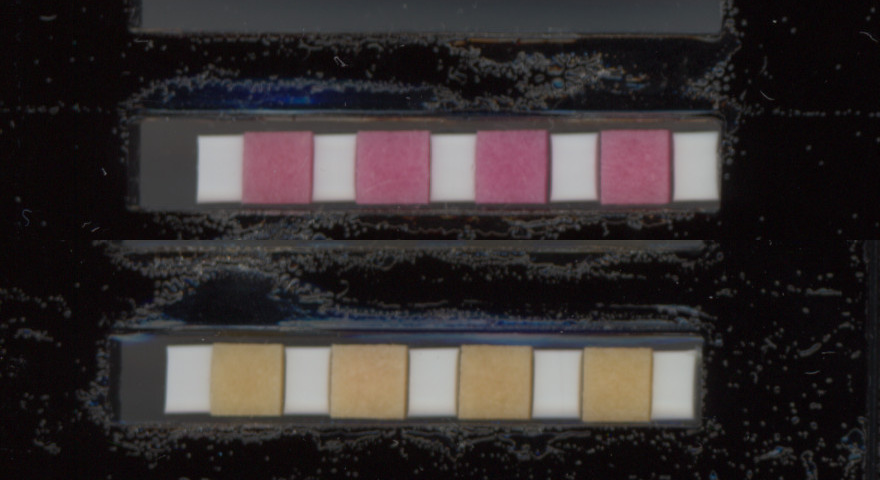

### Timelapse_Image61.jpg

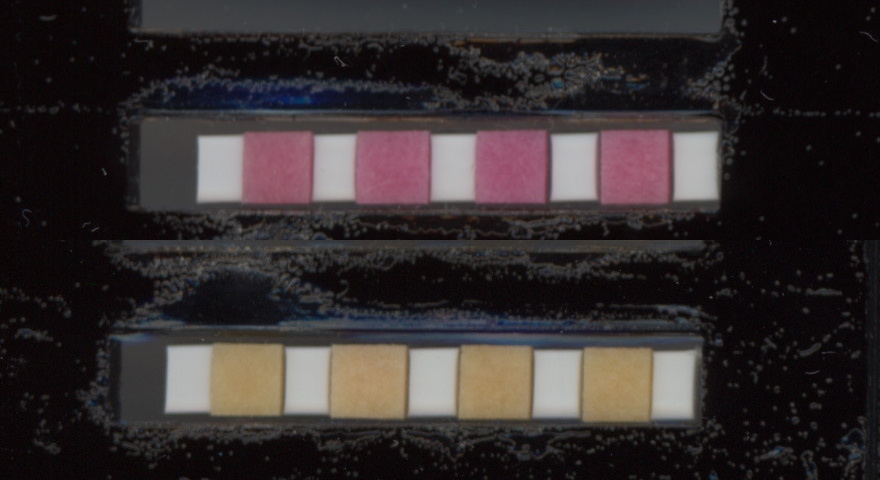

### Timelapse_Image62.jpg

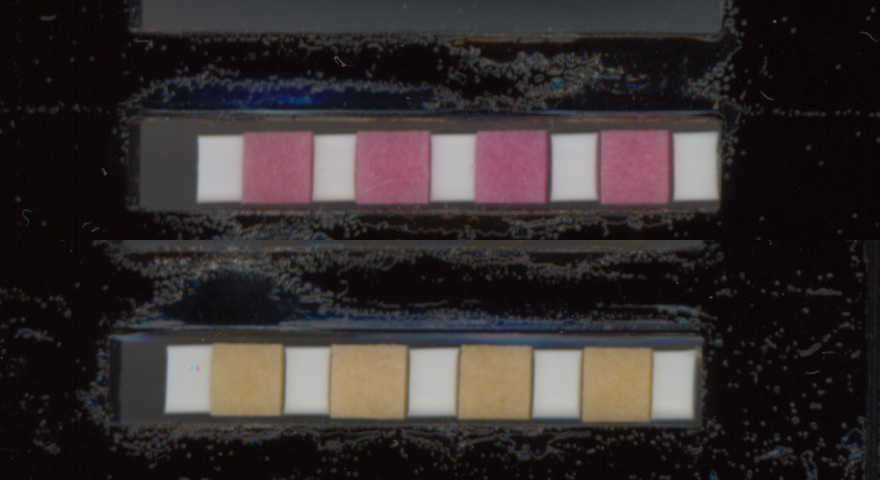

### Timelapse_Image63.jpg

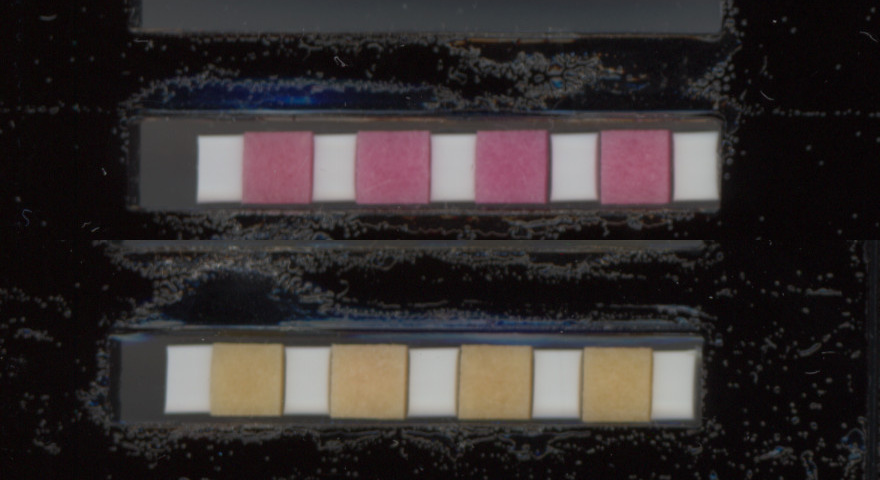

### Timelapse_Image64.jpg

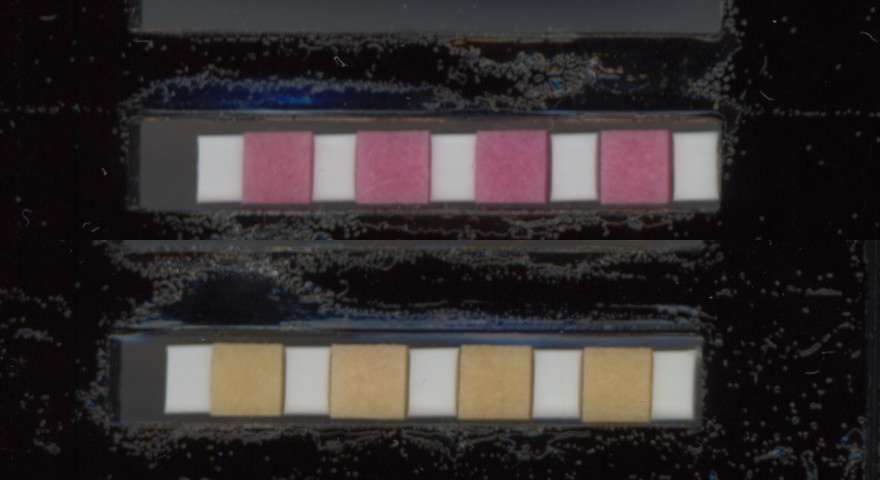

### Timelapse_Image65.jpg

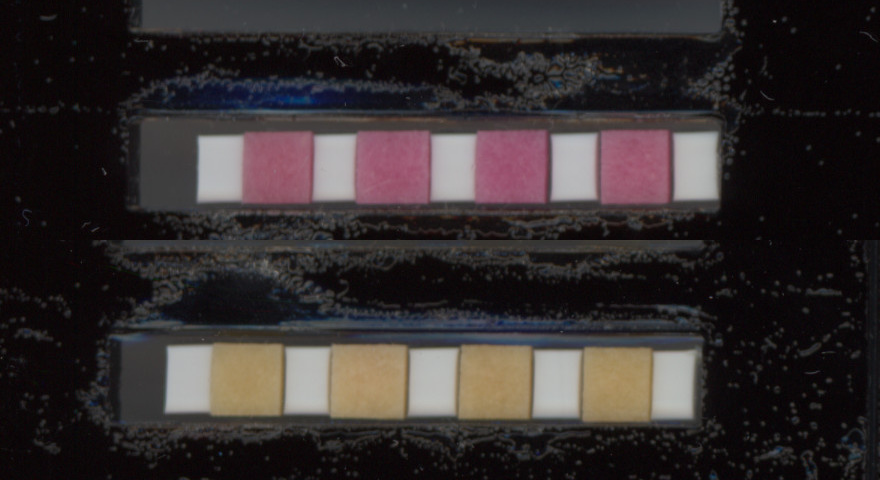

### Timelapse_Image66.jpg

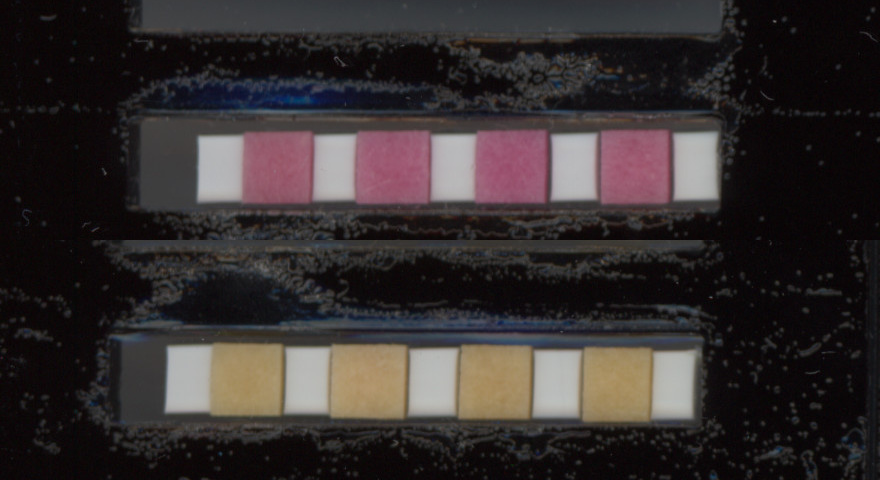

### Timelapse_Image67.jpg

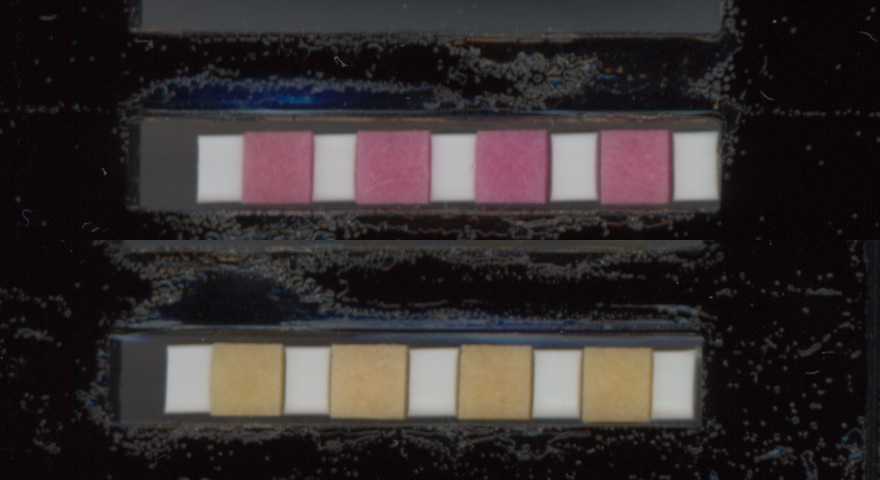

### Timelapse_Image68.jpg

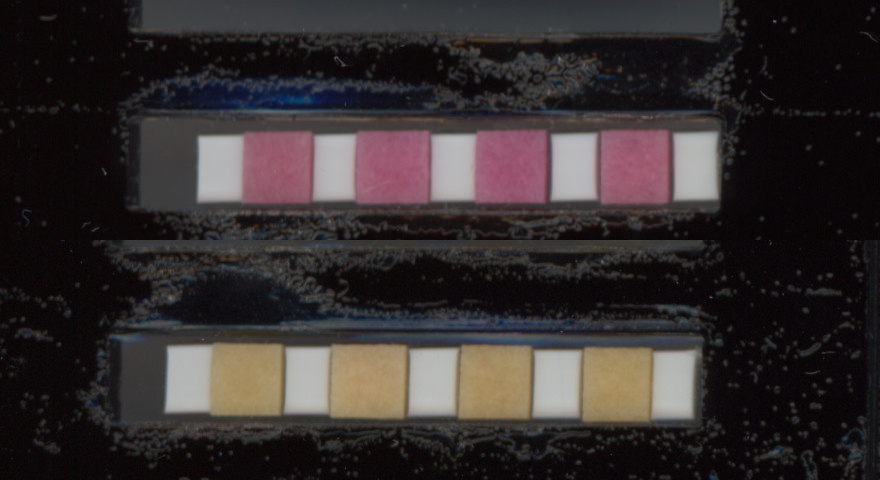

### Timelapse_Image69.jpg

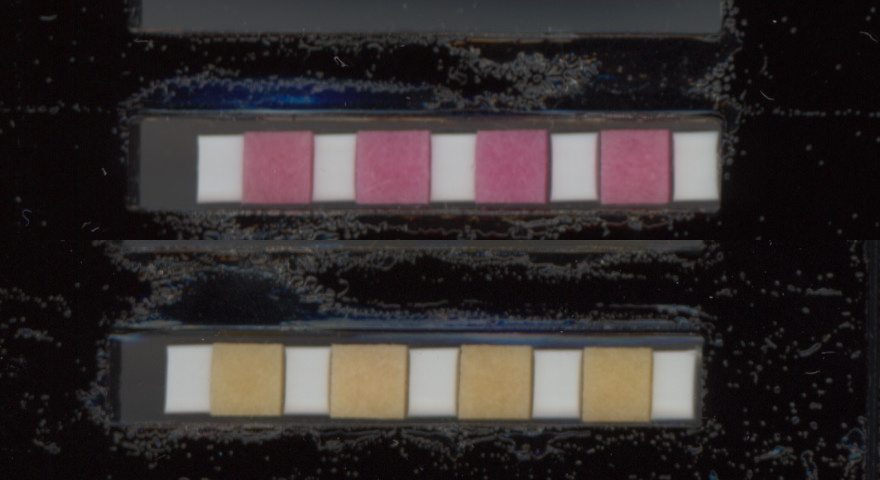

### Timelapse_Image70.jpg

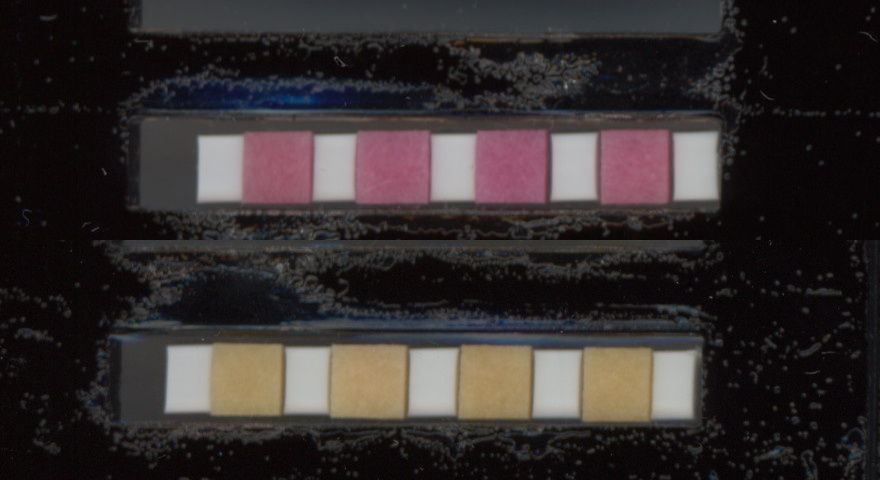

### Timelapse_Image71.jpg

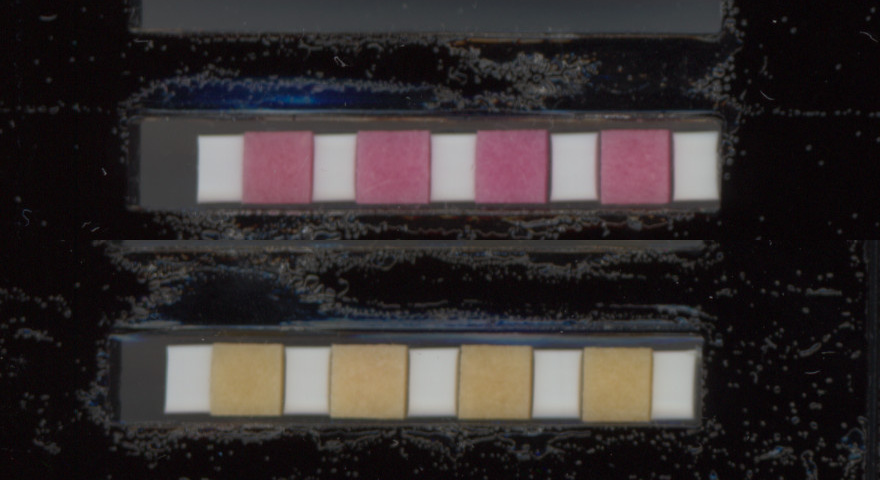

### Timelapse_Image72.jpg

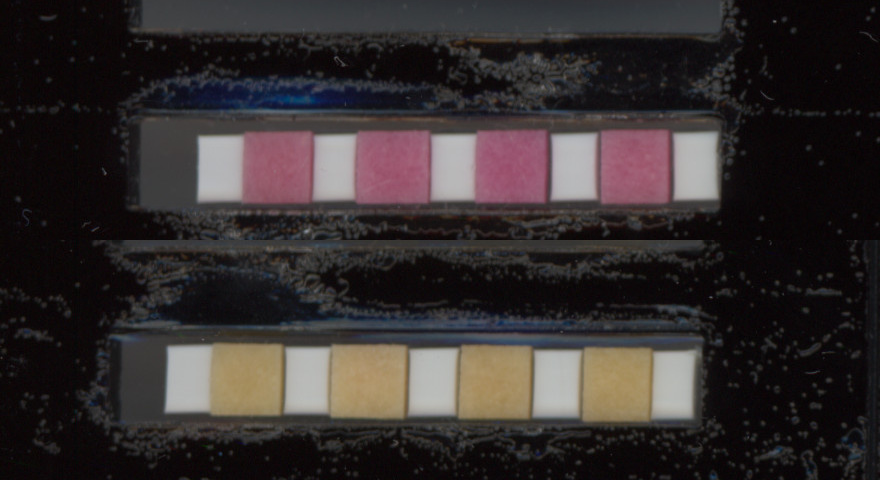

### Timelapse_Image73.jpg

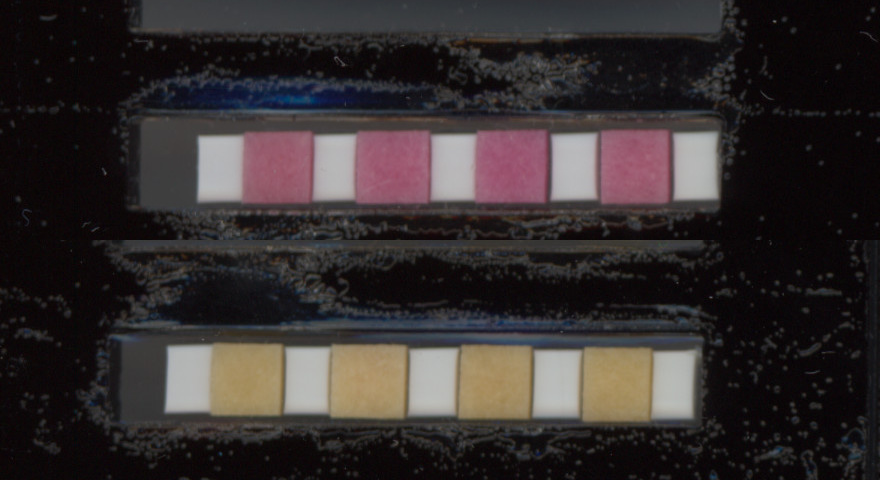

### Timelapse_Image74.jpg

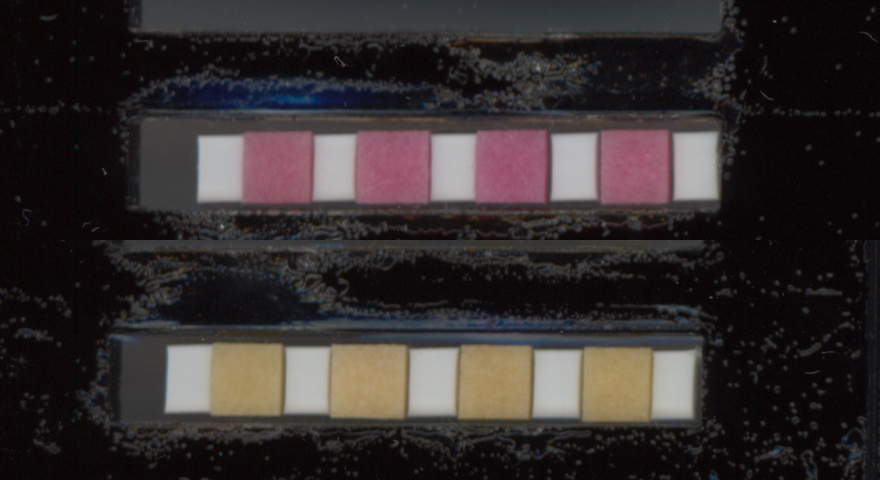

### Timelapse_Image75.jpg

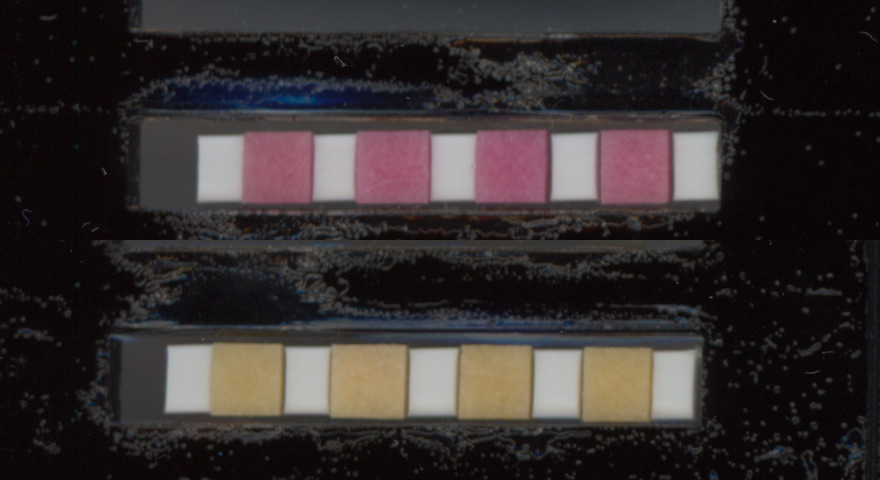

### Timelapse_Image76.jpg

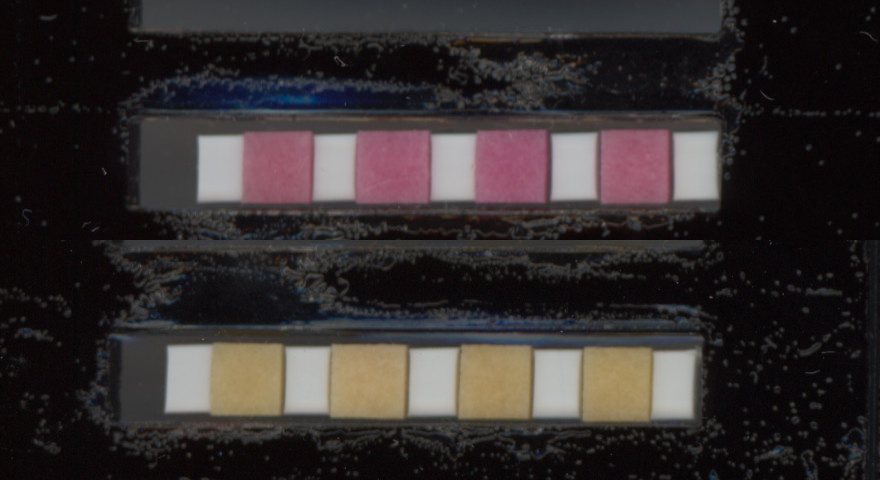

### Timelapse_Image77.jpg

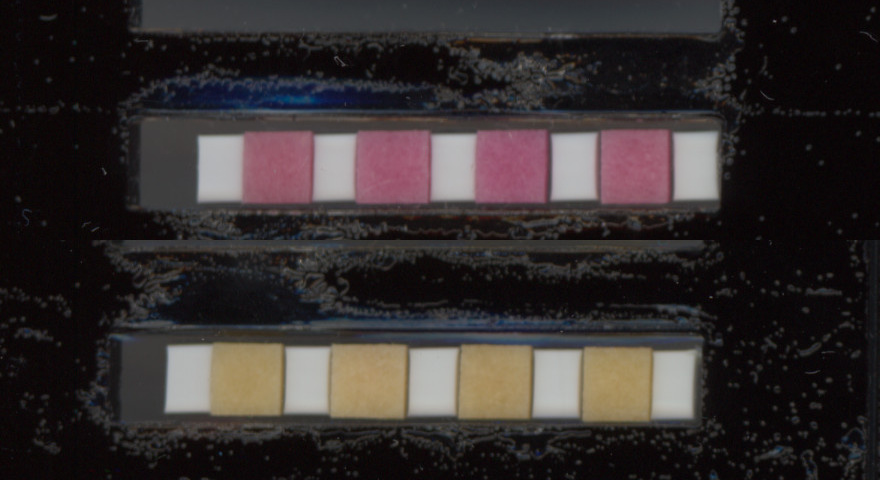

### Timelapse_Image78.jpg

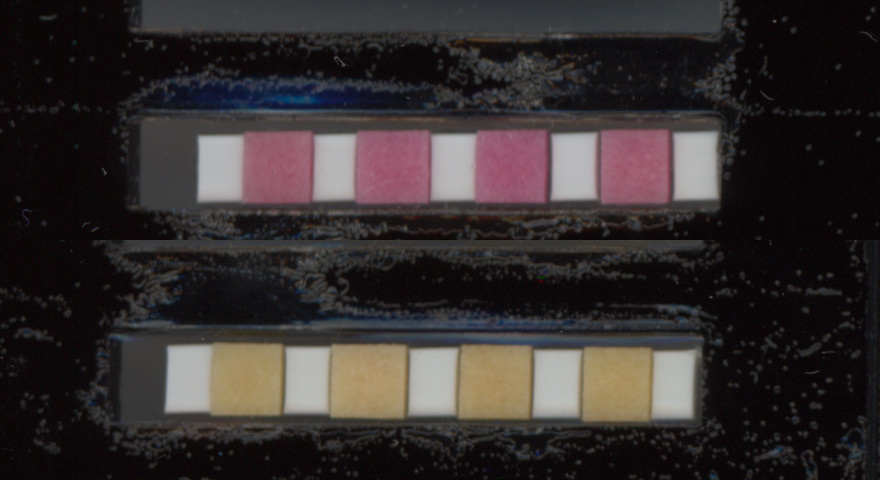

### Timelapse_Image79.jpg

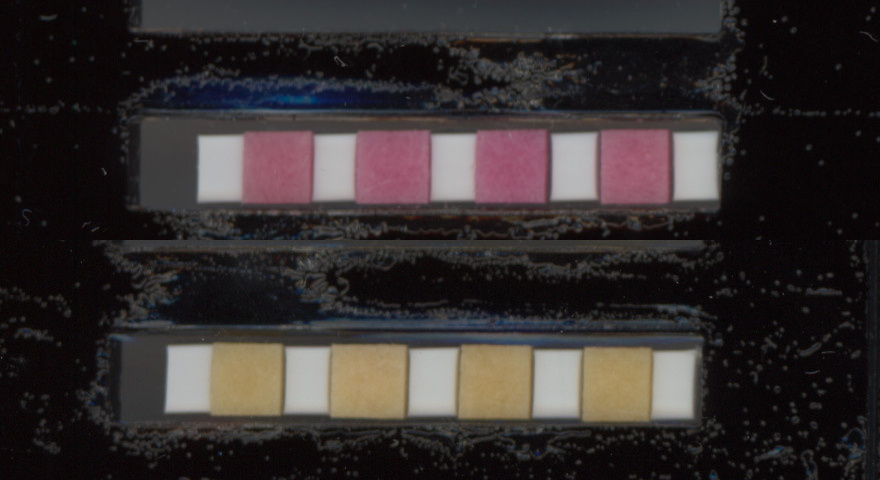

### Timelapse_Image80.jpg

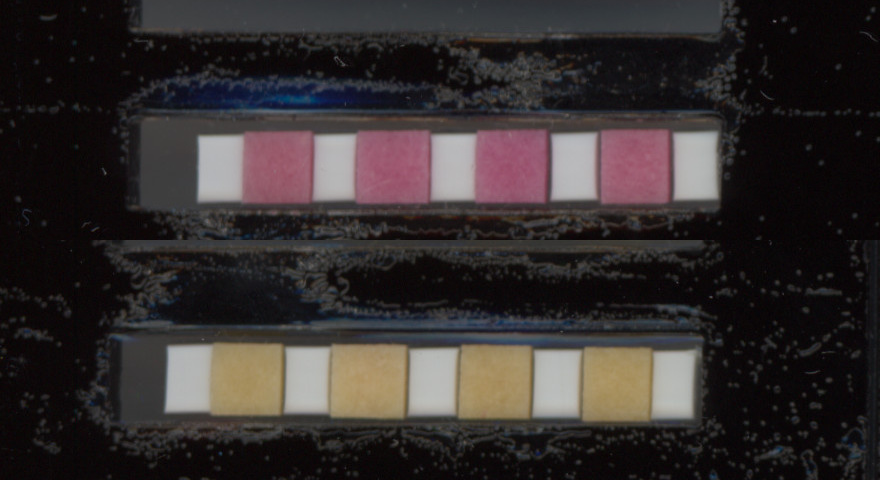

### Timelapse_Image81.jpg

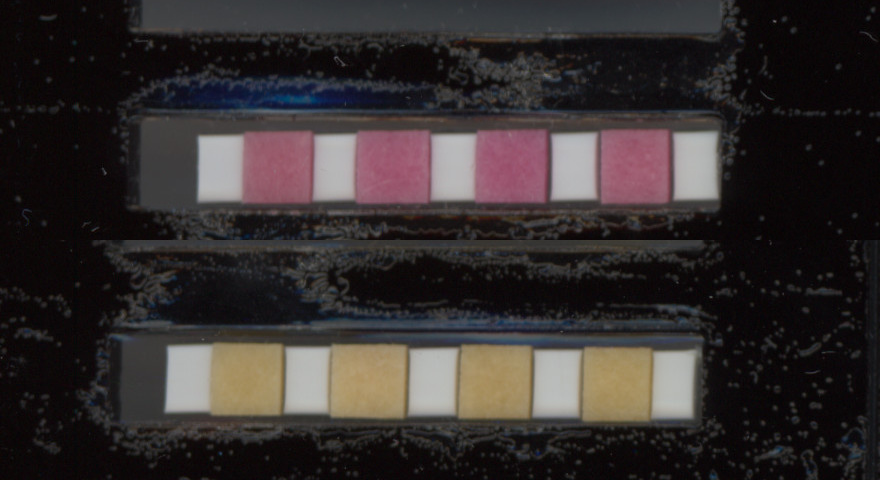

### Timelapse_Image82.jpg

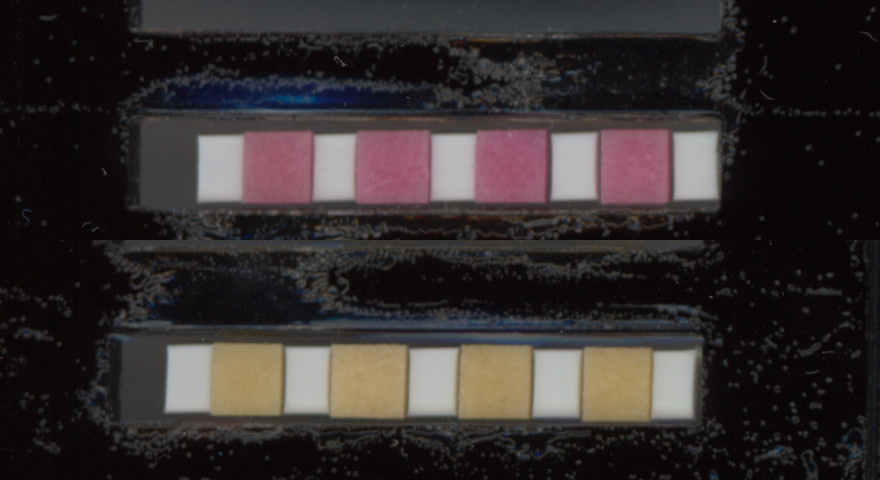

### Timelapse_Image83.jpg

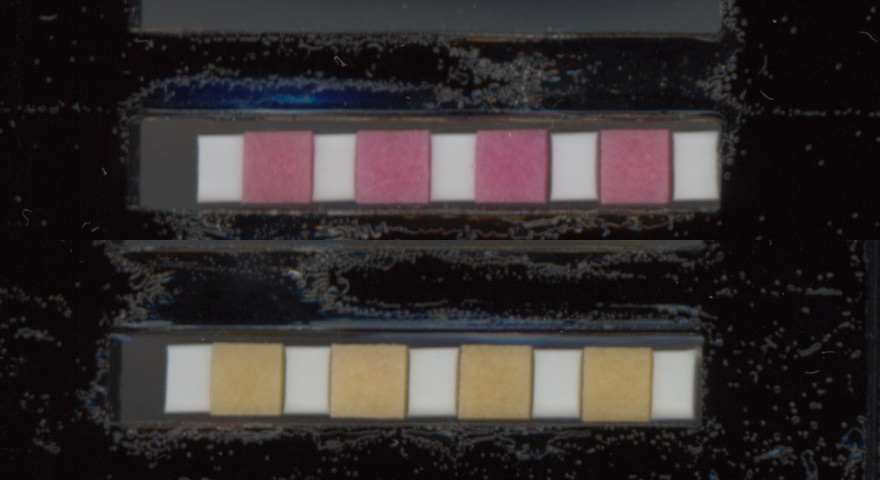

### Timelapse_Image84.jpg

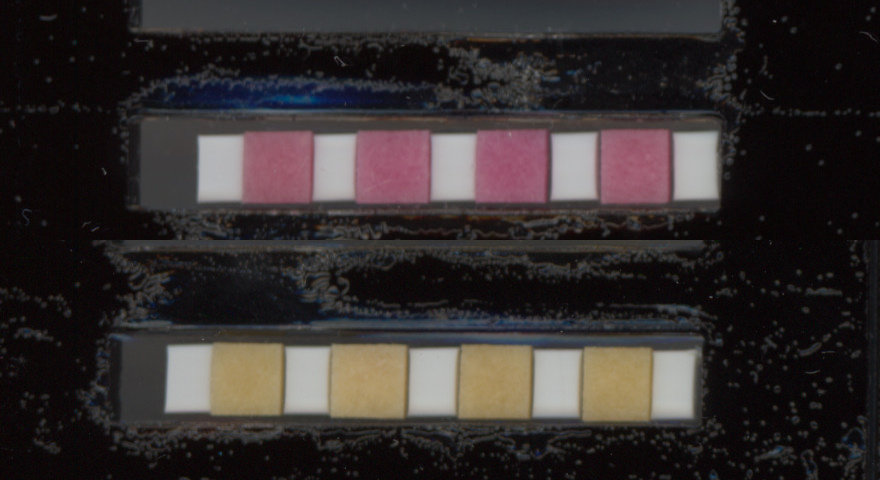

### Timelapse_Image85.jpg

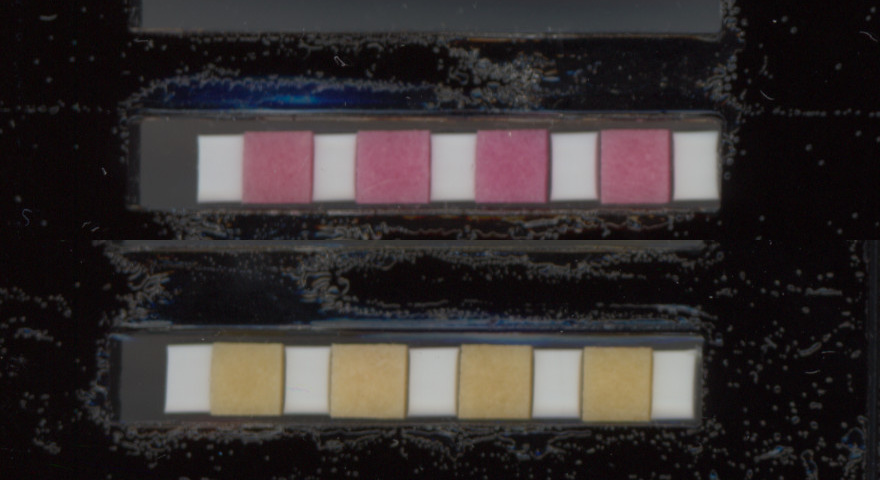

### Timelapse_Image86.jpg

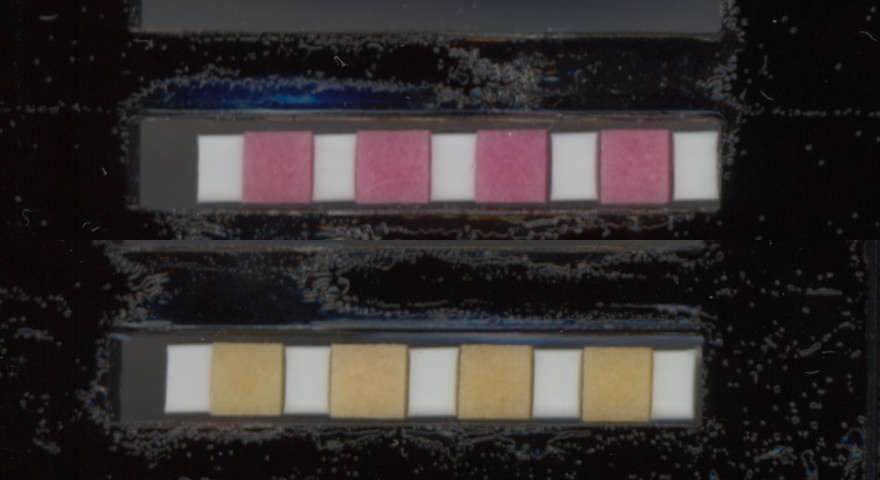

### Timelapse_Image87.jpg

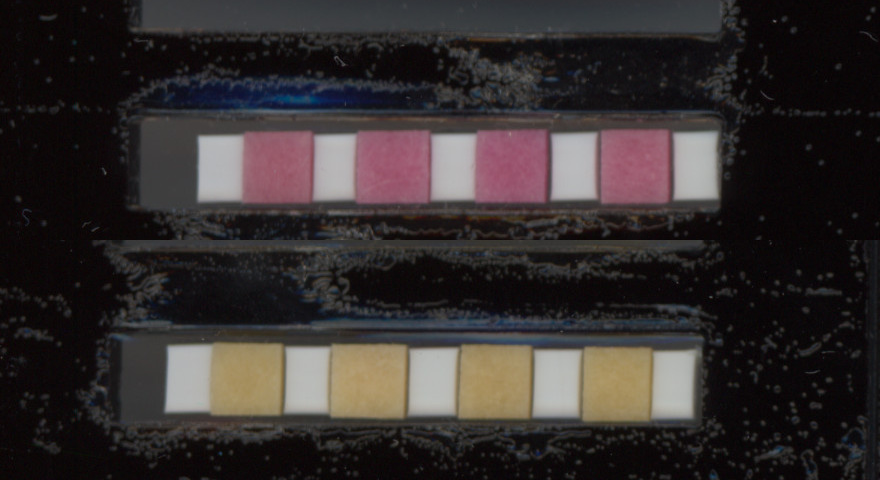

### Timelapse_Image88.jpg

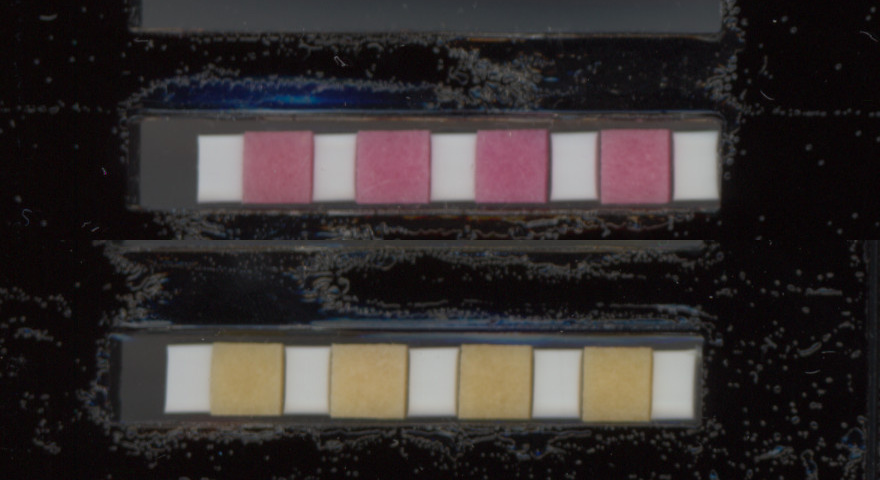
