## Supplementary figures and images for "ThermiQuant™ MegaScan: High-throughput isothermal reactor with quantitative colorimetric readout for paper-based nucleic acid amplification tests"

### AnnotatedImage_with_labels.png

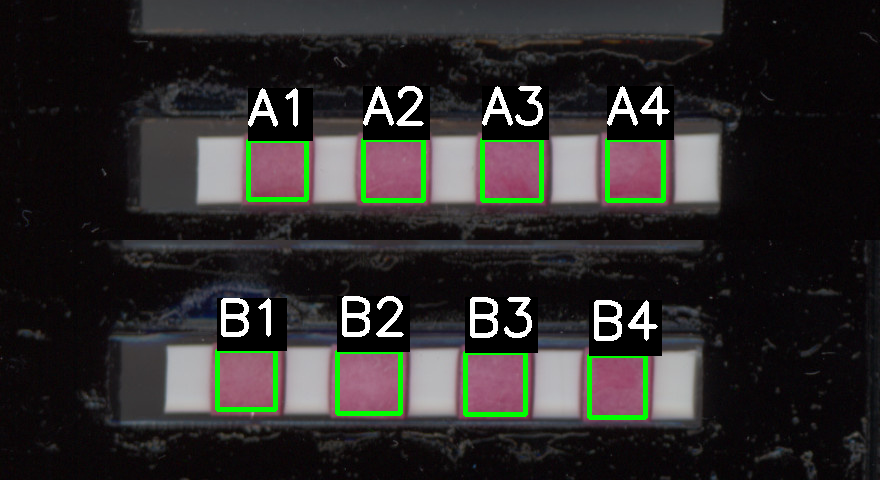

### first_combined_image_with_labels.png

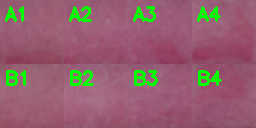

### ProcessedImage_1.png

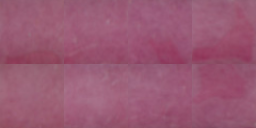

### ProcessedImage_2.png

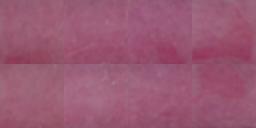

### ProcessedImage_3.png

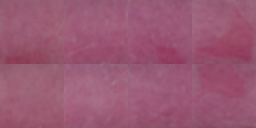

### ProcessedImage_4.png

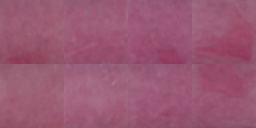

### ProcessedImage_5.png

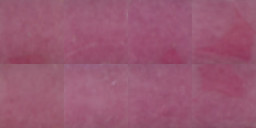

### ProcessedImage_6.png

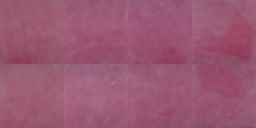

### ProcessedImage_7.png

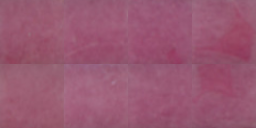

### ProcessedImage_8.png

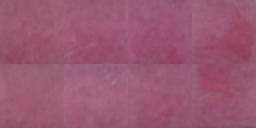

### ProcessedImage_9.png

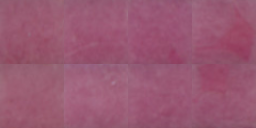

### ProcessedImage_10.png

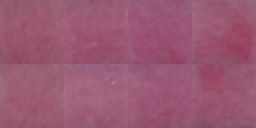

### ProcessedImage_11.png

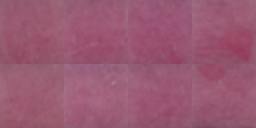

### ProcessedImage_12.png

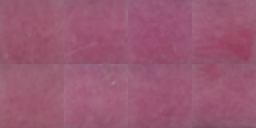

### ProcessedImage_13.png

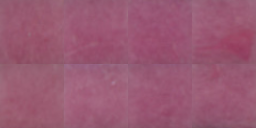

### ProcessedImage_14.png

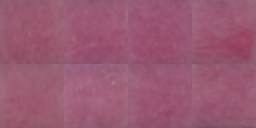

### ProcessedImage_15.png

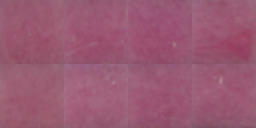

### ProcessedImage_16.png

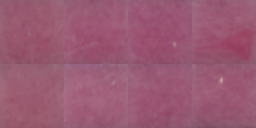

### ProcessedImage_17.png

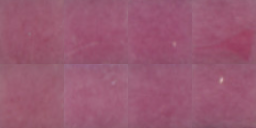

### ProcessedImage_18.png

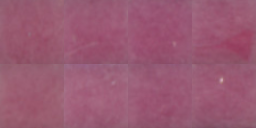

### ProcessedImage_19.png

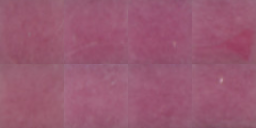

### ProcessedImage_20.png

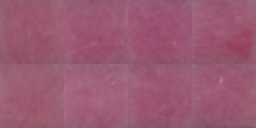

### ProcessedImage_21.png

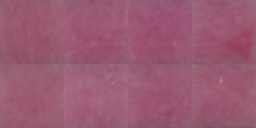

### ProcessedImage_22.png

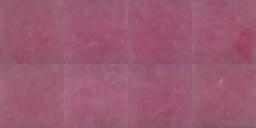

### ProcessedImage_23.png

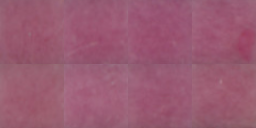

### ProcessedImage_24.png

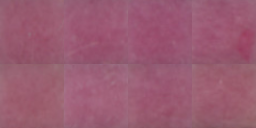

### ProcessedImage_25.png

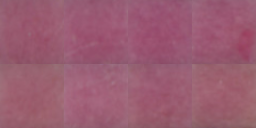

### ProcessedImage_26.png

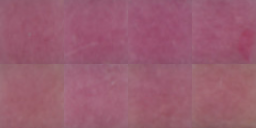

### ProcessedImage_27.png

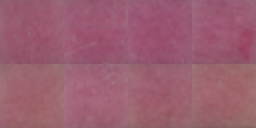

### ProcessedImage_28.png

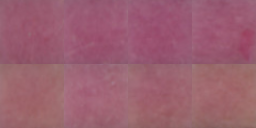

### ProcessedImage_29.png

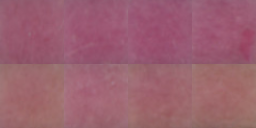

### ProcessedImage_30.png

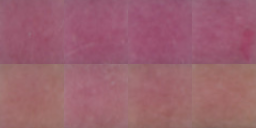

### ProcessedImage_31.png

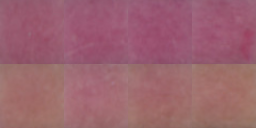

### ProcessedImage_32.png

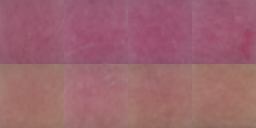

### ProcessedImage_33.png

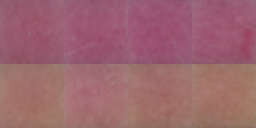

### ProcessedImage_34.png

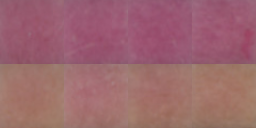

### ProcessedImage_35.png

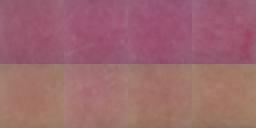

### ProcessedImage_36.png

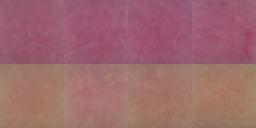

### ProcessedImage_37.png

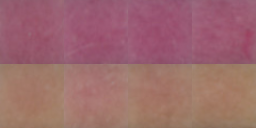

### ProcessedImage_38.png

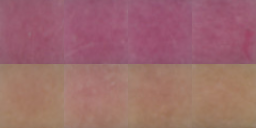

### ProcessedImage_39.png

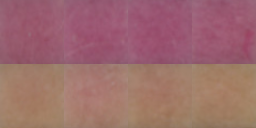

### ProcessedImage_40.png

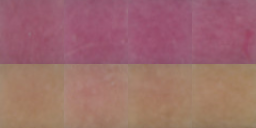

### ProcessedImage_41.png

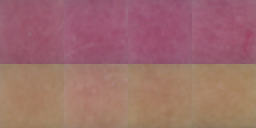

### ProcessedImage_42.png

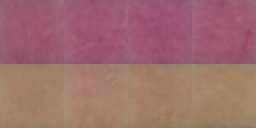

### ProcessedImage_43.png

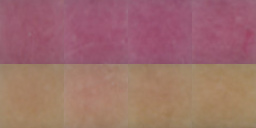

### ProcessedImage_44.png

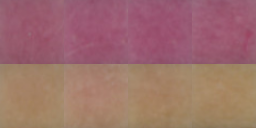

### ProcessedImage_45.png

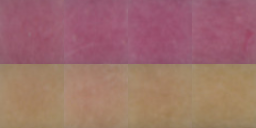

### ProcessedImage_46.png

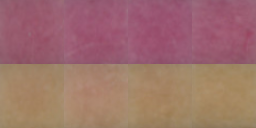

### ProcessedImage_47.png

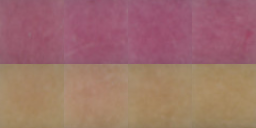

### ProcessedImage_48.png

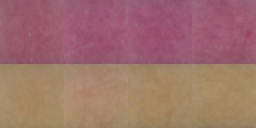

### ProcessedImage_49.png

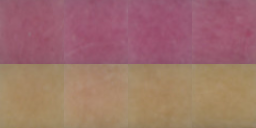

### ProcessedImage_50.png

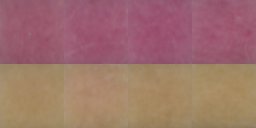

### ProcessedImage_51.png

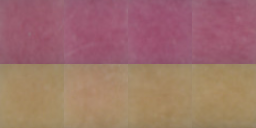

### ProcessedImage_52.png

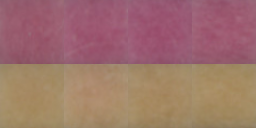

### ProcessedImage_53.png

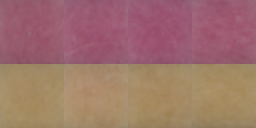

### ProcessedImage_54.png

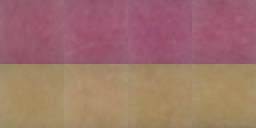

### ProcessedImage_55.png

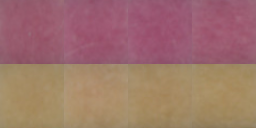

### ProcessedImage_56.png

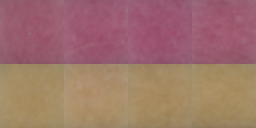

### ProcessedImage_57.png

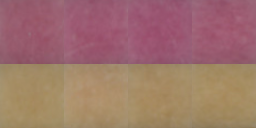

### ProcessedImage_58.png

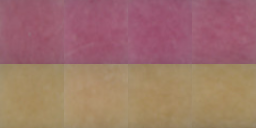

### ProcessedImage_59.png

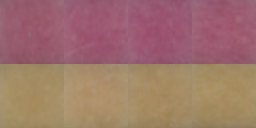

### ProcessedImage_60.png

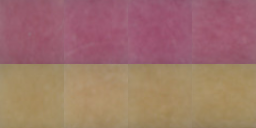

### ProcessedImage_61.png

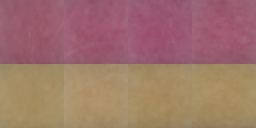

### ProcessedImage_62.png

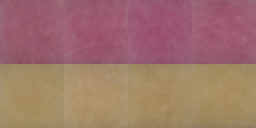

### ProcessedImage_63.png

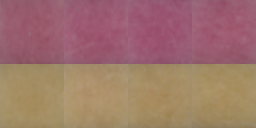

### ProcessedImage_64.png

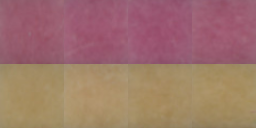

### ProcessedImage_65.png

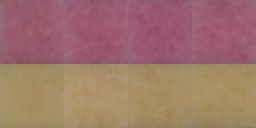

### ProcessedImage_66.png

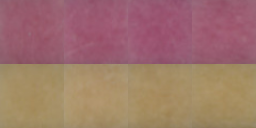

### ProcessedImage_67.png

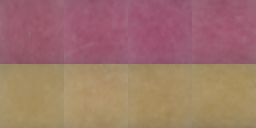

### ProcessedImage_68.png

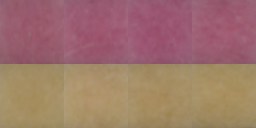

### ProcessedImage_69.png

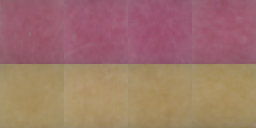

### ProcessedImage_70.png

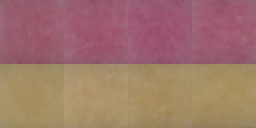

### ProcessedImage_71.png

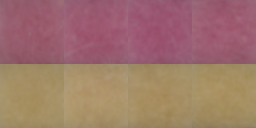

### ProcessedImage_72.png

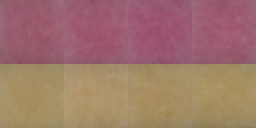

### ProcessedImage_73.png

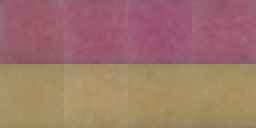

### ProcessedImage_74.png

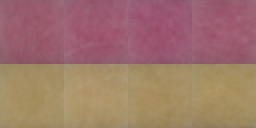

### ProcessedImage_75.png

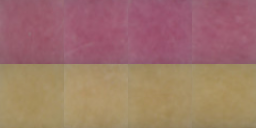

### ProcessedImage_76.png

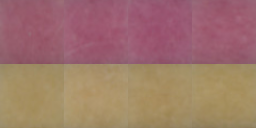

### ProcessedImage_77.png

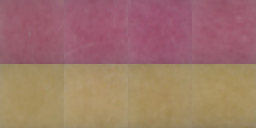

### ProcessedImage_78.png

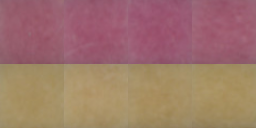

### ProcessedImage_79.png

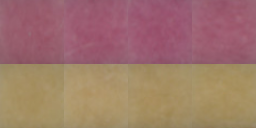

### ProcessedImage_80.png

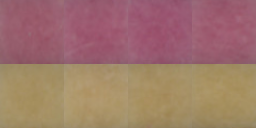

### ProcessedImage_81.png

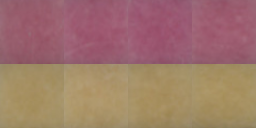

### ProcessedImage_82.png

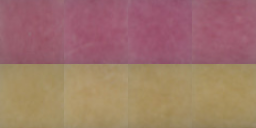

### ProcessedImage_83.png

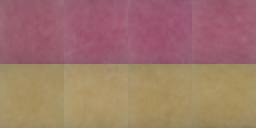

### ProcessedImage_84.png

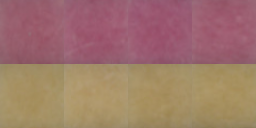

### ProcessedImage_85.png

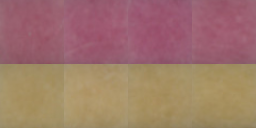

### ProcessedImage_86.png

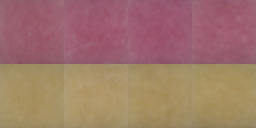

### ProcessedImage_87.png

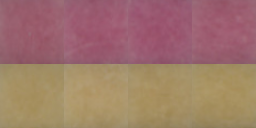

### ProcessedImage_88.png

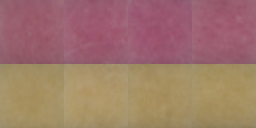

### Timelapse_Image01.jpg

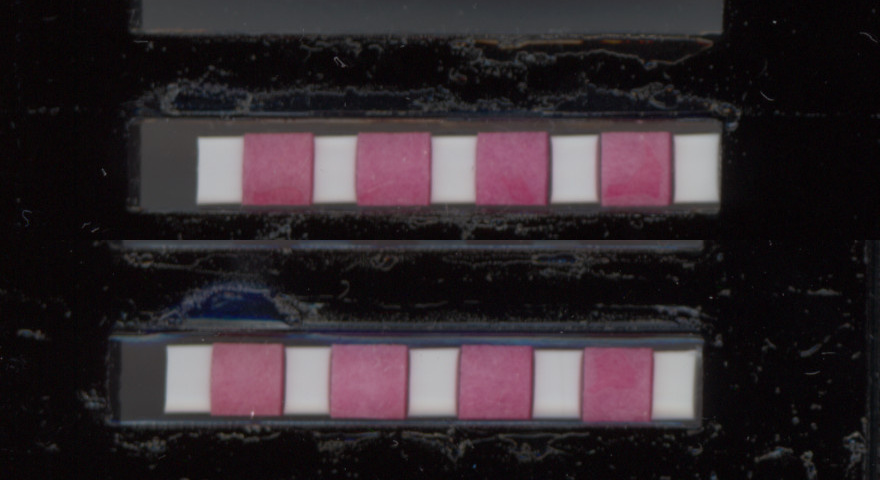

### Timelapse_Image02.jpg

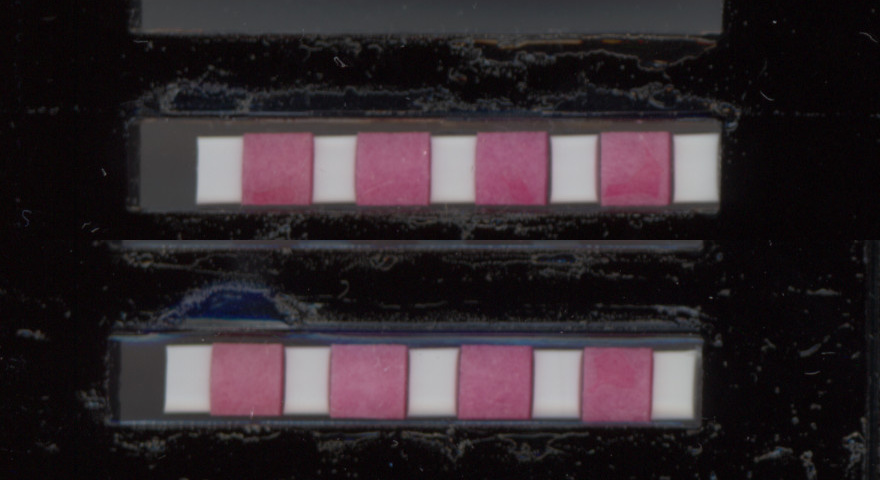

### Timelapse_Image03.jpg

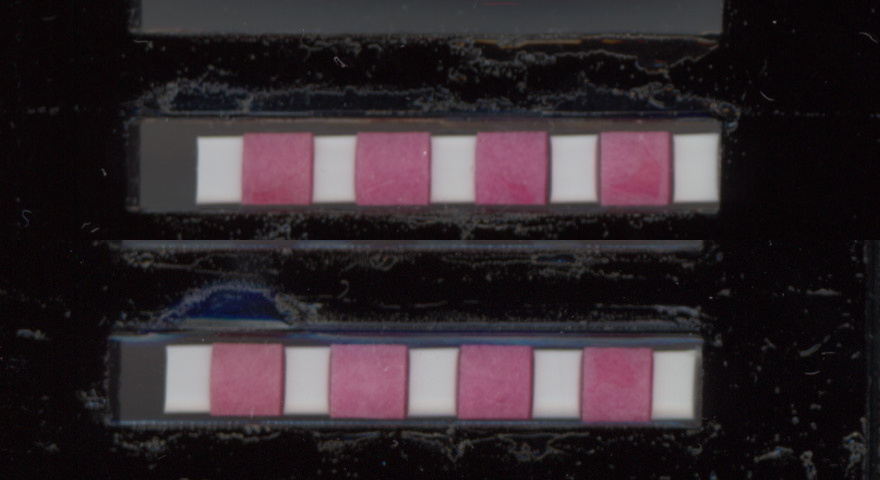

### Timelapse_Image04.jpg

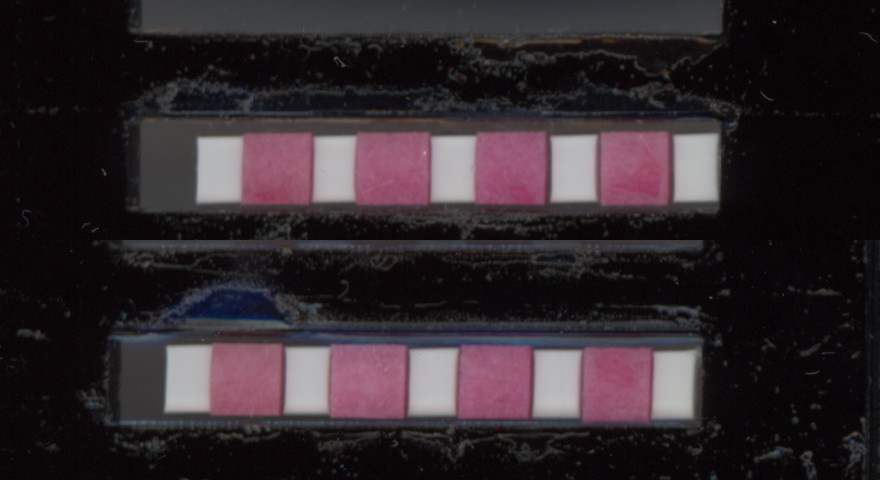

### Timelapse_Image05.jpg

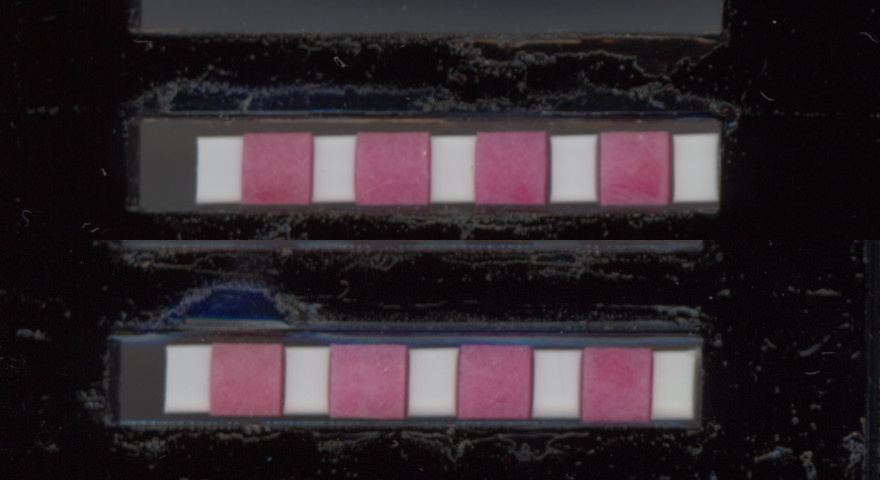

### Timelapse_Image06.jpg

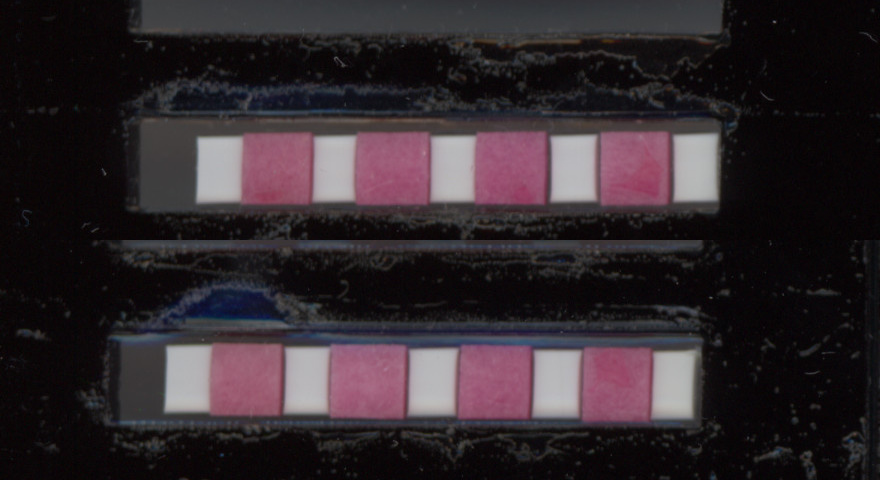

### Timelapse_Image07.jpg

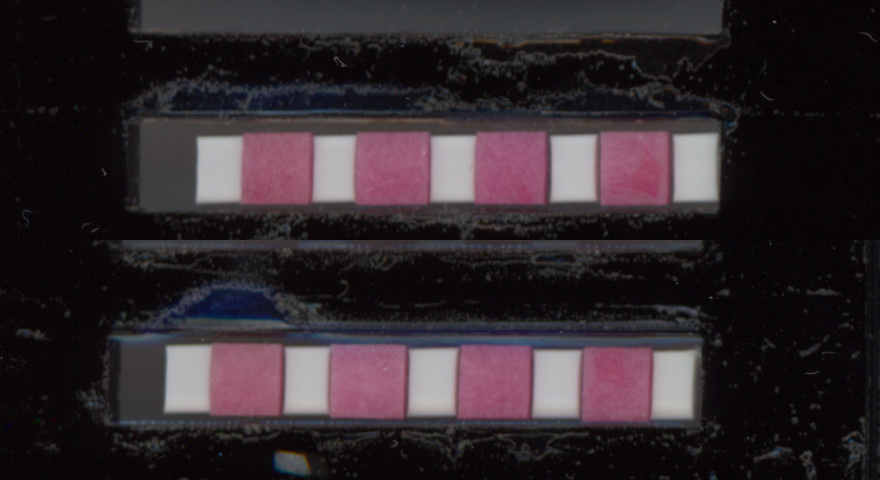

### Timelapse_Image08.jpg

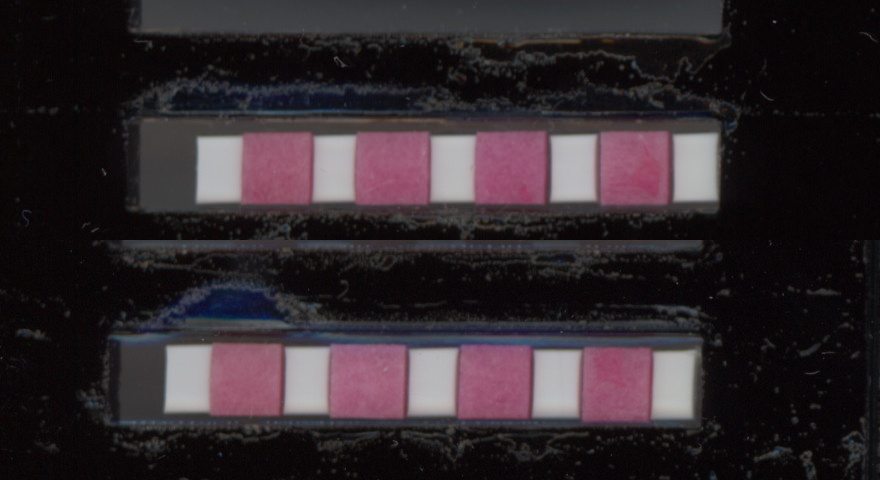

### Timelapse_Image09.jpg

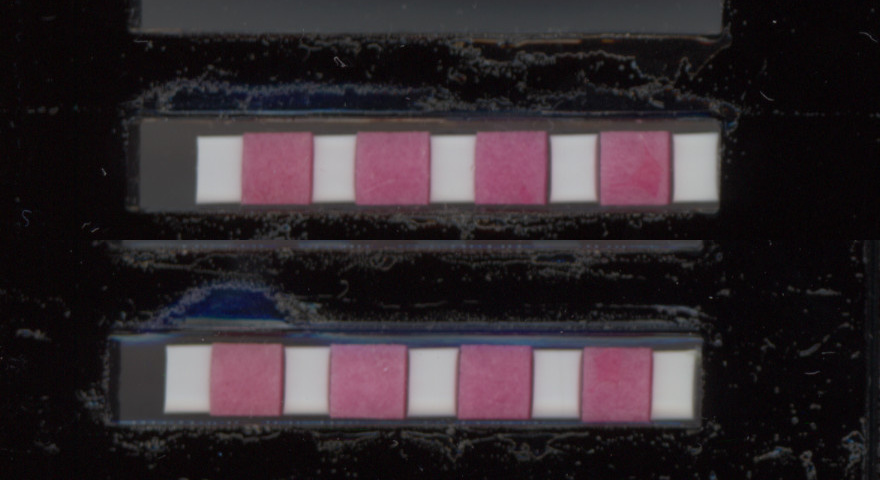

### Timelapse_Image10.jpg

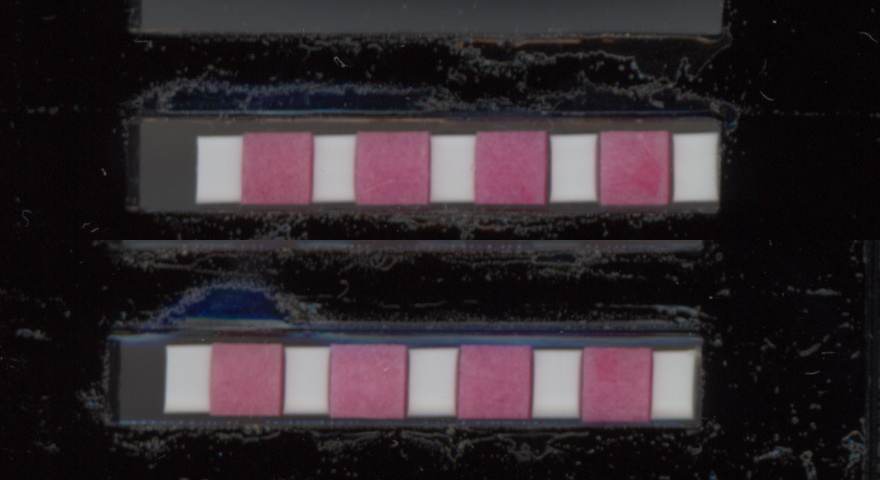
